# Estimating temporally variable selection intensity from ancient DNA data II

**DOI:** 10.1101/2023.07.10.548348

**Authors:** Wenyang Lyu, Xiaoyang Dai, Mark Beaumont, Feng Yu, Zhangyi He

## Abstract

Recent technological innovations, such as next generation sequencing and DNA hybridisation enrichment, have made it possible to recover DNA information from historical and archaeological biological materials, which has motivated the development of various statistical approaches for inferring selection from allele frequency time series data. Recently, He et al. (2023a,b) introduced methods that can utilise ancient DNA (aDNA) data in the form of genotype likelihoods, therefore enabling the modelling of sample uncertainty arising from DNA molecule damage and fragmentation. However, their performance suffers from the underlying dependency on the allele age. Here we introduce a novel particle marginal Metropolis-Hastings within Gibbs framework for Bayesian inference of time-varying selection from aDNA data in the form of genotype likelihoods. To circumvent the performance issue encountered in He et al. (2023a,b), we devise a novel numerical scheme for backward-in-time simulation of the Wright-Fisher diffusion and mix forward- and backward-in-time simulations in the particle filter for likelihood computation. Our framework also enables us to reconstruct the underlying population allele frequency trajectories, integrate temporal information in genotype likelihood calculations and test hypotheses on the drivers of past selection events. We conduct extensive evaluations through simulations and show its utility with an application to aDNA data from pigmentation loci in ancient horses.

## 1. Introduction

Recent advances in technologies for extracting and sequencing DNA molecules from historical and archaeological specimens have resulted in a rapid increase in both the quality and quantity of ancient DNA (aDNA) data, including those from humans (Mathieson et al., 2015; Allentoft et al., 2015; Ye et al., 2017), animals (Loog et al., 2017; Alves et al., 2019; Fages et al., 2019), and plants (Jaenicke-Despres et al., 2003; Ramos-Madrigal et al., 2016; Swarts et al., 2017). Simultaneous improvements in the recovery of aDNA have sparked increasing interest in going beyond demographic inference and opened the temporal dimension for studying past and present adaptation. Since the allele frequency trajectory through time from the incoming aDNA data itself provides us valuable information collect before, during and after genetic changes driven by selection, the additional temporal component has the promise of providing improved power for the inference of selection and its strength and timing of changes. Increased time resolution facilitates testing hypotheses on the drivers of past selection events such as domestication. See Dehasque et al. (2020) for a review.

The field of aDNA has recently experienced an exponential increase in terms of the amount of ancient and historical genomes sequenced and published, which has motivated a number of different statistical approaches for estimating selection coefficients and other population genetic quantities based on genetic time series data (*e*.*g*., Bollback et al., 2008; Malaspinas et al., 2012; Mathieson & McVean, 2013; Steinrücken et al., 2014; Foll et al., 2014, 2015; Shim et al., 2016; Schraiber et al., 2016; Ferrer-Admetlla et al., 2016; Mathieson, 2020; He et al., 2020a,b; Lyu et al., 2022; Mathieson & Terhorst, 2022; He et al., 2023a,b). Most of existing methods are built upon the hidden Markov model (HMM) framework of Williamson & Slatkin (1999), where the latent allele frequency trajectory of the population is characterised through the Wright-Fisher model (Fisher, 1922; Wright, 1931), and the observed allele frequency of the sample drawn from the underlying population at each sampling time point is modelled as a noisy observation of the latent population allele frequency. To remain computationally feasible, the Wright-Fisher model is typically approximated with its diffusion limit in the likelihood calculation, which was initiated by Bollback et al. (2008) and has been taken up later in Malaspinas et al. (2012), Steinrücken et al. (2014), Schraiber et al. (2016), Ferrer-Admetlla et al. (2016), He et al. (2020a,b), Lyu et al. (2022) and He et al. (2023a,b). These approaches have already been successfully applied in aDNA studies (*e*.*g*., Ludwig et al., 2009; Sandoval-Castellanos et al., 2017; Ye et al., 2017; Wutke et al., 2018). See Malaspinas (2016) for a review.

Despite numerous statistical approaches being developed for the inference of selection from aDNA data, most of them fail to consider the high error rate resulting from the damage of DNA molecules and the high missing rate arising from the fragmentation of DNA molecules, the two main characteristics of aDNA. The exceptions include Ferrer-Admetlla et al. (2016) that models genotype calling errors and He et al. (2020a) that allows genotype missing calls, which partially address this problem. More recently, He et al. (2023a) introduced a two-layer HMM framework, which allows their approach to work with the aDNA data in the form of genotype likelihoods, therefore enabling modelling sample uncertainty due to the damage and fragmentation of aDNA molecules. Working with genotype likelihoods by their method has been shown to boost power to infer selection from aDNA data, as compared to existing approaches based on inferred allele frequencies that do not model sample uncertainty. The procedure of He et al. (2023b), extended from He et al. (2023a), provides the flexibility of modelling linkage and epistasis in the inference of selection. However, both methods rely on the underlying assumption that mutation occurred prior to the initial sampling time point. That is, to use their methods we require the age of the mutant allele, which is typically unavailable, or exclude the samples drawn before the time that the mutant allele was first found in the sample as in He et al. (2023a,b), which can largely bias the inference of selection.

Here we introduce a novel Metropolis-Hastings (MH) within Gibbs framework for estimating temporally variable selection intensities from the aDNA data in the form of genotype likelihoods. Our approach is built upon the two-layer HMM framework (He et al., 2023a), therefore sample uncertainty arising from postmortem damage, high fragmentation, low coverage and small samples being considered. Our posterior computation is carried out through the MH-within-Gibbs framework (Tierney, 1994), in which we update the selection coefficient and the population allele frequency trajectory in the MH step with the particle marginal Metropolis-Hastings (PMMH) algorithm (Andrieu et al., 2010) and update the genotype of each sample individual in the Gibbs step. Our method therefore enables the reconstruction of underlying population allele frequency trajectories and the integration of temporal information in genotype likelihood corrections. To fix the aforementioned issues encountered in He et al. (2023a,b), we propose a numerical scheme to simulate the Wright-Fisher diffusion backwards in time and mix the forward- and backward-in-time simulations in our particle filter likelihood computation, which also allows us to jointly infer the timing when the mutant allele is created and/or lost in the underlying population.

We repeat the same simulation studies conducted in He et al. (2023a) to evaluate the performance of our method in detecting selection signatures, estimating selection coefficients, testing selection changes and correcting genotype likelihoods from the aDNA data in the form of genotype likelihoods, especially in poor quality (*i*.*e*., high missing rate and error rate). To show the applicability of our procedure, we reanalyse the aDNA data from pigmentation loci in ancient horses (Wutke et al., 2016).

## 2. Materials and Methods

In this section, we begin with a brief introduction of the Wright-Fisher diffusion for a single locus evolving subject to selection over time. We then detail our procedure for Bayesian inference of temporally variable selection from the aDNA data in the form of genotype likelihoods.

### 2.1. Wright-Fisher diffusion

We assume a diploid population of *N* (*t*) randomly mating individuals at time *t*, where *t* is measured in a unit of 2*N*_0_ generations for an arbitrary reference population size *N*_0_ that is fixed through time. At the locus of interest, denoted by 𝒜, there are two possible allele types, labelled 𝒜_0_ and 𝒜_1_, respectively. The symbol 𝒜_0_ represents the ancestral allele that originally exists in the population, and the symbol 𝒜_1_ denotes the mutant allele that arises in the population only once with the mutation rate *u* per generation. Suppose that selection takes the form of viability selection, and then the relative viabilities of the 𝒜_0_ 𝒜_0_, 𝒜_0_ 𝒜_1_ and 𝒜_1_ 𝒜_1_ genotypes are set to 1, 1 + *hs* and 1 + *s* per generation, respectively, where *s* ∈ [−1, +∞) is the selection coefficient and *h* ∈ [0, 1] is the dominance parameter.

As in He et al. (2023a), the mutant allele frequency trajectory of the population is modelled by the Wright-Fisher diffusion with selection (*i*.*e*., a standard diffusion limit of the Wright-Fisher model with selection). More specifically, we let *X* be the Wright-Fisher diffusion with selection, which models the mutant allele frequency evolving in the state space [0, 1] under selection and satisfies the stochastic differential equation (SDE)

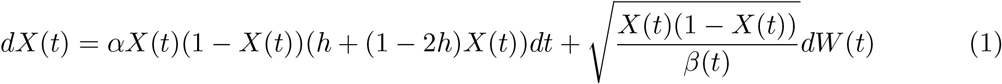

for *t* ≥ *t*_0_ with *X*(*t*_0_) = *x*_0_, where *α* = 2*N*_0_*s* and *β*(*t*) = *N* (*t*)*/N*_0_ as *N*_0_ goes to infinity, and *W* is the standard Brownian motion.

### 2.2. Bayesian inference of selection

We assume that the available data are always sampled from the underlying population at a finite number of distinct time points, say *t*_1_ *< t*_2_ *<* … *< t*_*K*_. At the *k*-th sampling time point, *N*_*k*_ individuals are drawn from the underlying population, and for individual *n*, let ***r***_*n,k*_ denote, in this generic notation, all of the reads at the locus of our interest. To match the timescale of the Wright-Fisher diffusion, the sampling time points are measured in units of 2*N*_0_ generations. The population genetic quantities of our interest in this work are the selection coefficient *s* and the dominance parameter *h*, represented by ***ϑ*** = (*s, h*) for ease of notation.

#### 2.2.1. Hidden Markov model

Our method is based on the two-layer HMM framework of He et al. (2023a) for the data on aDNA sequences (see Figure 1 for its graphical representation), where the first hidden layer *X* represents the frequency trajectory of the mutant allele in the population through time under the Wright-Fisher diffusion in Eq. (1), the second hidden layer ***G*** is the genotype of the individual in the sample, and the third observed layer ***R*** is the data on aDNA sequences.

**Figure 1.**
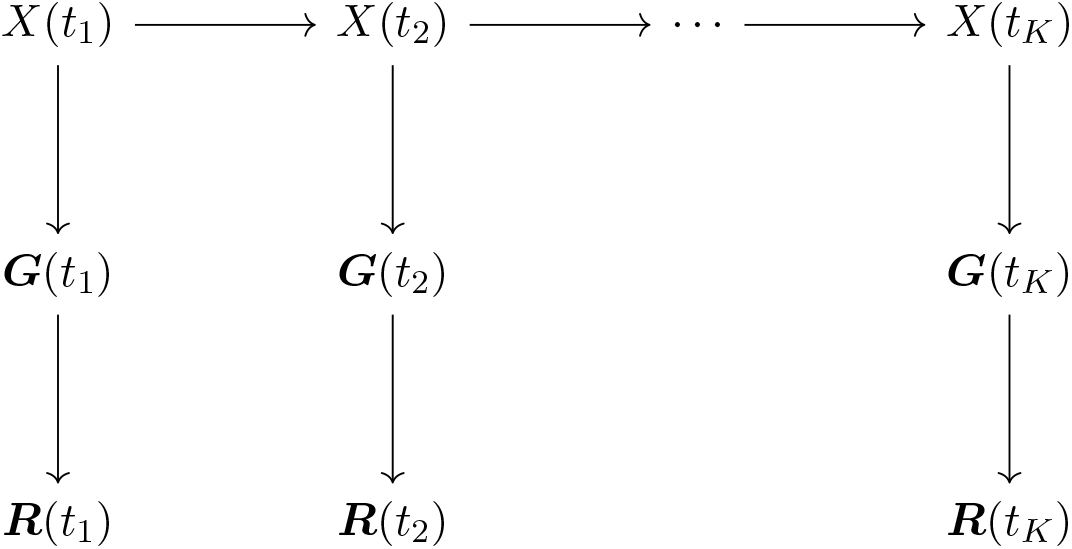
Graphical representation of the two-layer HMM framework introduced by He et al. (2023a) for the data on aDNA sequences.

We let ***x***_1:*K*_ = {*x*_1_, *x*_2_, …, *x*_*K*_} be the mutant allele frequency trajectory of the population at the sampling time points ***t***_1:*K*_ and ***g***_1:*K*_ = {***g***_1_, ***g***_2_, …, ***g***_*K*_ } be the genotypes of the individuals drawn from the population at the sampling time points ***t***_1:*K*_, where 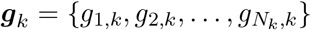 with *g*_*n,k*_ ∈ {0, 1, 2} representing the number of mutant alleles in individual *n* at sampling time point *t*_*k*_. From the two-layer HMM framework shown in Figure 1, we have

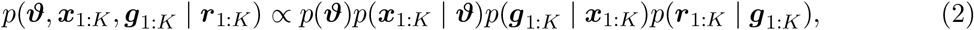

where ***r***_1:*K*_ = {***r***_1_, ***r***_2_, …, ***r***_*K*_} with 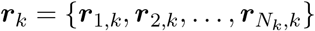.

In Eq. (2), we assume the prior *p*(***ϑ***) to be a uniform distribution over the parameter space. We decompose the probability distribution *p*(***x***_1:*K*_ | ***ϑ***) as

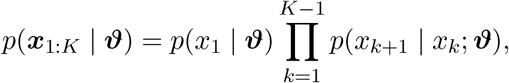

where *p*(*x*_1_ | ***ϑ***) can be an arbitrary prior distribution over the state space, and *p*(*x*_*k*+1_ | *x*_*k*_; ***ϑ***) satisfies the Kolmogorov backward equation (or its adjoint) associated with the Wright-Fisher diffusion in Eq. (1). We decompose the probability distribution *p*(***g***_1:*K*_ | ***x***_1:*K*_) as

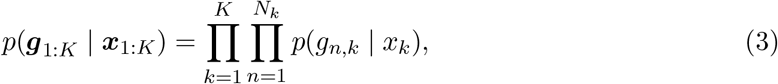

where

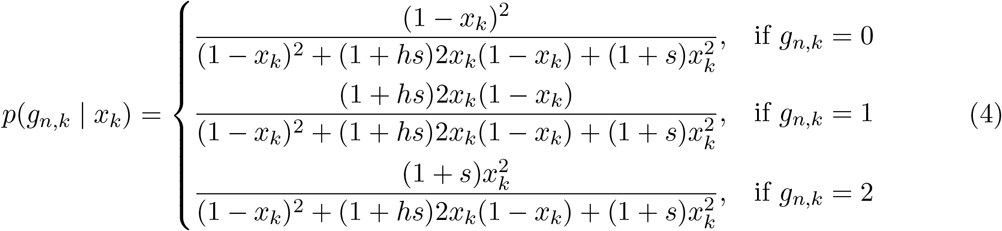

under the assumption that all individuals in the sample are drawn from the population in their adulthood (*i*.*e*., the stage after selection but before reproduction in the life cycle, see He et al. (2017)). We decompose the probability distribution *p*(***r***_1:*K*_ | ***g***_1:*K*_) as

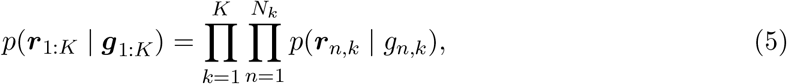

where *p*(***r***_*n,k*_ | *g*_*n,k*_), known as genotype likelihoods.

#### 2.2.2. Particle marginal Metropolis-Hastings within Gibbs

Given the decomposition of the posterior *p*(***ϑ, x***_1:*K*_, ***g***_1:*K*_ | ***r***_1:*K*_) in Eq. (2), Gibbs sampling (Turchin, 1971; Geman & Geman, 1984) is a natural framework for the posterior computation. We partition the model components into three blocks: the population genetic parameters ***ϑ***, the population mutant allele frequency trajectory ***x***_1:*K*_ and the sample individual genotypes ***g***_1:*K*_, and their corresponding full conditional probability distributions can then be formulated as

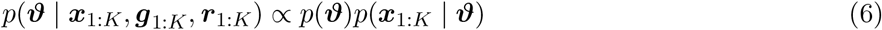

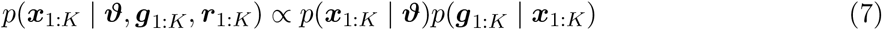

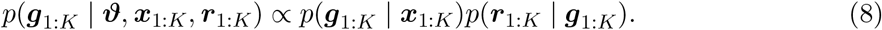

From Eqs. (6)–(8), we see that the full conditional probability distribution *p*(***g***_1:*K*_ | ***ϑ, x***_1:*K*_, ***r***_1:*K*_) is available with Eqs. (3)–(5), but the remaining two are computationally infeasible to generate from. We therefore resort to the MH-within-Gibbs algorithm introduced by Tierney (1994) as a hybrid framework that incorporates the MH algorithm (Metropolis et al., 1953; Hastings, 1970) to implement the sampling steps when Gibbs sampling fails. More specifically, each iteration we update the sample individual genotypes ***g***_1:*K*_ using the full conditional probability distribution *p*(***g***_1:*K*_ | ***ϑ, x***_1:*K*_, ***r***_1:*K*_) in the Gibbs step and update the population genetic parameters ***ϑ*** and the population mutant allele frequency trajectory ***x***_1:*K*_ in the MH step.

In the MH-within-Gibbs algorithm, the population genetic parameters ***ϑ*** and the population mutant allele frequency trajectory ***x***_1:*K*_ are commonly updated in two separate MH steps each iteration. Following He et al. (2023a), through the PMMH algorithm (Andrieu et al., 2010), we jointly update the population genetic parameters ***ϑ*** and the population mutant allele frequency trajectory ***x***_1:*K*_. More specifically, the marginal likelihood

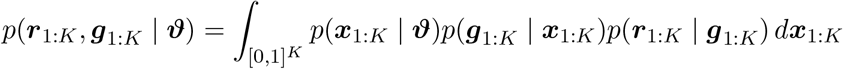

is approximated with the bootstrap particle filter (Gordon et al., 1993) in the PMMH procedure, where particles evolve under the Wright-Fisher SDE in Eq. (1) by the Euler-Maruyama scheme. The product of the average weights of the set of particles at the sampling time points ***t***_1:*K*_ yields an unbiased estimate of the marginal likelihood *p*(***r***_1:*K*_, ***g***_1:*K*_ | ***ϑ***), denoted by 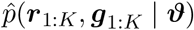, and the population mutant allele frequency trajectory ***x***_1:*K*_ is sampled once from the final set of particles with their corresponding weights (see He et al. (2023a) for details).

For clarity, we write down the PMMH-within-Gibbs algorithm for our posterior computation:

Step 1: Initialise ***ϑ, x***_1:*K*_ and ***g***_1:*K*_ :

Step 1a: Draw 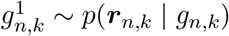 for *n* = 1, 2, …, *N*_*k*_ and *k* = 1, 2, …, *K*.

Step 1b: Draw ***ϑ***^1^ ∼ *p*(***ϑ***).

Step 1c: Run a bootstrap particle filter with ***ϑ***^1^ and 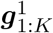 to get 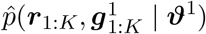 and 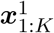.

Repeat Steps 2 and 3 until a sufficiently large sample of ***ϑ, x***_1:*K*_ and ***g***_1:*K*_ have been obtained:

Step 2: Update ***g***_1:*K*_ (Gibbs step):

Step 2a: Draw 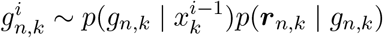 for *n* = 1, 2, …, *N*_*k*_ and *k* = 1, 2, …, *K*.

Step 3: Update ***ϑ*** and ***x***_1:*K*_ (MH step):

Step 3a: Draw ***ϑ***^*i*^ ∼ *q*(***ϑ*** | ***ϑ***^*i*−1^).

Step 3b: Run a bootstrap particle filter with ***ϑ***^*i*^ and 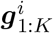 to get 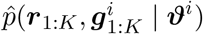 and 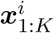.

Step 3c: Accept ***ϑ***^*i*^ and 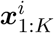 with

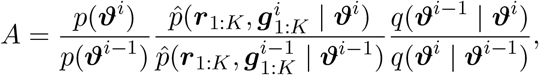

otherwise set ***ϑ***^*i*^ = ***ϑ***^*i*−1^ and 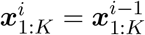.

Note that the superscript *i* represents the iteration label in our procedure presented above. We use random walk proposals for each component of the population genetic parameters ***ϑ*** in what follows unless otherwise specified.

With a large enough sample of the population genetic parameters ***ϑ***, the population mutant allele frequency trajectory ***x***_1:*K*_ and the sample individual genotypes ***g***_1:*K*_, we can yield the maximum a posteriori probability (MAP) estimates for the population genetic parameters ***ϑ*** By calculating the mode of the posterior *p*(***ϑ*** | ***r***_1:*K*_), which can be computed with the samples of the population genetic parameters through nonparametric density estimation techniques (see, *e*.*g*., Izenman, 1991). We can take the mean of the samples of the population mutant allele frequency trajectory to be the estimate for the population mutant allele frequency trajectory ***x***_1:*K*_ and the mean of the samples of the sample individual genotypes to be the posterior probabilities for the sample individual genotypes ***g***_1:*K*_.

Following He et al. (2023a), we allow the selection coefficient *s* to be piecewise constant over time, *e*.*g*., let the selection coefficient *s*(*t*) = *s*^−^ if *t < τ* otherwise *s*(*t*) = *s*^+^, where *τ* is the time of an event that might change selection such as the times of plant and animal domestication. Our procedure can be directly employed to estimate the selection coefficients *s*^−^ and *s*^+^ for any prespecified time *τ*, which can be achieved by simulating the Wright-Fisher diffusion in Eq. (1) with the selection coefficient *s*^−^ for *t < τ* and *s*^+^ for *t* ≥ *τ*, respectively, in the bootstrap particle filter. In this case, our framework can also test the hypotheses *H*_0_ : Δ*s* = 0 against *H*_1_ : Δ*s* ≠ 0 through the posterior *p*(Δ*s* | ***r***_1:*K*_), where Δ*s* = *s*^+^ − *s*^−^ is the change in the selection coefficient at time *τ*. Our procedure naturally lends itself to allowing multiple events that might change selection.

#### 2.2.3. Backward-in-time simulation

In our PMMH-within-Gibbs procedure, the marginal likelihood is calculated through a bootstrap particle filter, where particles are simulated according to the Wright-Fisher SDE in Eq. (1) by the Euler-Maruyama scheme. The forward-in-time simulation of the Wright-Fisher diffusion will result in the same issue encountered in He et al. (2023a) that only samples drawn after the time that mutation occurred can be taken as input, but the age of the mutant allele is commonly not available in practice. To address this issue, we introduce a novel numerical scheme for simulating the Wright-Fisher diffusion backwards in time to replace the forward-in-time simulation in the bootstrap particle filter described above. See Griffiths (2003) and Coop & Griffiths (2004) for alternative methods of the backward-in-time simulation of the Wright-Fisher diffusion.

Suppose that we only move in one time step Δ_*t*_ from *x* to *x* or *x ±* Δ_*x*_, *i*.*e*., move to one of the neighbouring points of *x* if moving at all, as long as we match up the mean and variance with those of the Wright-Fisher SDE in Eq. (1), then as the step sizes Δ_*t*_ and Δ_*x*_ become finer, the numerical approximation still converges to the Wright-Fisher diffusion *X*. More specifically, given *X*(*t*) = *x*, we let *p*^−^(*t, x*) and *p*^+^(*t, x*) denote the probability that *X*(*t* + Δ_*t*_) is at 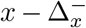 and 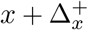, respectively, which are the two neighbouring points of *x*. For generality, we allow 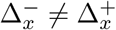. Let Δ*X*(*t*) = *X*(*t* + Δ_*t*_) − *X*(*t*), and then we have

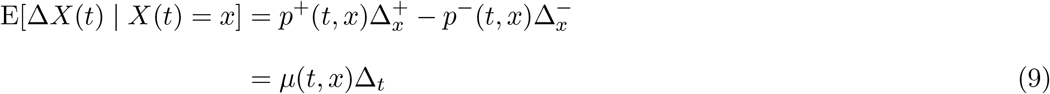

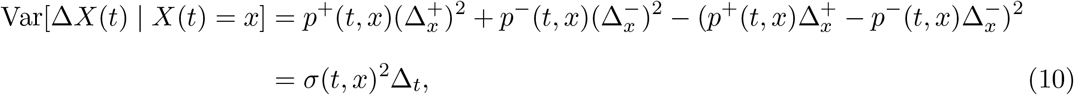

where *μ*(*t, x*) = *αx*(1 − *x*)(*h* + (1 − 2*h*)*x*) and 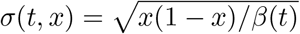. From Eq. (9), we have 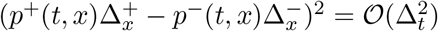, so we can drop it in Eq. (10) and simply solve

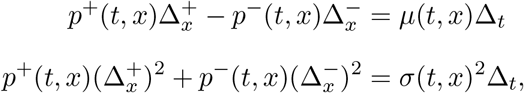

whence we obtain

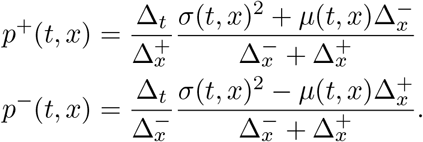

Given *X*(*t* + Δ_*t*_) = *x*′, as a function of *x*, Pr(*X*(*t* + Δ_*t*_) = *x*′| *X*(*t*) = *x*) is non-zero at only three points, 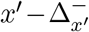, *x* and 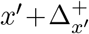, with values equal to 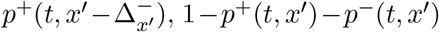 and 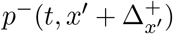, respectively. Let

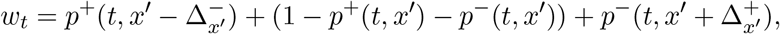

and then we can approximate this measure (not necessarily a probability measure) by drawing a value *x* for *X*(*t*) according to the probability distribution

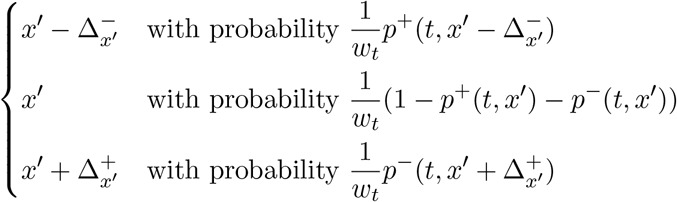

and assigning weight *w*_*t*_ to *x*. The weights at every time step multiply. Note that for simplicity, we fix the grid point closest to 0 to be 1*/*(2*N*), thereby 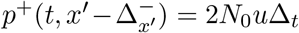 if *x*′= 1*/*(2*N*). With the backward-in-time simulation of the Wright-Fisher diffusion in the bootstrap particle filter, our method can avoid the dependence of the timing when the mutant allele is created but will require the timing when the mutant allele is lost, which is also commonly unavailable in practice. To circumvent the issues arising from the forward- and backward-in-time simulations, we can mix the forward- and backward-in-time simulations of the Wright-Fisher diffusion in the bootstrap particle filter. More specifically, we simulate the Wright-Fisher diffusion forwards in time from the sampling time point when the individual with the highest likelihood of containing at least one copy of the mutant allele is collected, prior to which we simulate the Wright-Fisher diffusion backwards in time. Also, we can co-estimate the timing when the mutant allele is created if we run the backward-in-time simulation until the mutant allele frequency being absorbed at 0 and the timing when the mutant allele is lost if we run the forward-in-time simulation until the mutant allele frequency being absorbed at 0.

## 3. Results

In this section, we repeat the same simulation studies as those performed in He et al. (2023a) to evaluate the performance of our approach. We also apply our method to reanalyse the aDNA data from previous studies of Ludwig et al. (2009), Pruvost et al. (2011) and Wutke et al. (2016), where they collected and sequenced a total of 201 ancient horse samples ranging from a pre-to a post-domestication period for pigmentation loci. We assume a single event that might change selection unless otherwise specified.

### 3.1. Performance evaluation

In our simulation studies, following He et al. (2023a), we consider a total of 801 generations starting from generation 0 and draw a sample of 10 individuals every 40 generations, 210 sample individuals in total that approaches the size of ancient horse samples in Wutke et al. (2016). We assume that the only event that might change selection occurs in generation 350, and therefore take the selection coefficient to be *s*(*k*) = *s*^−^ for *k <* 350 otherwise *s*(*k*) = *s*^+^, where we draw the selection coefficients *s*^−^ and *s*^+^ uniformly from [−0.05, 0.05]. We take the dominance parameter to be *h* = 0.5 (*i*.*e*., assuming codominance) and use a bottleneck demographic history, like the demographic history of horses reported in Der Sarkissian et al. (2015), with the population size *N* (*k*) = 32000 for *k <* 200, *N* (*k*) = 8000 for 200 ≤ *k <* 400, and *N* (*k*) = 16000 for *k* ≥ 400.

We run a group of simulations to test our method for different scenarios, where we consider the selection coefficients *s*^−^ *<* 0, *s*^−^ = 0, *s*^−^ *>* 0 and *s*^+^ *< s*^−^, *s*^+^ = *s*^−^, *s*^+^ *> s*^−^, respectively, giving rise to nine possible combinations in total. For each combination, we produce 200 datasets in the form of genotype likelihoods through the following procedure:

Repeat Step 1 until 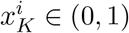 for *i* = 1, 2, …, 10000:

Step 1: Generate *s*^−^, *s*^+^ and *x*_1_.

Step 1a: Draw *s*^−^, *s*^+^ from a uniform distribution over [−0.05, 0.05] and *x*_1_ from a uniform distribution over [0.1, 0.9].

Step 1b: Simulate 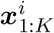 ith *s*^−^, *s*^+^ and *x*_1_ through the Wright-Fisher model with selection for *i* = 1, 2, …, 10000.

Step 2: Simulate ***x***_1:*K*_ with *s*^−^, *s*^+^ and *x*_1_ through the Wright-Fisher model with selection.

Repeat Step 3 for *k* = 1, 2, …, *K*:

Step 3: Generate *p*(***r***_*n,k*_ | *g*) for *g* = 0, 1, 2 and *n* = 1, 2, …, *N*_*k*_:

Step 3a: Draw *g*_*n,k*_ ∼ *p*(*g* | *x*_*k*_) in Eq. (4).

Step 3b: Draw *p*(***r***_*n,k*_ | *g*) for *g* = 0, 1, 2 from a Dirichlet distribution of order 3 with 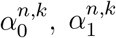, and 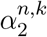.

Note that the superscript *i* represents the replicate label. Such a procedure avoids the mismatch between the underlying model in the data generation process and that in the selection inference procedure (He et al., 2023a). We set the parameter 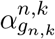 to *ϕψ* and the other two to (1 − *ϕ*)*ψ/*2, where *ϕ* and *ψ* are the parameters that control the quality of the simulated dataset in terms of the missing rate (MR) and error rate (ER) with a common threshold for genotype calling (*i*.*e*., 10 times more likely, see Kim et al., 2011). Here we set the parameters *ϕ* = 0.85 and *ψ* = 0.5, giving rise to an average MR of 20.39% with a standard deviation (SD) of 2.79% and an average ER of 5.28% with an SD of 1.53%. The performance evaluation results for other data qualities can be found in the supplement.

For each simulated dataset, we pick a uniform prior over [−1, 1] for both selection coefficients *s*^−^ and *s*^+^, and adopt a reference population size of *N*_0_ = 16000. We run 10000 PMMH-within-Gibbs iterations with 1000 particles. We discard 5000 burn-in iterations and thin the remaining samples by recording every fifth value. In the particle filter, we run three different simulations of the Wright-Fisher diffusion, which are the full forward-in-time simulation, the full backward-in-time simulation and the combination of forward- and backward-in-time simulations, respectively. In the forward-in-time simulation with the Euler–Maruyama scheme, each generation is divided into five subintervals. In the backward-in-time simulation with our numerical scheme presented in Section 2.2.3, each generation is partitioned into 31 subintervals and the state space [0, 1] is divided into 1177 subintervals with the mutation rate *u* = 10^−8^. In the combination of forward-and backward-in-time simulations, we run a forward-in-time simulation from generation 280 to 800 and a backward-in-time simulation from generation 280 to 0, respectively. We also run the procedure of He et al. (2023a) with the same settings for each simulated dataset as comparison. In our simulation studies, the performance of our PMMH-within-Gibbs procedure with these three different simulation strategies is expected to be similar since our data generating mechanism ensures that in the underlying population the mutant allele arises before the first sampling time point and survival till the last sampling time point. We will show and discuss the improvement brought about by the mix of the forward- and backward-in-time simulations in Section 4. Moreover, our procedure is expected to have a better performance in the inference of post-event selection than pre-event selection since more samples are drawn after the occurrence of the event and distributed over a longer time period.

#### 3.1.1. Performance in detecting selection signatures

Figure 2 shows the performance of our method in detecting selection signatures for different selection scenarios, presented in the receiver operating characteristic (ROC) curves. The ROC curve is produced by plotting the true-positive rate (TPR) against the false-positive rate (FPR), where the TPR and FPR are achieved for each value of the posterior probability for the selection coefficient being used as a threshold to classify a locus as being selected. We calculate the area under the ROC curve (AUC) to measure the performance. We find from Figure 2 that all ROC curves of our method are located close to the upper left hand corner in the ROC space, and their AUC values vary from 0.76 to 0.79 for pre-event selection and from 0.91 to 0.93 for post-event selection, which means that our method has good ability to discriminate between negative and non-negative selection (or between positive and non-positive selection). Although our procedure does not perform as well as the approach of He et al. (2023a) in identifying selection signatures, the performance difference will decline and may even be eliminated as the data quality improves (see Supporting Information, Figure S1).

**Figure 2.**
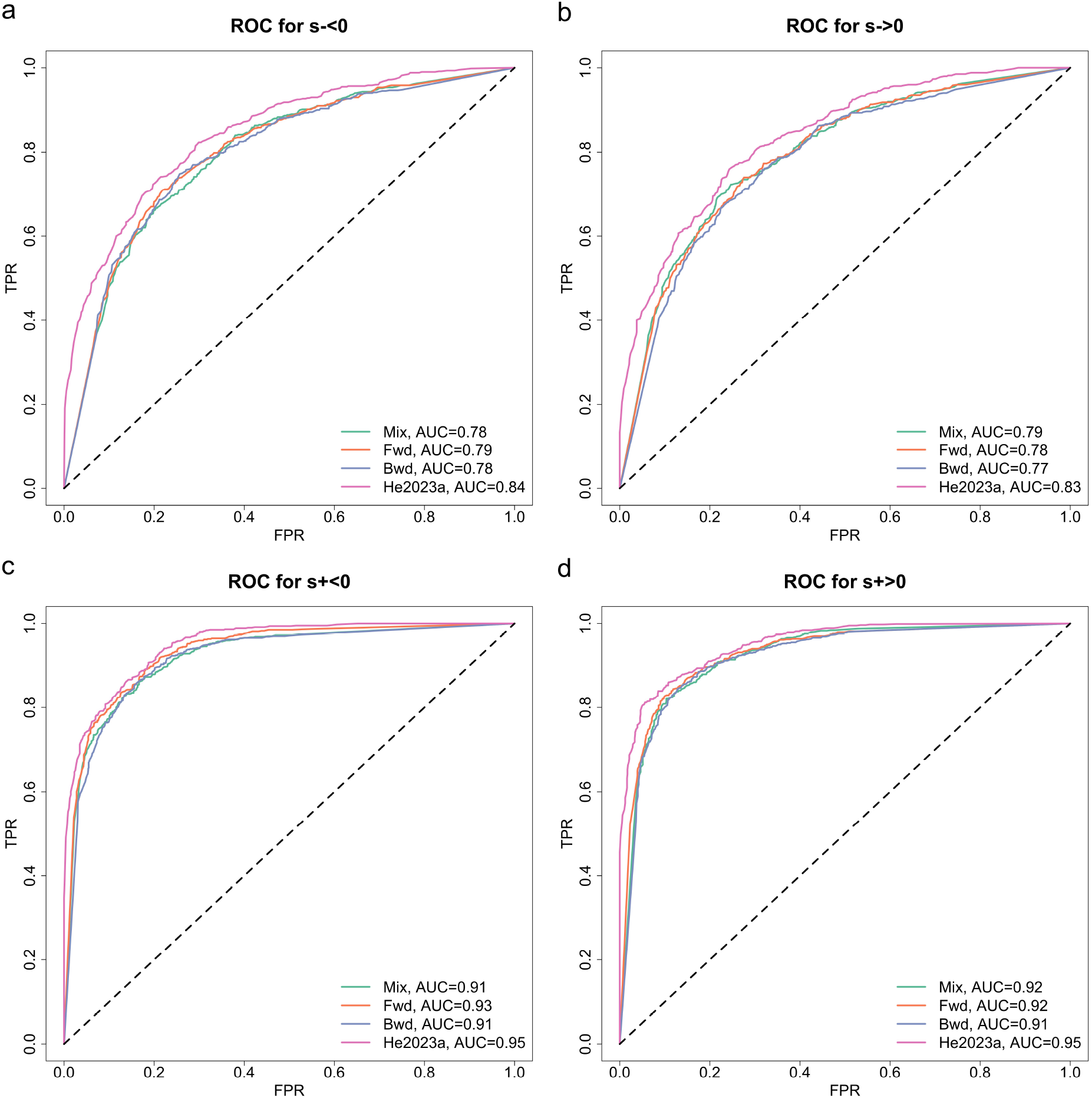
ROC curves for detecting selection signatures across different selection scenarios.

#### 3.1.2. Performance in estimating selection coefficients

Figure 3 shows the performance of our method in estimating selection coefficients for different selection scenarios, presented in the boxplots of the bias in our estimation of selection coefficients with the box as the first and third quartiles with the median in the middle and the tips of the whiskers as the 2.5%-quantile and the 97.5%-quantile, respectively. See Supporting Information, Table S1, for the bias and root mean square error (RMSE). As shown in Figure 3, our approach can produce nearly median-unbiased estimates across all selection scenarios. As compared to the method of He et al. (2023a), our procedure has similar bias but larger uncertainty. Combining with the boxplot results for other data qualities (see Supporting Information, Figure S2 and Table S2), we find that our approach has smaller bias than the method of He et al. (2023a) but still with larger uncertainty when the data quality deteriorates, and the bias and uncertainty of our estimate tend to be close to those of the method of He et al. (2023a) when the data quality improves.

**Figure 3.**
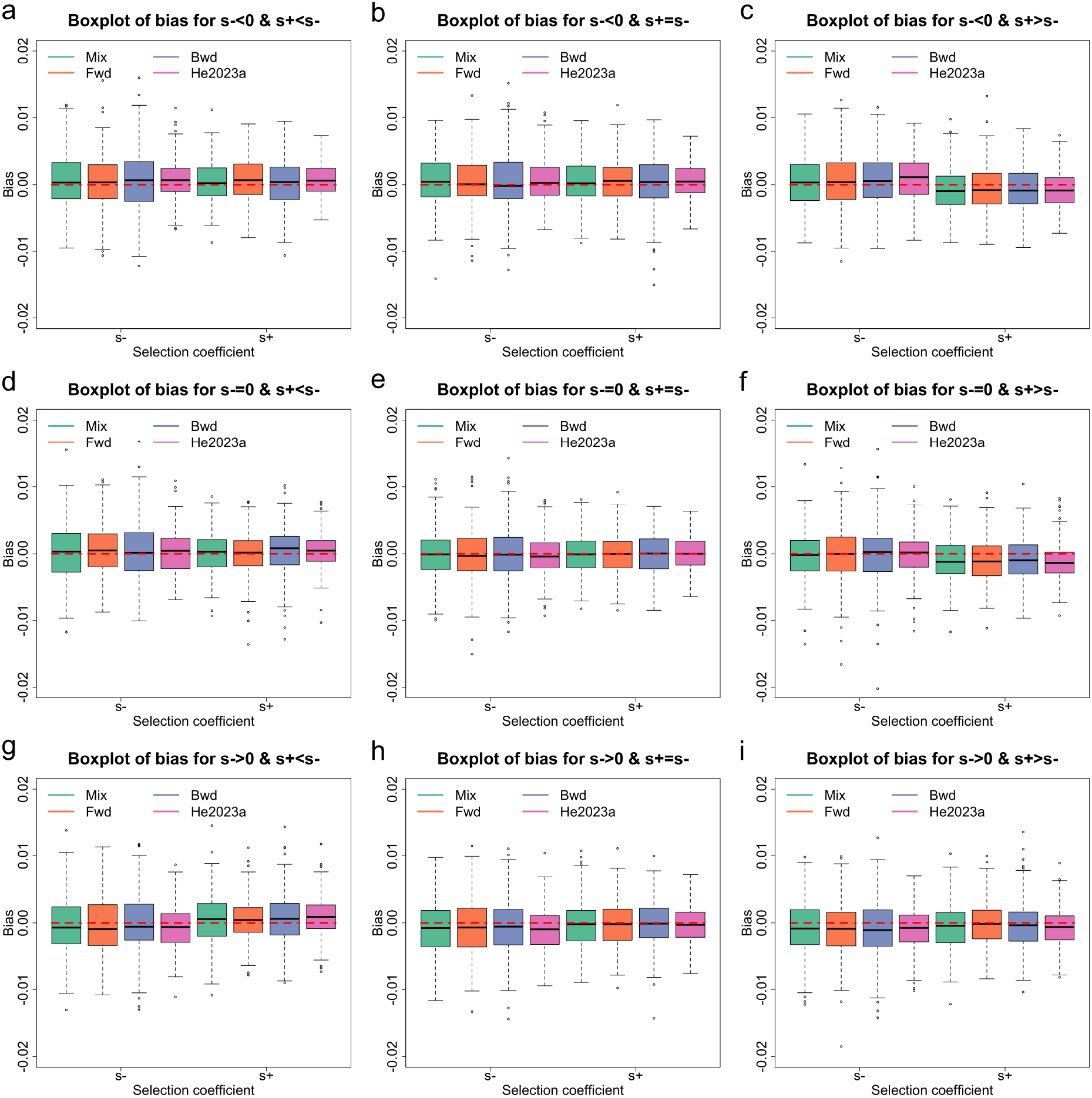
Boxplots for the bias of the selection coefficient estimates across different selection scenarios.

#### 3.1.3. Performance in testing selection changes

Figure 4 illustrates the performance of our method in testing selection changes for different selection scenarios, presented in the ROC curves with their AUC values, where we compute the TPR and FPR for each value of the posterior probability for the selection change being used as a threshold to classify a locus as experiencing a shift in selection. As shown in Figure 4, our procedure demonstrates superior performance in discriminating between a negative change and a non-negative change (or a positive change and a non-positive change) in selection. Similar to selection signature detection, the approach of He et al. (2023a) exhibits better performance in testing selection changes than our method, but the gap will narrow and may even be eliminated when the data quality improves (see the Supporting Information, Figure S3).

**Figure 4.**
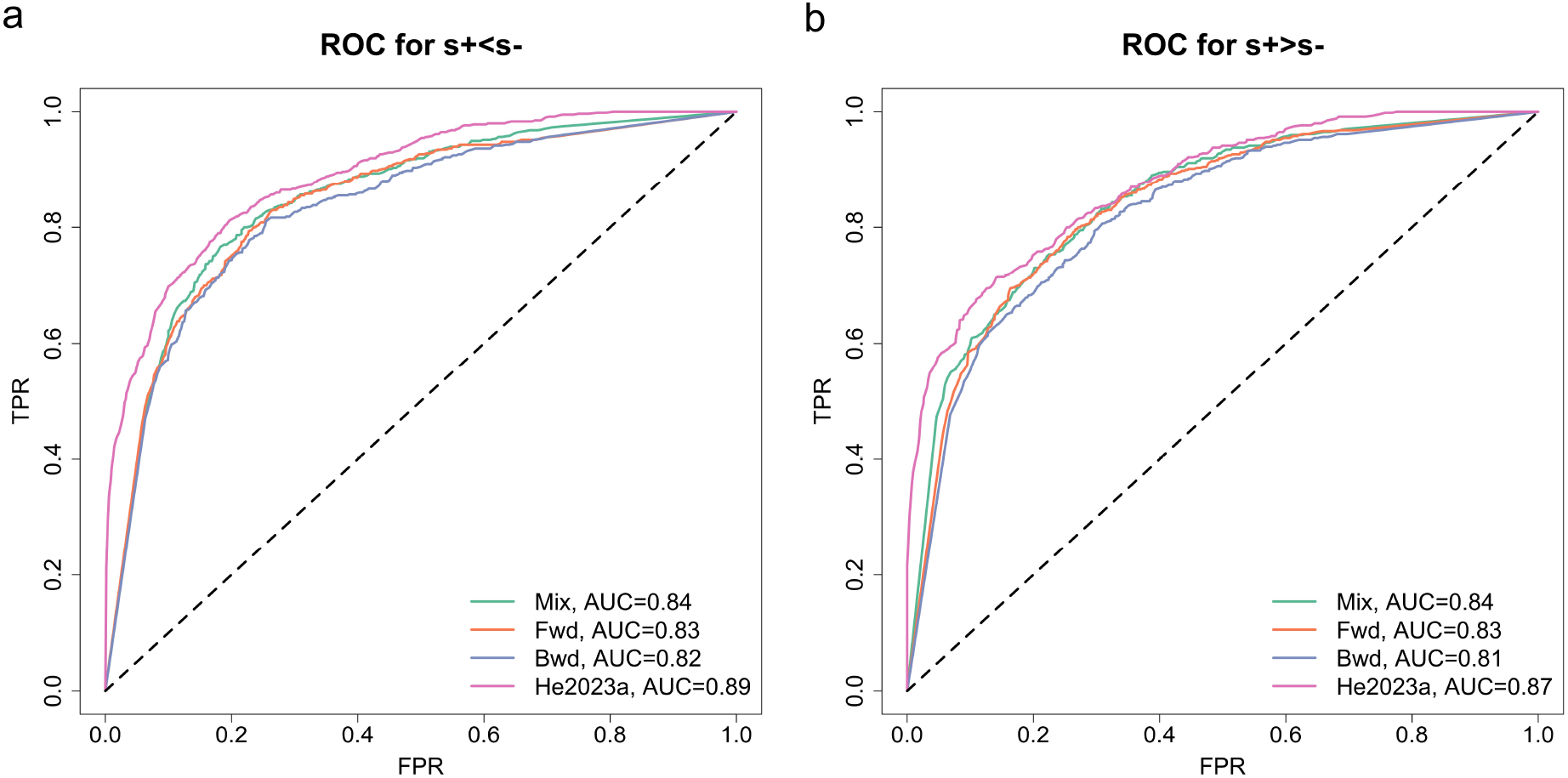
ROC curves for testing selection changes across different selection scenarios.

#### 3.1.4. Performance in correcting genotype likelihoods

Figure 5 shows the performance of our method in correcting genotype likelihoods for different selection scenarios, presented in the boxplots of the MR and ER computed with the 10 times more likely threshold. The mean and SD of the MR and ER are left in Supporting Information, Table S3. We find from Figure 5 that the MR and ER of genotype posteriors are smaller than those of genotype likelihoods across all selection scenarios, *i*.*e*., a 12.16% decline in the average MR and a 12.13% fall in the average ER compared to genotype likelihoods. Combining with the boxplot results for other data qualities (see Supporting Information, Figure S4 and Table S4), we observe that genotype posteriors enable the average MR to drop by 10.40%–19.29% and the average ER to decline by 1.42%–12.52% compared to genotype likelihoods, whose average MR and ER vary from 7.48% to 45.44% and from 0.53% to 9.60%, respectively.

**Figure 5.**
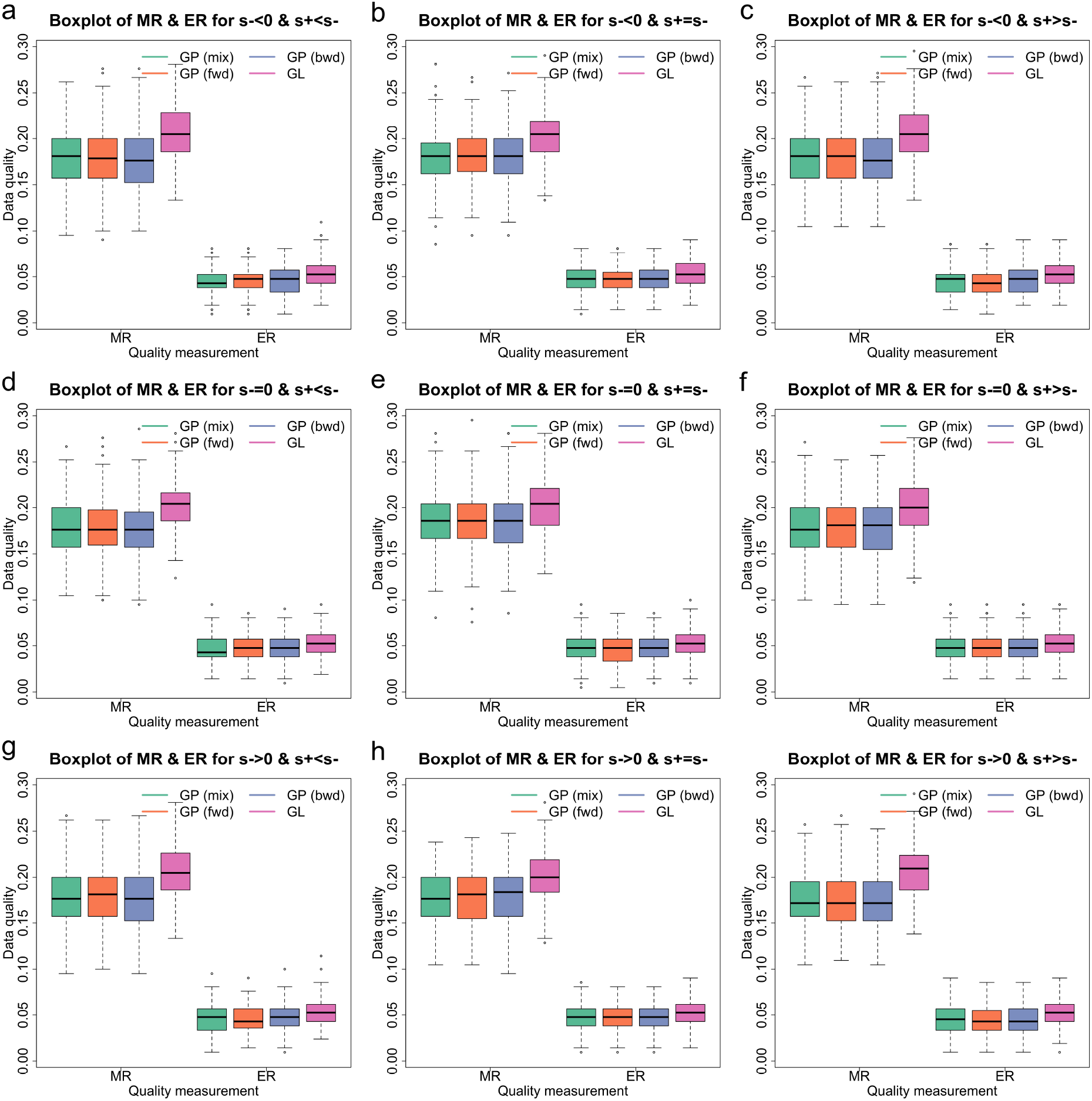
Boxplots for the AMR and AER of the genotype posteriors across different selection scenarios. GP and GL are shorthands for genotype posterior and genotype likelihood.

### 3.2. Selection of horse coat colouration

We employ our procedure to infer selection acting on the *MC1R, KIT13* and *TRPM1* genes, responsible for the chestnut coat colour, tobiano coat pattern and leopard complex coat pattern in horses, respectively, with the data on aDNA sequences from previous studies of Ludwig et al. (2009), Pruvost et al. (2011) and Wutke et al. (2016), which have been illustrated to be involved to ecological, environmental and cultural shifts (Ludwig et al., 2009, 2015; Wutke et al., 2016). We explore the hypothesis that human preferences for horse coat colours and patterns changed during the Middle Ages when horses started to be differentiated by use (see Wutke et al., 2016, and references therein).

Given that the ancestors of modern domestic horses only spread out from the Pontic–Caspian steppe 4200 years ago when they promptly started to become dominant across Eurasia (Librado et al., 2021), we restrict our analysis to the ancient horse samples no older than 4200 years ago as in He et al. (2023b). To convert called genotypes to genotype likelihoods, following He et al. (2023a), for each sample individual we assign 1 to the called genotype and 0 to the other two if the genotype is called; otherwise, we assign equal genotype likelihoods to all possible (ordered) genotypes that are normalised to sum 1. See Supporting Information, Table S5, for the genotype likelihoods.

We adopt a generation of eight years and a mutation rate of *u* = 7.24 *×* 10^−9^ per generation reported by Orlando et al. (2013). We incorporate the time-varying demographic history of the horse population estimated by Der Sarkissian et al. (2015) and pick a reference population size of *N*_0_ = 16000. For each gene, we take the dominance parameter *h* to be that reported in Wutke et al. (2016). In the PMMH-within-Gibbs, we run 20000 iterations with a burn-in of 10000 and a thinning of 5. In the particle filter, we mix the forward- and backward-in-time simulations of the Wright-Fisher diffusion, *i*.*e*., for each gene, we run a forward-in-time simulation from the time that the mutant allele was first observed in the sample, before which we run a backward-in-time simulation till the starting sampling time point. The other settings are the same as we used in Section 3.1. The selection coefficient estimates with their 95% highest posterior density (HPD) intervals are summarised in Supporting Information, Table S6, and the genotype posteriors are recorded in Supporting Information, Table S7.

#### 3.2.1. Selection of the chestnut coat colour

The horse gene *MC1R* is located on chromosome 3, with mutations coding for the recessive chestnut coat (Corbin et al., 2020). The resulting posteriors for *MC1R* are shown in Figure 6. Our estimate of the selection coefficient for the *MC1R* mutation is 0.0116 with 95% HPD interval [0.0039, 0.0227] in pre-medieval horses and 0.0092 with 95% HPD interval [−0.0238, 0.0311] in medieval horses. Our estimates show that the *MC1R* mutation was positively selected since the domestication of modern horses (*i*.*e*., the posterior probability for positive selection is 0.995 for pre-medieval horses but 0.694 for medieval horses). Our posterior for the change in the selection coefficient is approximately symmetric about 0 (see Figure 6d), which suggests that the evidence to reject the hypothesis that no change occurred in selection of the *MC1R* mutation during the Middle Ages is not sufficient. As illustrated in Figure 6c, we see a significant consecutive increase in the *MC1R* mutation frequency from modern horse domestication.

**Figure 6.**
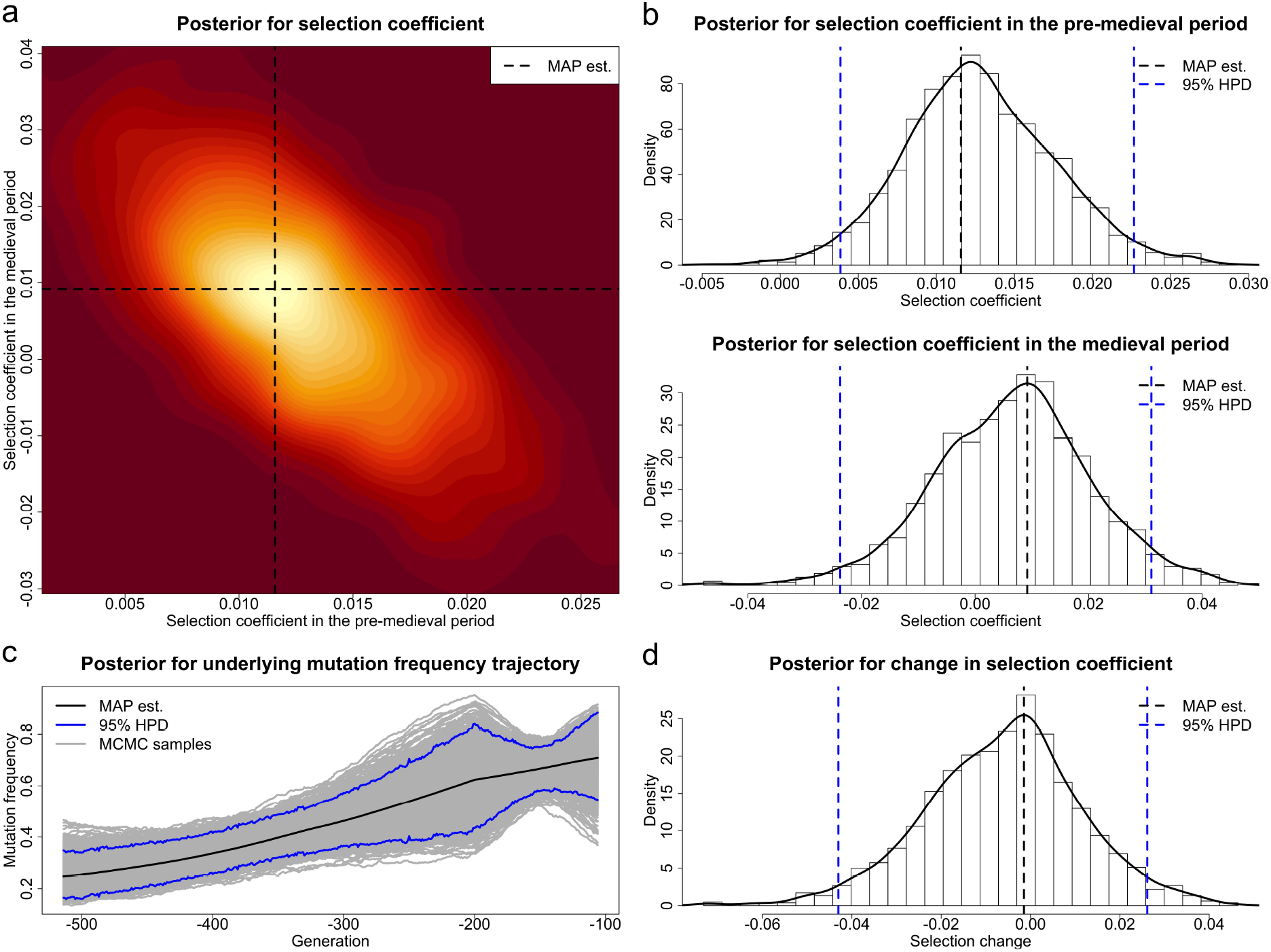
Posteriors for the selection coefficients of the *MC1R* mutation in the pre-medieval and medieval period and the underlying frequency trajectory of the *MC1R* mutation in the population.

#### 3.2.2. Selection of the tobiano coat pattern

The horse gene *KIT13* is located on chromosome 3, with mutations coding for the dominant tobiano coat (Brooks et al., 2007). The resulting posteriors for *KIT13* are illustrated in Figure 7. During the pre-medieval period, our estimate of the selection coefficient for the *KIT13* mutation is 0.0100 with 95% HPD interval [0.0010, 0.0190], strong evidence for positive selection acting on pre-medieval horses with the posterior probability for positive selection being 0.984. During the medieval period, our estimate of the selection coefficient for the *KIT13* mutation is −0.0544 with 95% HPD interval [−0.0922, −0.01335], strong evidence for negative selection acting on medieval horses with the posterior probability for negative selection being 1.000. The posterior probability for a negative shift occurring in selection of the *KIT13* mutation during the Middle Ages is 1.000. Our estimate for the underlying *KIT13* mutation frequency trajectory shows an increase in the *KIT13* mutation frequency from modern horse domestication followed by a drop during the Middle Ages.

**Figure 7.**
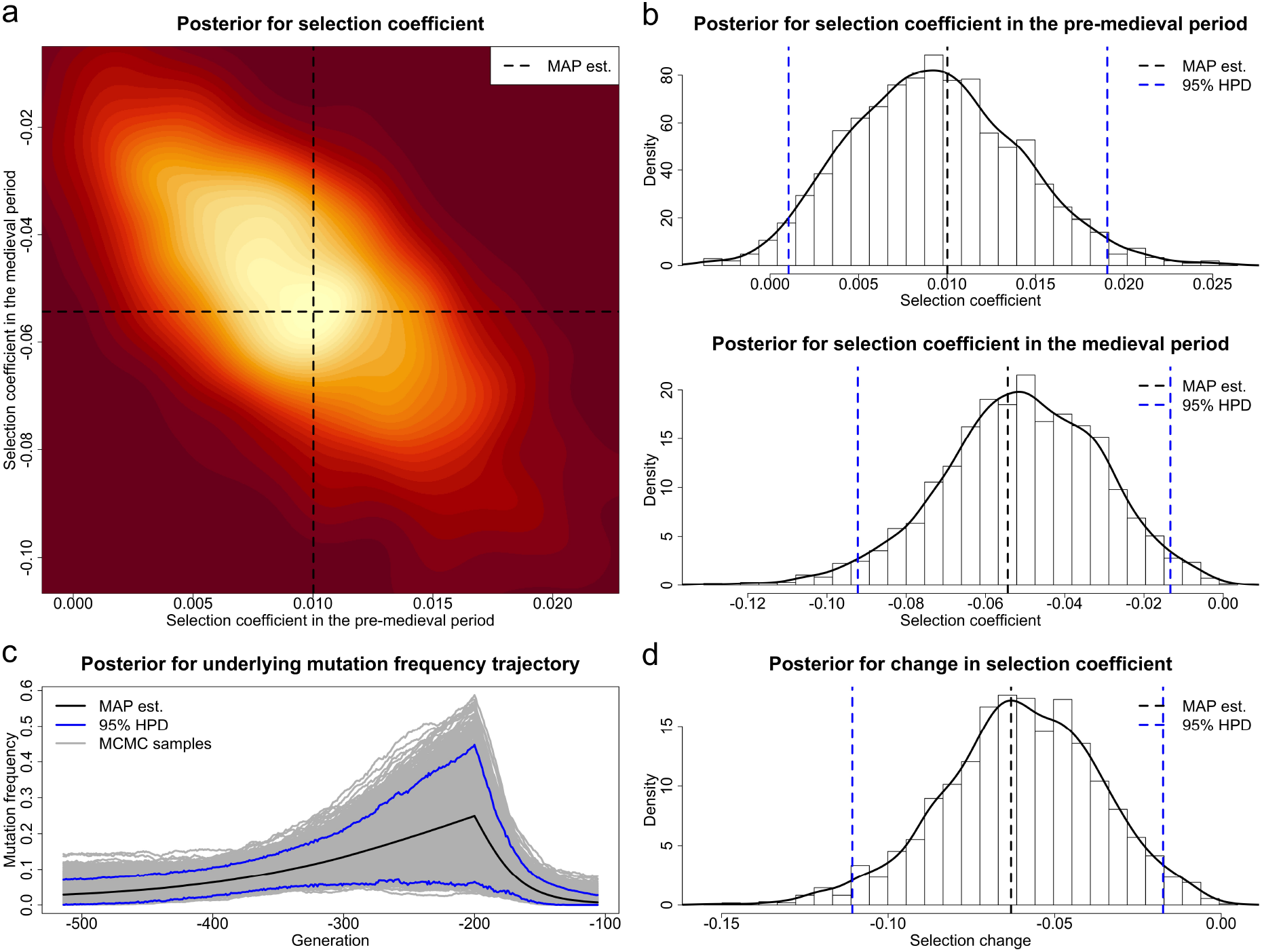
Posteriors for the selection coefficients of the *KIT13* mutation in the pre-medieval and medieval period and the underlying frequency trajectory of the *KIT13* mutation in the population.

#### 3.2.3. Selection of the leopard complex coat pattern

The horse gene *TRPM1* is located on chromosome 1, with mutations coding for the incompletely dominant leopard complex coat (Terry et al., 2004). The resulting posteriors for *TRPM1* are illustrated in Figure 8. Our estimate of the selection coefficient for the *TRPM1* mutation is 0.0090 with 95% HPD interval [−0.0199, 0.0268] before the medieval period and −0.0511 with 95% HPD interval [−0.1299, 0.0437] in the medieval period. Since both 95% HPD intervals contain 0, the evidence to reject the hypothesis that the *TRPM1* mutation experienced selection from modern horse domestication is not sufficient. As shown in Figure 6d, our posterior for the change in the selection coefficient provides weak evidence of a negative change that took place in selection of the *TRPM1* mutation during the Middle Ages (*i*.*e*., the posterior probability for a negative change is is 0.796). However, we can still observe from Figure 6c a gradual increase in the *TRPM1* mutation frequency in pre-medieval horses and a slow decrease in medieval horses.

**Figure 8.**
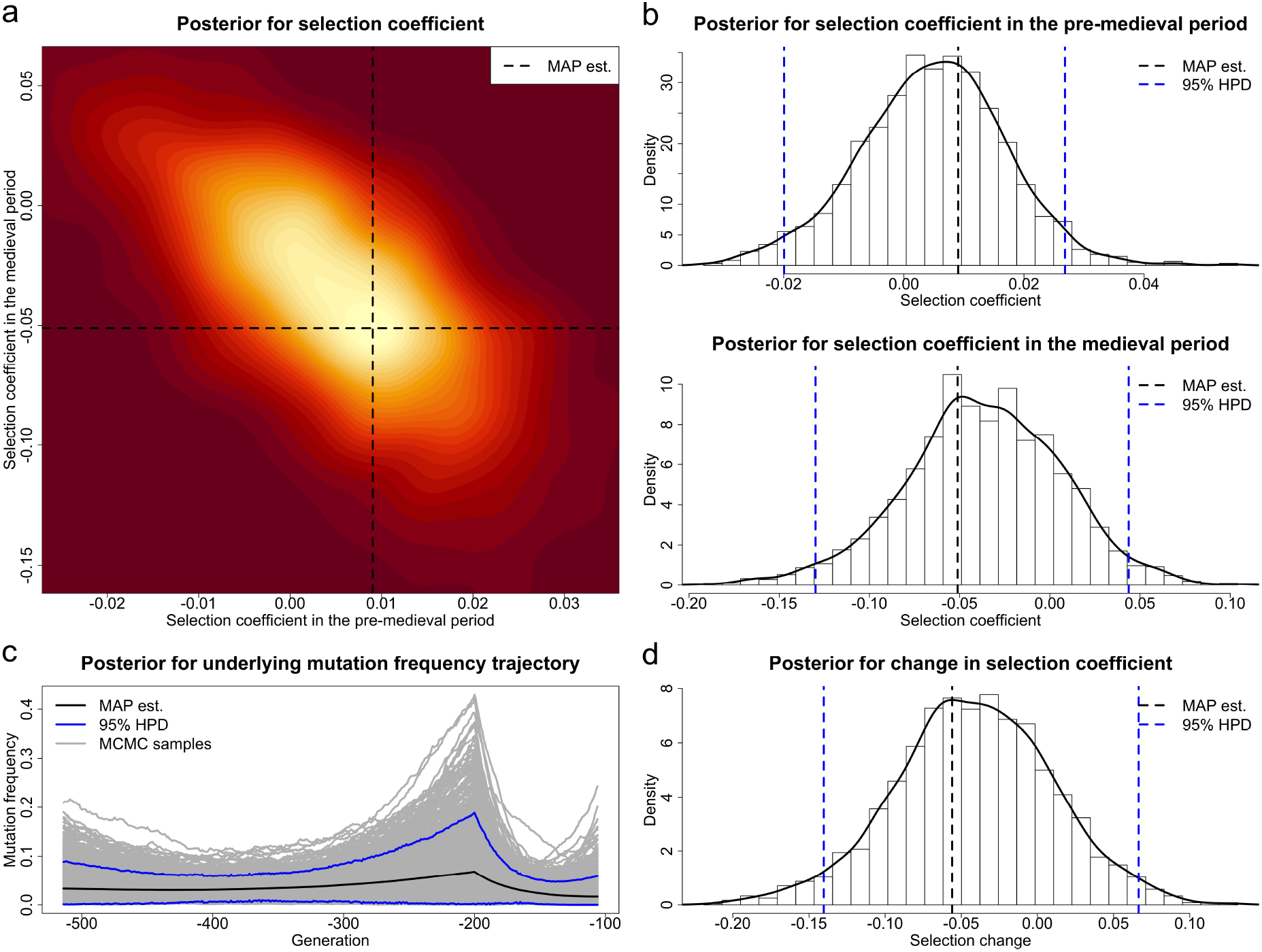
Posteriors for the selection coefficients of the *TRPM1* mutation in the pre-medieval and medieval period and the underlying frequency trajectory of the *TRPM1* mutation in the population.

## 4. Discussion

In this work, we have introduced a novel Bayesian procedure for estimating temporally variable selection intensities from the aDNA data in the form of genotype likelihoods. Our approach is built upon the two-layer HMM framework of He et al. (2023a), which facilitates the incorporation of sample uncertainty resulting from the damage and fragmentation of aDNA molecules. With the PMMH-within-Gibbs algorithm, our approach also permits the reconstruction of underlying population allele frequency trajectories and the integration of temporal information in genotype likelihood corrections. In the MH step, the marginal likelihood is calculated through the particle filter, in which particles are generated by mixing the forward- and backward-in-time simulations of the Wright-Fisher diffusion. Such a setup enables us to jointly estimate the timing when the mutant allele is created and/or lost in the underlying population, therefore fixing the issue encountered in He et al. (2023a,b) that the age of the mutant allele is required. Moreover, our approach provides the flexibility of modelling a time-varying demographic history.

We have tested our method by repeating the same simulation studies as in He et al. (2023a), showing that our approach can deliver accurate selection inferences based on the aDNA data in the form of genotype likelihoods even though samples are sparsely distributed in time with small uneven sizes and in poor qualities. Also, we have illustrated that compared to genotype likelihoods, genotype posteriors produced through our approach, by integrating additional temporal information, allow significant improvement in the process of determining uncertain genotypes of the sample. We therefore expect our procedure to facilitate other population genetics analyses. Although our method has only been evaluated in the case of a bottleneck demographic history and codominance, in principle, the conclusions drawn here hold for other demographic histories and levels of dominance (see Supporting Information, Figures S5 and S6, as well as Tables S8 and S9, for additional simulation studies). Notice that the level of dominance is prespecified in this work, but our method can jointly estimate the dominance parameter as in He et al. (2023a).

We have shown the applicability of our approach with an illustrative case on the aDNA data of ancient horse remains from previous studies of Ludwig et al. (2009), Pruvost et al. (2011) and Wutke et al. (2016), which were genotyped at the pigmentation loci (*e*.*g*., *MC1R, KIT13* and *TRPM1*). Our results show that chestnut, tobiano and leopard complex horses were favoured by selection from modern horse domestication, probably resulting from a need to separate domestic horses from their wild counterparts. Horse coats exhibiting light solid colours or white spotting patterns could facilitate horse husbandry as it was easier to keep track of the horses that were not camouflaged (Fang et al., 2009; Wutke et al., 2016). From our results, we also observe that human preferences for some horse coat colours and patterns changed during the Middle Ages, probably due to shifts in uses, cultures and religions. For example, tobiano and leopard complex horses became negatively selected in the medieval period, which could be explained as resulting in part from a known congenital defect (*i*.*e*., congenital stationary night blindness caused by the *TRPM1* mutation (Bellone et al., 2013)), a lower religious prestige, a reduced need to separate domestic horses from their wild counterparts, pleiotropic disadvantages or novel developments in weaponry during the Middle Ages (Wutke et al., 2016).

As discussed in He et al. (2023a,b), their approaches suffer from the underlying dependency on the allele age, caused by the forward-in-time simulation of the Wright-Fisher diffusion in the particle filter. He et al. (2023a,b) excluded the samples obtained before the first sampling time point that the mutant allele have been found in the sample when the allele age is unavailable, which however can bias the inference of selection. By mixing the forward- and backward-in-time simulations of the Wright-Fisher diffusion in the particle filter, our method can overcome this drawback. To illustrate this, we run the same analysis of the aDNA data from pigmentation loci in ancient horses through our PMMH-within-Gibbs procedure, but in the particle filter, we run the full forward- and backward-in-time simulations of the Wright-Fisher diffusion, respectively. We exclude the samples collected earlier than the time that the mutant allele have been first observed in the sample for the forward-in-time simulation and exclude the samples drawn later than the time that the mutant allele have been last observed in the sample for the backward-in-time simulation, and all other settings are the same as in Section 3.1. The resulting posteriors are shown in Supporting Information, Figures S7–S12, and the resulting estimates are summarised in Supporting Information, Tables S10 and S11, showing that sample exclusion required by our procedure with the full forward-or backward-in-time simulation of the Wright-Fisher diffusion can significantly alter the results of selection inference. For example, with the mix of the forward- and backward-in-time simulations, we find strong evidence supporting that the *KIT13* mutation was positively selected from the domestication of modern horses (*i*.*e*., the posterior probability for positive selection is 0.984) and then became negatively selected during the medieval period (*i*.*e*., the posterior probability for negative selection is 0.999), which matches the archaeological evidence and historical records (see Wutke et al., 2016, and references therein). However, we no longer see enough evidence for positive selection before the medieval period through the full forward-in-time simulation (*i*.*e*., the posterior probability for positive selection is 0.725) and for negative selection during the medieval period through the full backward-in-time simulation (*i*.*e*., the posterior probability for positive selection is 0.876).

It is important to note that incorporating the temporal component could not only improve the inference of selection (see Malaspinas, 2016; Dehasque et al., 2020, and references therein) but also facilitate other population genetic analyses. For example, to our knowledge, all existing methods for genotype likelihood calculations ignore the temporal dimension provided by aDNA data, but in this work, we have demonstrated that genotype likelihoods can be further improved by incorporating temporal information through our PMMH-within-Gibbs procedure, therefore benefiting genotype likelihood based analyses, *e*.*g*., genotype calling and imputation, especially for aDNA studies. This motivates an important future research direction on the methodology development for the analysis of aDNA data by incorporating the temporal component, *e*.*g*., how to integrate the temporal dimension into existing approaches for imputing genotypes (Rubinacci et al., 2021; Ausmees & Nettelblad, 2023) and inferring heterozygosity (Kousathanas et al., 2017; Renaud et al., 2019).

Our results confirm a better or comparable performance of our approach in comparison with the method of He et al. (2023a), especially when samples need to be excluded in their procedure due to the poor knowledge of the allele age. Our simulation studies illustrate that our approach may not perform as well as the method of He et al. (2023a) on extremely poor-quality data. This is because our method simultaneously estimates the genotype of each sample individual, rather than integrating out the uncertainty encountered in genotype likelihoods like He et al. (2023a). However, improving the quality of genotype likelihoods by integrating the temporal dimension is a highly desirable feature in aDNA studies, and moreover, the performance difference in the inference of selection is negligible when the data quality is not too poor. Our procedure therefore is expected to be a convincing alternative for aDNA studies.

One key limitation of our approach (inheriting from He et al. (2023a)) applied to the inference of selection from aDNA data is that all loci are assumed to be independent of each other, which can be easily violated in practice once there exist interactions between genes such as epistasis and linkage. Ignoring these interactions could greatly bias the inference of selection (He et al., 2020a, 2023b). Based on He et al. (2023b), our work can be naturally extended to two genes with epistasis and/or linkage, and the key challenge is how to extend our backward-in-time simulation procedure for the four-dimensional Wright-Fisher diffusion. For multiple interacting genes, such an extension becomes theoretically challenging and computationally prohibitive. By combining with a blockwise updating scheme (*i*.*e*., we partition the population genetic parameters into two blocks, the genome-shared parameter and the locus-specific parameter, and iteratively update one block at a time in our PMMH step), our framework can also be directly extended to jointly estimate the population size like Foll et al. (2015) and Ferrer-Admetlla et al. (2016). Moreover, as in He et al. (2023a,b), the number and timing of the events that might change selection are required to be prespecified in this work, which largely limits its applicability in aDNA studies. A future extension of our method will be to jointly infer selection with its strength and timing of changes, as in Mathieson (2020) and Mathieson & Terhorst (2022). This advance is expected to provide a much deeper insight into the nature of evolution and adaptation with aDNA.

## Acknowledgements

We thank Robert C. Griffiths for helpful comments on the backward-in-time simulation of the Wright-Fisher diffusion. This work was carried out using the computational facilities of the Advanced Computing Research Centre, University of Bristol - http://www.bristol.ac.uk/acrc/.

## Data Accessibility Statement

The authors state that all data necessary for confirming the conclusions of the present work are represented fully within the article. Source code implementing the approach described in this work is available at https://github.com/zhangyi-he/WFM-1L-DiffusApprox-PMMHwGibbs/.

## Author Contributions

F.Y. and Z.H. designed the project and developed the approach; W.L. and Z.H. implemented the approach; W.L. and X.D. analysed the data under the supervision of M.B., F.Y. and Z.H.; Z.H. wrote the manuscript; W.L., X.D., M.B. and F.Y. reviewed the manuscript.

