## Supporting Information for "Estimating temporally variable selection intensity from ancient DNA data II"

### File S1. Additional results for analysis of simulated data

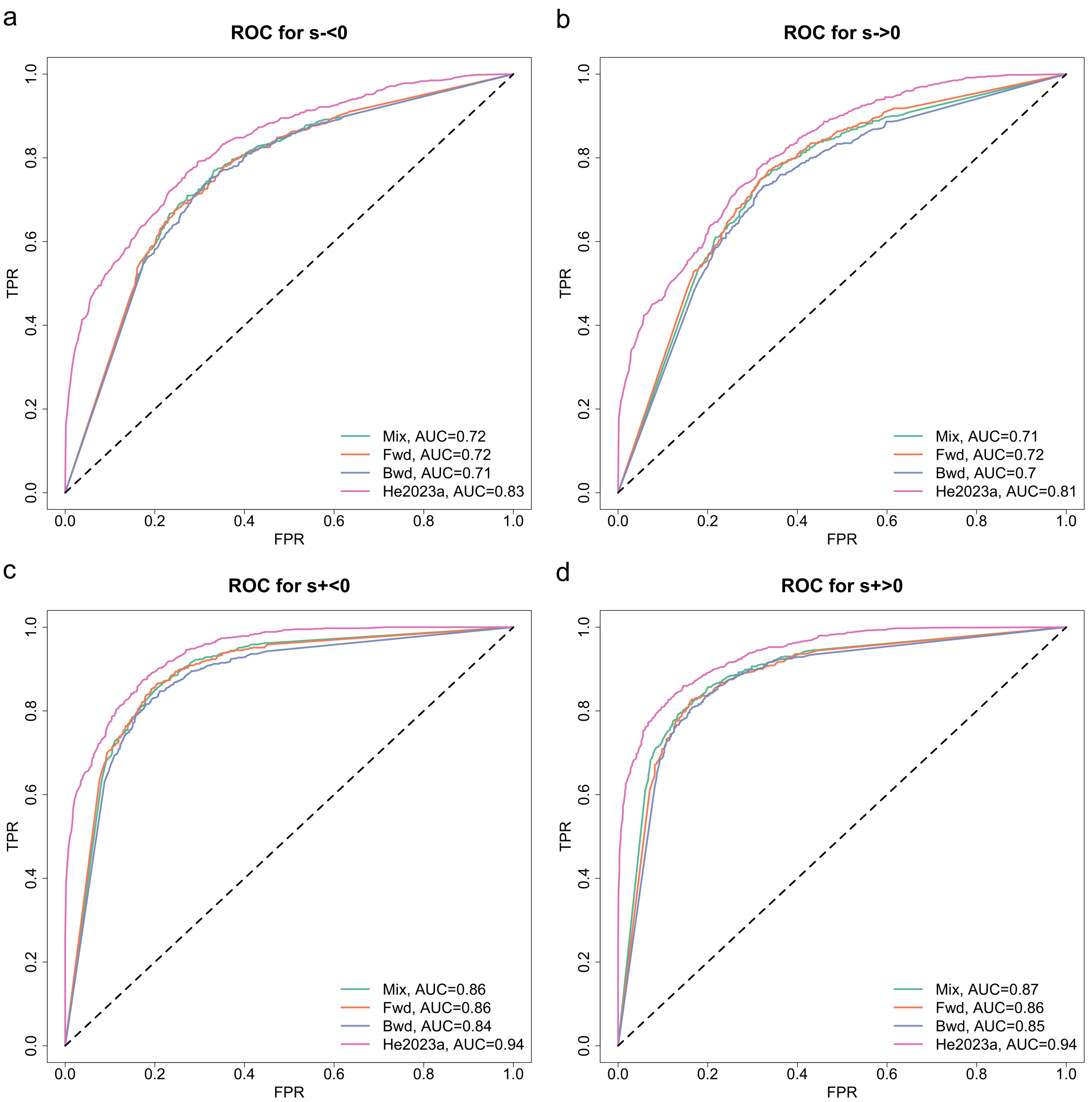


(a) On average simulated datasets comprise of 30.86% genotype missing calls with an SD of 3.27% and 9.60% genotype calling errors with an SD of 2.04% ($\phi = 0.75$ and $\psi = 0.5$).

Figure S1: ROC curves for detecting selection signatures across different selection scenarios and data qualities.

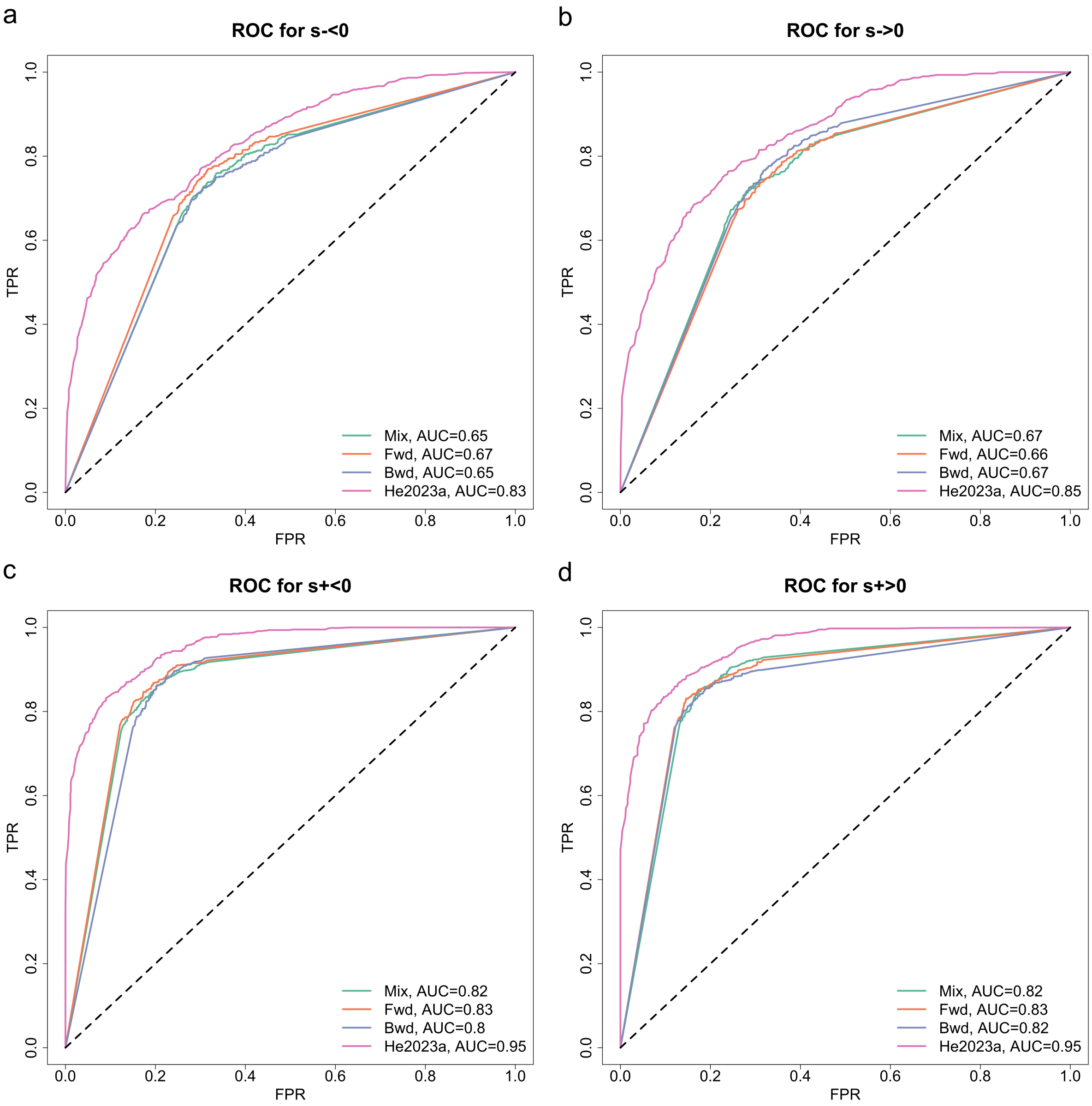


(b) On average simulated datasets comprise of 45.44% genotype missing calls with an SD of 3.35% and 3.83% genotype calling errors with an SD of 1.36% ($\phi = 0.75$ and $\psi = 1.0$).

Figure S1: ROC curves for detecting selection signatures across different selection scenarios and data qualities, continued.

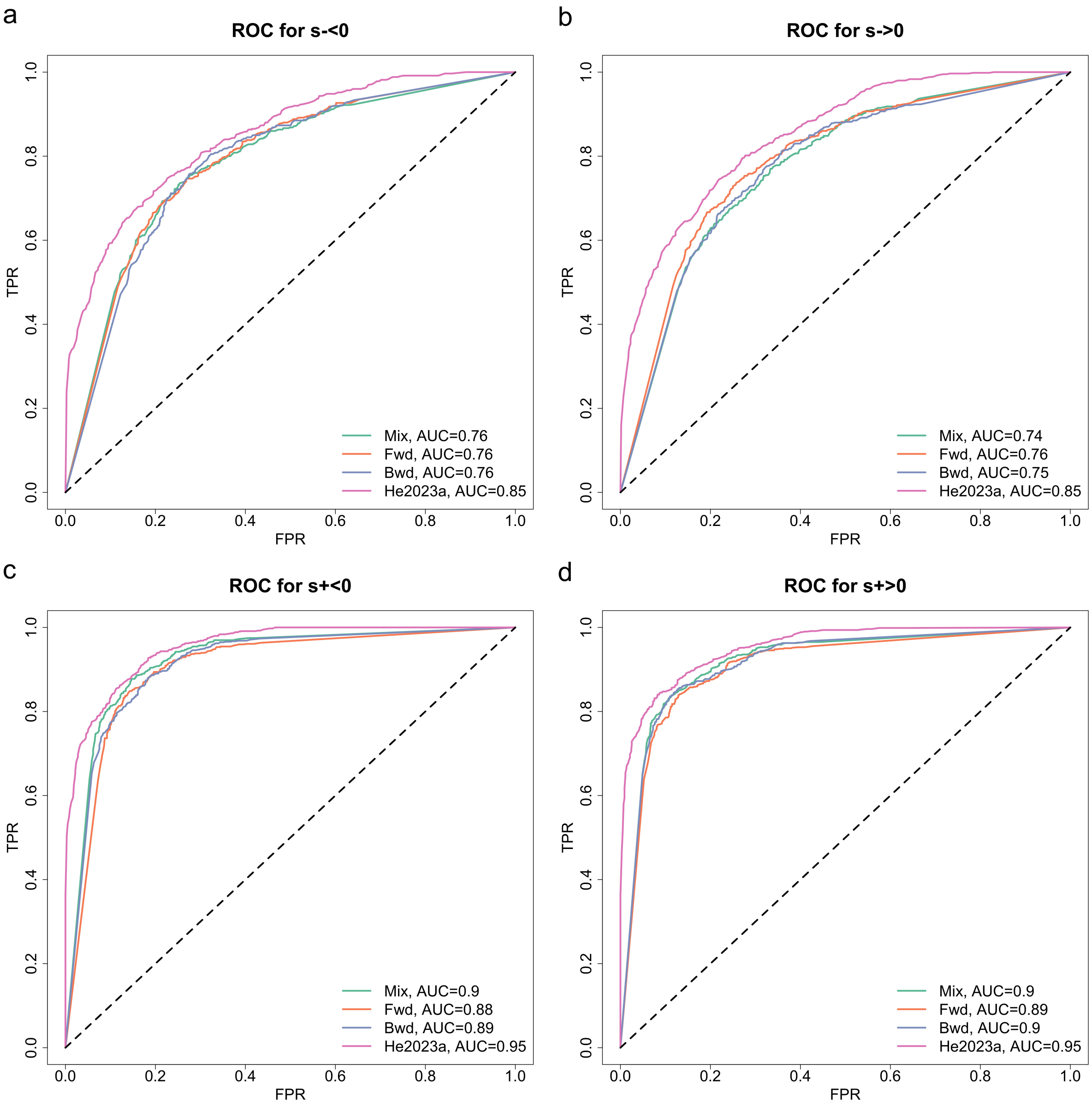


(c) On average simulated datasets comprise of 30.05% genotype missing calls with an SD of 3.18% and 1.90% genotype calling errors with an SD of 0.95% ($\phi = 0.85$ and $\psi = 1.0$).

Figure S1: ROC curves for detecting selection signatures across different selection scenarios and data qualities, continued.

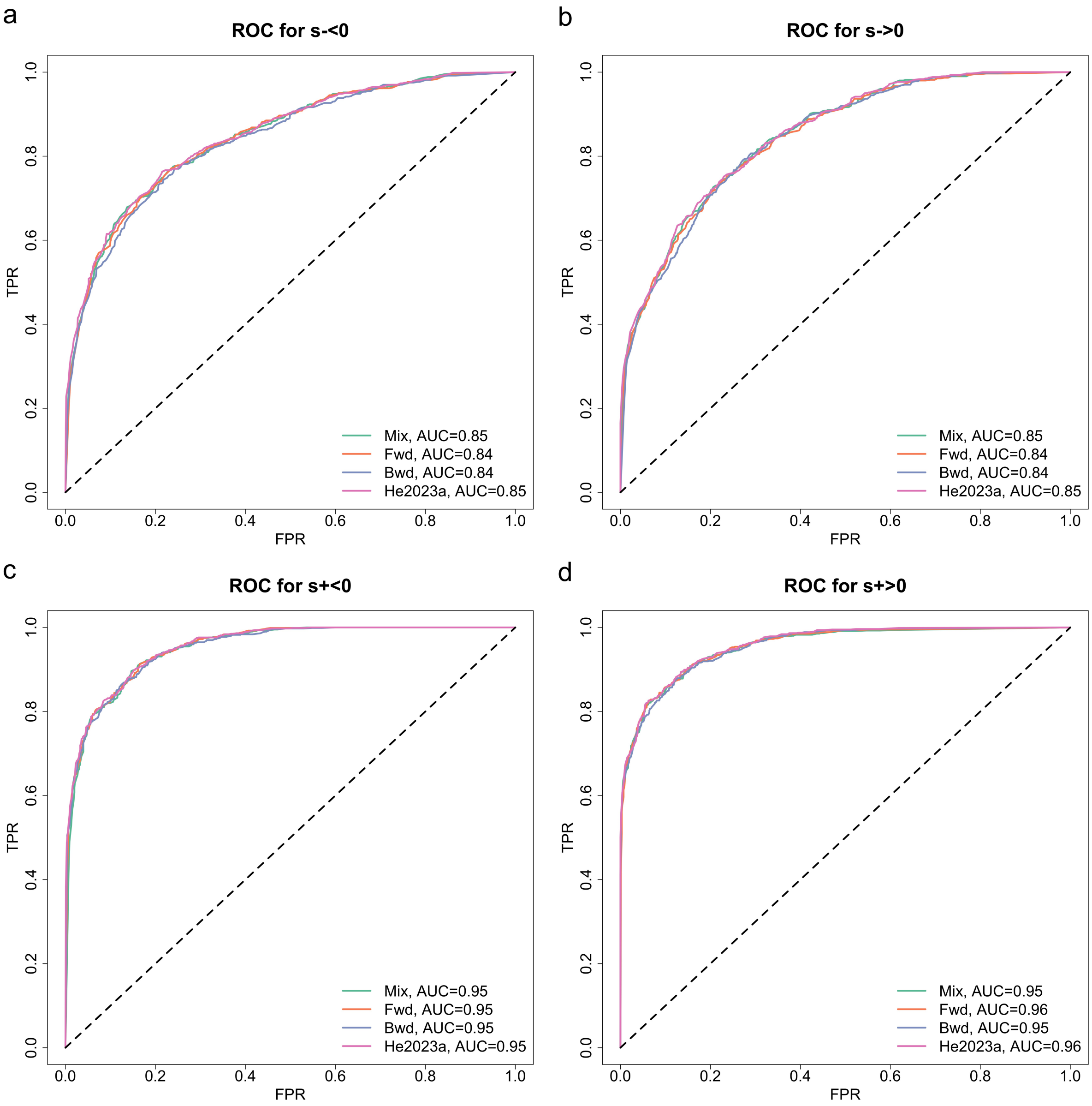


(d) On average simulated datasets comprise of 7.48% genotype missing calls with an SD of 1.86% and 1.64% genotype calling errors with an SD of 0.86% ($\phi = 0.95$ and $\psi = 0.5$).

Figure S1: ROC curves for detecting selection signatures across different selection scenarios and data qualities, continued.

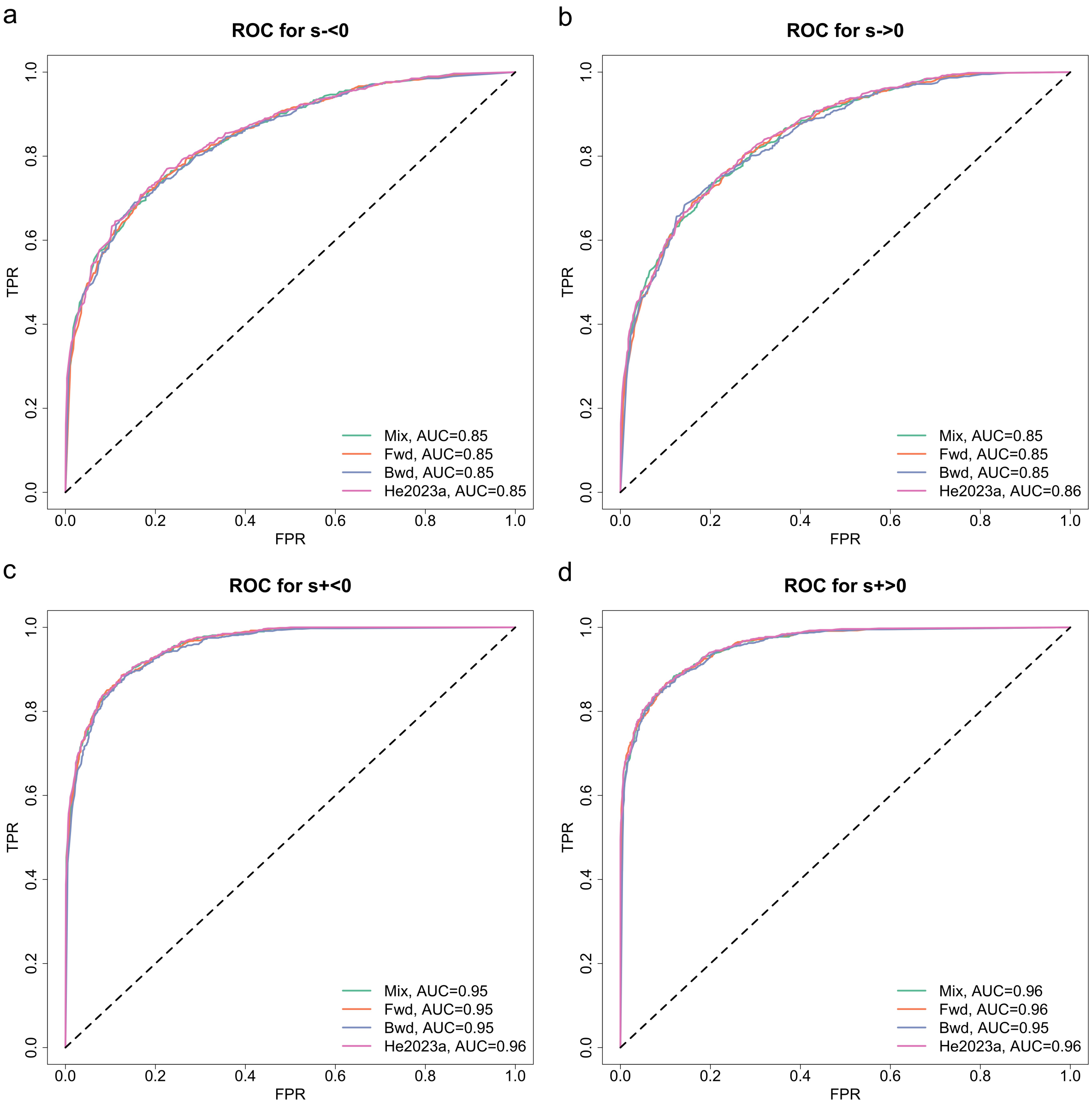


(e) On average simulated datasets comprise of 10.96% genotype missing calls with an SD of 2.08% and 0.53% genotype calling errors with an SD of 0.50% ($\phi = 0.95$ and $\psi = 1.0$).

Figure S1: ROC curves for detecting selection signatures across different selection scenarios and data qualities, continued.

| Selection coefficient | Selection scenario | Mix | | Fwd | | Bwd | | He2023a | |
|---|---|---|---|---|---|---|---|---|---|
| | | Bias | RMSE | Bias | RMSE | Bias | RMSE | Bias | RMSE |
| $s^-$ | $s^- < 0$, $s^+ < s^-$ | 0.00066 | 0.00421 | 0.00043 | 0.00406 | 0.00053 | 0.00467 | 0.00076 | 0.00332 |
| | $s^- < 0$, $s^+ = s^-$ | 0.00041 | 0.00387 | 0.00049 | 0.00394 | 0.00028 | 0.00454 | 0.00065 | 0.00332 |
| | $s^- < 0$, $s^+ > s^-$ | 0.00034 | 0.00381 | 0.00063 | 0.00413 | 0.00057 | 0.00407 | 0.00091 | 0.00332 |
| | $s^- = 0$, $s^+ < s^-$ | 0.00025 | 0.00413 | 0.00059 | 0.00397 | 0.00028 | 0.00419 | 0.00036 | 0.00327 |
| | $s^- = 0$, $s^+ = s^-$ | -0.00001 | 0.00366 | -0.00024 | 0.00396 | -0.00013 | 0.00413 | -0.00026 | 0.00292 |
| | $s^- = 0$, $s^+ > s^-$ | -0.00028 | 0.00364 | -0.00016 | 0.00448 | -0.00002 | 0.00437 | -0.00001 | 0.00338 |
| | $s^- > 0$, $s^+ < s^-$ | -0.00038 | 0.00447 | -0.00056 | 0.00430 | -0.00013 | 0.00459 | -0.00076 | 0.00345 |
| | $s^- > 0$, $s^+ = s^-$ | -0.00083 | 0.00410 | -0.00073 | 0.00426 | -0.00063 | 0.00435 | -0.00094 | 0.00335 |
| | $s^- > 0$, $s^+ > s^-$ | -0.00098 | 0.00429 | -0.00097 | 0.00432 | -0.00105 | 0.00444 | -0.00097 | 0.00333 |
| $s^+$ | $s^- < 0$, $s^+ < s^-$ | 0.00047 | 0.00322 | 0.00060 | 0.00349 | 0.00030 | 0.00372 | 0.00069 | 0.00264 |
| | $s^- < 0$, $s^+ = s^-$ | 0.00019 | 0.00330 | 0.00041 | 0.00348 | 0.00025 | 0.00381 | 0.00053 | 0.00267 |
| | $s^- < 0$, $s^+ > s^-$ | -0.00077 | 0.00337 | -0.00062 | 0.00343 | -0.00069 | 0.00359 | -0.00087 | 0.00286 |
| | $s^- = 0$, $s^+ < s^-$ | 0.00033 | 0.00326 | 0.00003 | 0.00328 | 0.00042 | 0.00376 | 0.00041 | 0.00268 |
| | $s^- = 0$, $s^+ = s^-$ | 0.00004 | 0.00304 | -0.00008 | 0.00298 | -0.00012 | 0.00322 | 0.00000 | 0.00249 |
| | $s^- = 0$, $s^+ > s^-$ | -0.00095 | 0.00339 | -0.00100 | 0.00344 | -0.00088 | 0.00355 | -0.00106 | 0.00301 |
| | $s^- > 0$, $s^+ < s^-$ | 0.00049 | 0.00381 | 0.00056 | 0.00335 | 0.00070 | 0.00387 | 0.00082 | 0.00307 |
| | $s^- > 0$, $s^+ = s^-$ | -0.00018 | 0.00356 | -0.00030 | 0.00334 | -0.00014 | 0.00341 | -0.00027 | 0.00268 |
| | $s^- > 0$, $s^+ > s^-$ | -0.00047 | 0.00335 | -0.00024 | 0.00321 | -0.00027 | 0.00372 | -0.00060 | 0.00283 |

Table S1: Mean bias and RMSE in the selection coefficient estimates across different selection scenarios, corresponding to Figure 3.

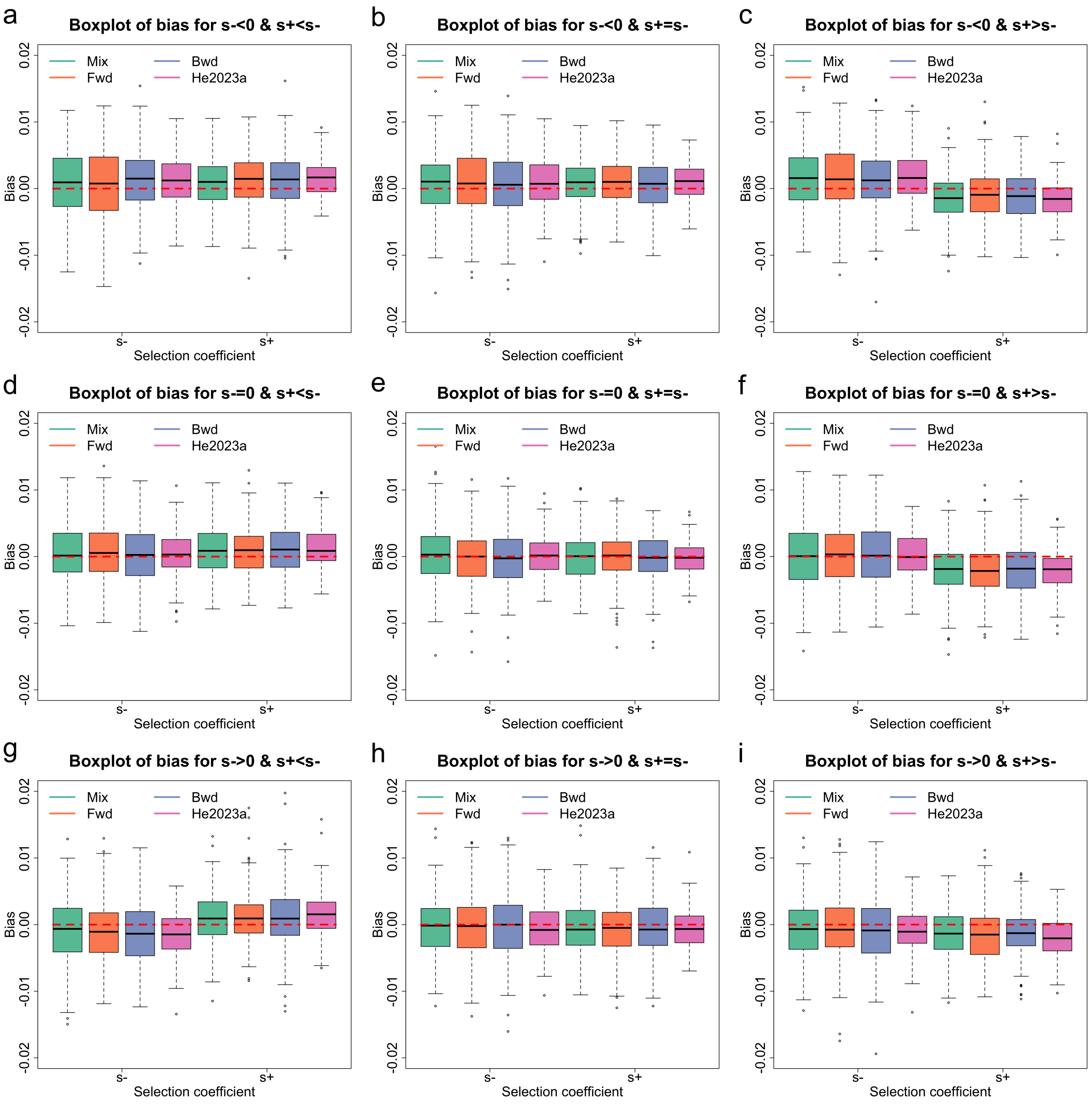


(a) On average simulated datasets comprise of 30.86% genotype missing calls with an SD of 3.27% and 9.60% genotype calling errors with an SD of 2.04% ($\phi = 0.75$ and $\psi = 0.5$).

Figure S2: Boxplots for the bias of the selection coefficient estimates across different selection scenarios and data qualities.

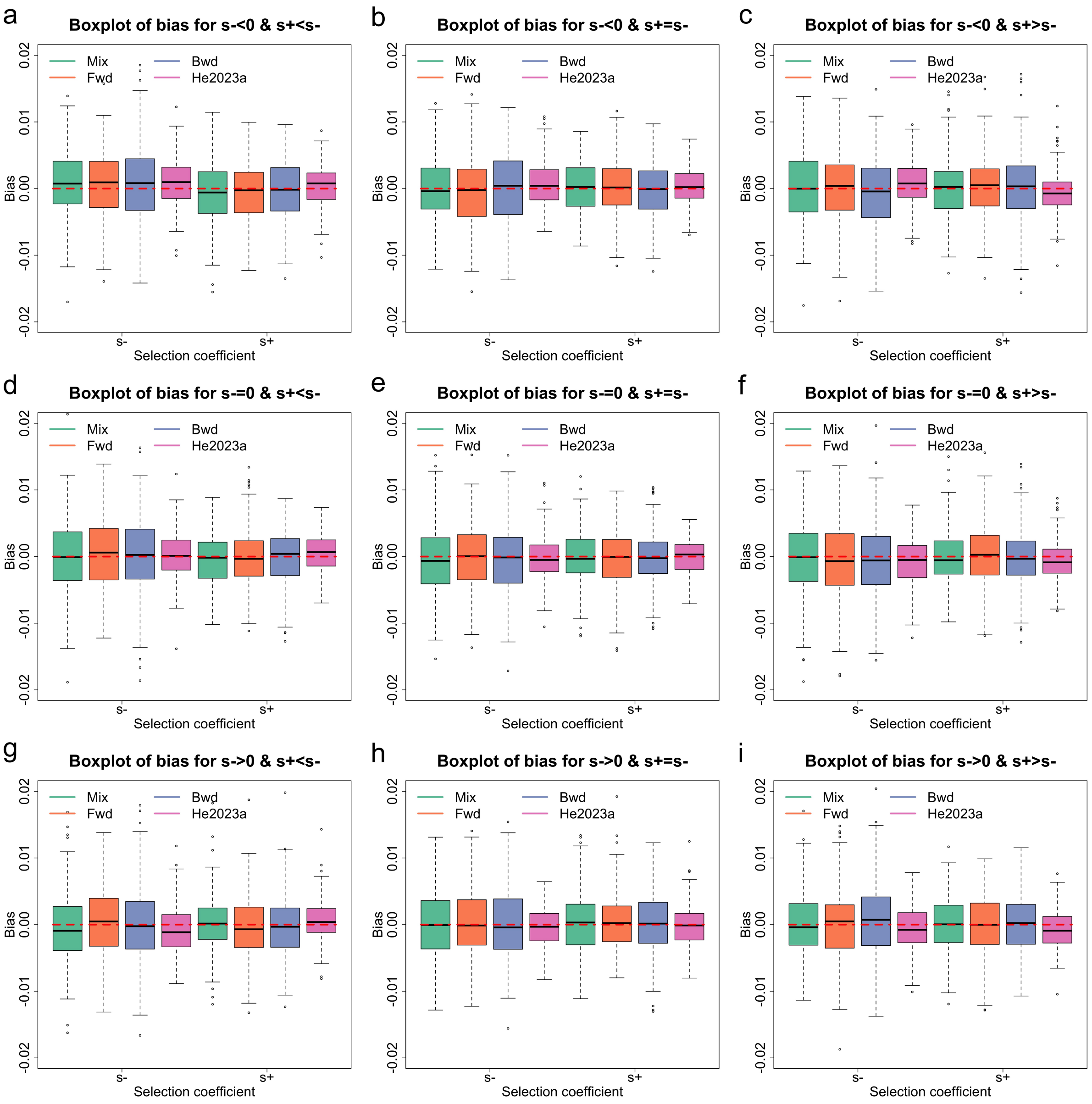


(b) On average simulated datasets comprise of 45.44% genotype missing calls with an SD of 3.35% and 3.83% genotype calling errors with an SD of 1.36% ($\phi = 0.75$ and $\psi = 1.0$).

Figure S2: Boxplots for the bias of the selection coefficient estimates across different selection scenarios and data qualities, continued.

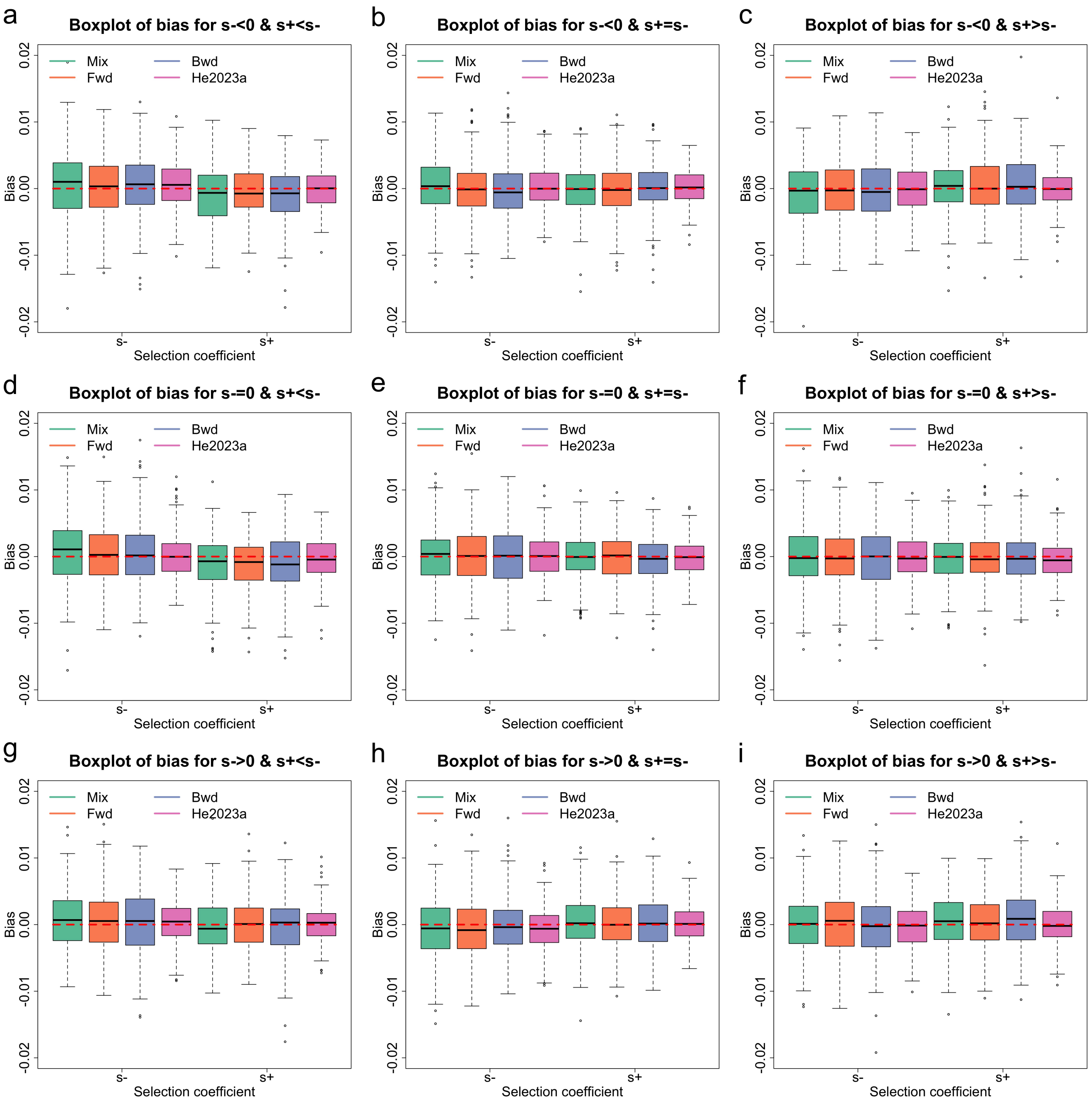


(c) On average simulated datasets comprise of 30.05% genotype missing calls with an SD of 3.18% and 1.90% genotype calling errors with an SD of 0.95% ($\phi = 0.85$ and $\psi = 1.0$).

Figure S2: Boxplots for the bias of the selection coefficient estimates across different selection scenarios and data qualities, continued.

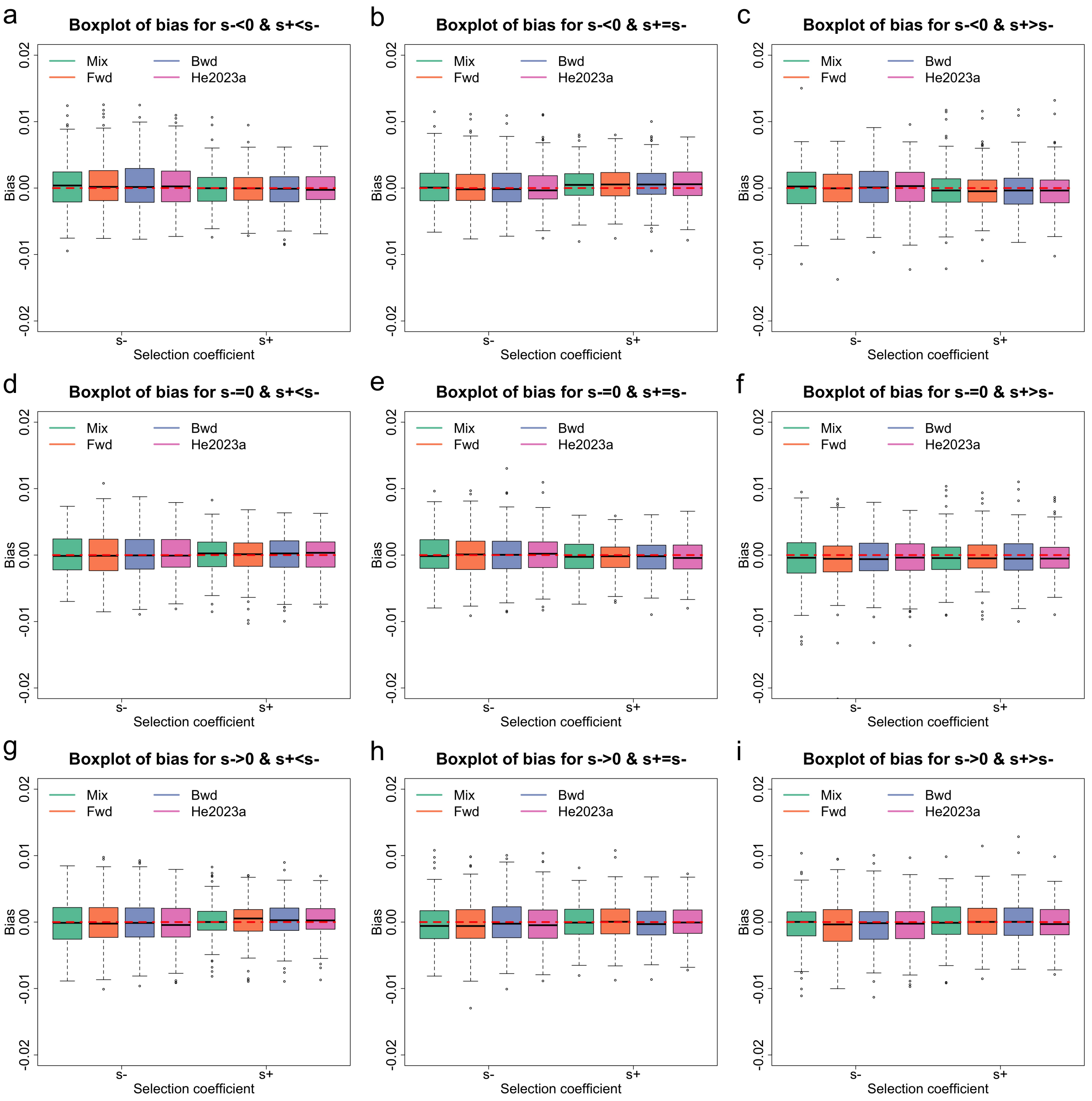


(d) On average simulated datasets comprise of 7.48% genotype missing calls with an SD of 1.86% and 1.64% genotype calling errors with an SD of 0.86% ($\phi = 0.95$ and $\psi = 0.5$).

Figure S2: Boxplots for the bias of the selection coefficient estimates across different selection scenarios and data qualities, continued.

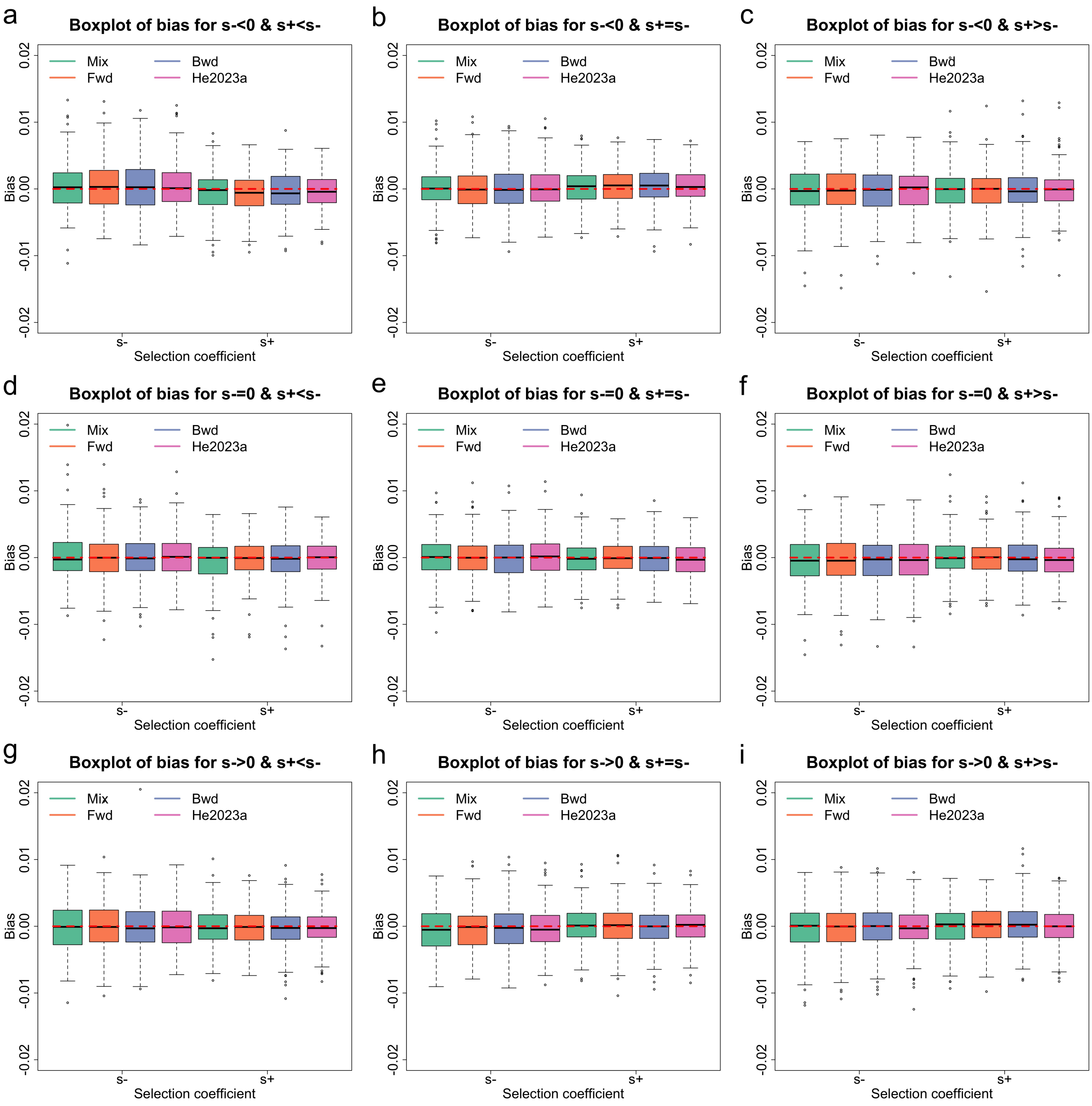


(e) On average simulated datasets comprise of 10.96% genotype missing calls with an SD of 2.08% and 0.53% genotype calling errors with an SD of 0.50% ($\phi = 0.95$ and $\psi = 1.0$).

Figure S2: Boxplots for the bias of the selection coefficient estimates across different selection scenarios and data qualities, continued.

| Selection coefficient | Selection scenario | Mix | | Fwd | | Bwd | | He2023a | |
|---|---|---|---|---|---|---|---|---|---|
| | | Bias | RMSE | Bias | RMSE | Bias | RMSE | Bias | RMSE |
| | $s^- < 0,\ s^+ < s^-$ | 0.00087 | 0.00496 | 0.00061 | 0.00559 | 0.00121 | 0.00476 | 0.00125 | 0.00387 |
| | $s^- < 0,\ s^+ = s^-$ | 0.00086 | 0.00454 | 0.00067 | 0.00481 | 0.00052 | 0.00482 | 0.00085 | 0.00354 |
| | $s^- < 0,\ s^+ > s^-$ | 0.00151 | 0.00477 | 0.00157 | 0.00495 | 0.00131 | 0.00476 | 0.00177 | 0.00381 |
| | $s^- = 0,\ s^+ < s^-$ | 0.00039 | 0.00433 | 0.00051 | 0.00436 | 0.00029 | 0.00440 | 0.00045 | 0.00328 |
| $s^-$ | $s^- = 0,\ s^+ = s^-$ | 0.00032 | 0.00437 | -0.00018 | 0.00404 | -0.00018 | 0.00413 | 0.00000 | 0.00300 |
| | $s^- = 0,\ s^+ > s^-$ | 0.00005 | 0.00472 | 0.00021 | 0.00445 | 0.00036 | 0.00485 | 0.00012 | 0.00320 |
| | $s^- > 0,\ s^+ < s^-$ | -0.00072 | 0.00478 | -0.00096 | 0.00472 | -0.00116 | 0.00486 | -0.00159 | 0.00368 |
| | $s^- > 0,\ s^+ = s^-$ | -0.00026 | 0.00448 | -0.00020 | 0.00461 | -0.00026 | 0.00479 | -0.00068 | 0.00343 |
| | $s^- > 0,\ s^+ > s^-$ | -0.00071 | 0.00456 | -0.00066 | 0.00494 | -0.00094 | 0.00486 | -0.00085 | 0.00334 |
| | $s^- < 0,\ s^+ < s^-$ | 0.00088 | 0.00386 | 0.00128 | 0.00407 | 0.00115 | 0.00439 | 0.00147 | 0.00307 |
| | $s^- < 0,\ s^+ = s^-$ | 0.00068 | 0.00372 | 0.00092 | 0.00380 | 0.00065 | 0.00404 | 0.00101 | 0.00284 |
| | $s^- < 0,\ s^+ > s^-$ | -0.00149 | 0.00384 | -0.00097 | 0.00386 | -0.00112 | 0.00387 | -0.00163 | 0.00322 |
| | $s^- = 0,\ s^+ < s^-$ | 0.00107 | 0.00393 | 0.00090 | 0.00374 | 0.00108 | 0.00425 | 0.00127 | 0.00311 |
| $s^+$ | $s^- = 0,\ s^+ = s^-$ | -0.00024 | 0.00364 | -0.00011 | 0.00350 | -0.00022 | 0.00363 | -0.00025 | 0.00241 |
| | $s^- = 0,\ s^+ > s^-$ | -0.00198 | 0.00424 | -0.00202 | 0.00437 | -0.00178 | 0.00442 | -0.00203 | 0.00345 |
| | $s^- > 0,\ s^+ < s^-$ | 0.00099 | 0.00431 | 0.00106 | 0.00402 | 0.00108 | 0.00456 | 0.00147 | 0.00351 |
| | $s^- > 0,\ s^+ = s^-$ | -0.00044 | 0.00410 | -0.00072 | 0.00393 | -0.00026 | 0.00414 | -0.00061 | 0.00295 |
| | $s^- > 0,\ s^+ > s^-$ | -0.00152 | 0.00399 | -0.00150 | 0.00431 | -0.00121 | 0.00368 | -0.00206 | 0.00359 |

(a) On average simulated datasets comprise of 30.86% genotype missing calls with an SD of 3.27% and 9.60% genotype calling errors with an SD of 2.04% ($\phi = 0.75$ and $\psi = 0.5$), corresponding to Figure S2a.

Table S2: Mean bias and RMSE in the selection coefficient estimates across different selection scenarios and data qualities, corresponding to Figure S2.

| Selection coefficient | Selection scenario | Mix | | Fwd | | Bwd | | He2023a | |
|---|---|---|---|---|---|---|---|---|---|
| | | Bias | RMSE | Bias | RMSE | Bias | RMSE | Bias | RMSE |
| $s^-$ | $s^- < 0$, $s^+ < s^-$ | 0.00076 | 0.00509 | 0.00070 | 0.00525 | 0.00055 | 0.00612 | 0.00083 | 0.00374 |
| | $s^- < 0$, $s^+ = s^-$ | -0.00012 | 0.00466 | -0.00036 | 0.00529 | 0.00000 | 0.00529 | 0.00069 | 0.00343 |
| | $s^- < 0$, $s^+ > s^-$ | 0.00020 | 0.00552 | 0.00002 | 0.00500 | -0.00058 | 0.00569 | 0.00072 | 0.00356 |
| | $s^- = 0$, $s^+ < s^-$ | 0.00011 | 0.00537 | 0.00033 | 0.00519 | 0.00023 | 0.00579 | 0.00021 | 0.00358 |
| | $s^- = 0$, $s^+ = s^-$ | -0.00031 | 0.00539 | -0.00005 | 0.00518 | -0.00049 | 0.00529 | -0.00035 | 0.00336 |
| | $s^- = 0$, $s^+ > s^-$ | -0.00048 | 0.00555 | -0.00075 | 0.00574 | -0.00065 | 0.00545 | -0.00068 | 0.00370 |
| | $s^- > 0$, $s^+ < s^-$ | -0.00045 | 0.00538 | 0.00040 | 0.00513 | 0.00005 | 0.00546 | -0.00087 | 0.00367 |
| | $s^- > 0$, $s^+ = s^-$ | 0.00014 | 0.00507 | -0.00001 | 0.00495 | -0.00002 | 0.00527 | -0.00042 | 0.00307 |
| | $s^- > 0$, $s^+ > s^-$ | -0.00013 | 0.00488 | -0.00007 | 0.00516 | 0.00058 | 0.00540 | -0.00064 | 0.00359 |
| $s^+$ | $s^- < 0$, $s^+ < s^-$ | -0.00051 | 0.00464 | -0.00065 | 0.00460 | -0.00027 | 0.00443 | 0.00047 | 0.00305 |
| | $s^- < 0$, $s^+ = s^-$ | 0.00023 | 0.00398 | 0.00020 | 0.00403 | -0.00015 | 0.00418 | 0.00034 | 0.00273 |
| | $s^- < 0$, $s^+ > s^-$ | -0.00012 | 0.00467 | 0.00043 | 0.00451 | 0.00018 | 0.00523 | -0.00071 | 0.00316 |
| | $s^- = 0$, $s^+ < s^-$ | -0.00035 | 0.00391 | 0.00003 | 0.00412 | -0.00021 | 0.00445 | 0.00069 | 0.00288 |
| | $s^- = 0$, $s^+ = s^-$ | -0.00008 | 0.00405 | -0.00018 | 0.00414 | -0.00008 | 0.00401 | -0.00002 | 0.00283 |
| | $s^- = 0$, $s^+ > s^-$ | -0.00024 | 0.00426 | 0.00026 | 0.00463 | -0.00010 | 0.00469 | -0.00066 | 0.00301 |
| | $s^- > 0$, $s^+ < s^-$ | 0.00013 | 0.00431 | -0.00060 | 0.00449 | -0.00030 | 0.00475 | 0.00066 | 0.00307 |
| | $s^- > 0$, $s^+ = s^-$ | 0.00039 | 0.00460 | 0.00050 | 0.00435 | 0.00035 | 0.00480 | -0.00014 | 0.00298 |
| | $s^- > 0$, $s^+ > s^-$ | 0.00028 | 0.00424 | -0.00015 | 0.00433 | 0.00031 | 0.00430 | -0.00078 | 0.00292 |

(a) On average simulated datasets comprise of 45.44% genotype missing calls with an SD of 3.35% and 3.83% genotype calling errors with an SD of 1.36% ($\phi = 0.75$ and $\psi = 1.0$), corresponding to Figure S2b.

Table S2: Mean bias and RMSE in the selection coefficient estimates across different selection scenarios and data qualities, corresponding to Figure S2, continued.

| Selection coefficient | Selection scenario | Mix | | Fwd | | Bwd | | He2023a | |
|---|---|---|---|---|---|---|---|---|---|
| | | Bias | RMSE | Bias | RMSE | Bias | RMSE | Bias | RMSE |
| $s^-$ | $s^- < 0$, $s^+ < s^-$ | 0.00067 | 0.00507 | 0.00038 | 0.00445 | 0.00053 | 0.00474 | 0.00058 | 0.00348 |
| | $s^- < 0$, $s^+ = s^-$ | 0.00026 | 0.00452 | -0.00027 | 0.00436 | -0.00024 | 0.00432 | 0.00030 | 0.00299 |
| | $s^- < 0$, $s^+ > s^-$ | -0.00067 | 0.00455 | -0.00032 | 0.00429 | -0.00025 | 0.00447 | 0.00005 | 0.00331 |
| | $s^- = 0$, $s^+ < s^-$ | 0.00068 | 0.00488 | 0.00016 | 0.00437 | 0.00047 | 0.00500 | 0.00010 | 0.00350 |
| | $s^- = 0$, $s^+ = s^-$ | 0.00023 | 0.00423 | 0.00010 | 0.00439 | 0.00011 | 0.00454 | 0.00000 | 0.00325 |
| | $s^- = 0$, $s^+ > s^-$ | -0.00017 | 0.00470 | -0.00028 | 0.00452 | -0.00016 | 0.00518 | -0.00020 | 0.00344 |
| | $s^- > 0$, $s^+ < s^-$ | 0.00074 | 0.00466 | 0.00051 | 0.00480 | 0.00027 | 0.00473 | 0.00015 | 0.00320 |
| | $s^- > 0$, $s^+ = s^-$ | -0.00057 | 0.00462 | -0.00070 | 0.00436 | -0.00026 | 0.00423 | -0.00058 | 0.00323 |
| | $s^- > 0$, $s^+ > s^-$ | -0.00008 | 0.00437 | 0.00014 | 0.00483 | -0.00025 | 0.00533 | -0.00026 | 0.00331 |
| $s^+$ | $s^- < 0$, $s^+ < s^-$ | -0.00099 | 0.00408 | -0.00047 | 0.00376 | -0.00087 | 0.00412 | -0.00006 | 0.00280 |
| | $s^- < 0$, $s^+ = s^-$ | -0.00007 | 0.00362 | -0.00027 | 0.00382 | 0.00011 | 0.00368 | 0.00015 | 0.00264 |
| | $s^- < 0$, $s^+ > s^-$ | 0.00046 | 0.00431 | 0.00043 | 0.00417 | 0.00055 | 0.00453 | -0.00002 | 0.00288 |
| | $s^- = 0$, $s^+ < s^-$ | -0.00100 | 0.00436 | -0.00111 | 0.00394 | -0.00092 | 0.00433 | -0.00031 | 0.00309 |
| | $s^- = 0$, $s^+ = s^-$ | -0.00010 | 0.00359 | -0.00011 | 0.00362 | -0.00041 | 0.00368 | -0.00009 | 0.00271 |
| | $s^- = 0$, $s^+ > s^-$ | -0.00018 | 0.00373 | -0.00021 | 0.00403 | -0.00016 | 0.00405 | -0.00049 | 0.00300 |
| | $s^- > 0$, $s^+ < s^-$ | -0.00021 | 0.00396 | 0.00004 | 0.00403 | -0.00038 | 0.00412 | 0.00019 | 0.00280 |
| | $s^- > 0$, $s^+ = s^-$ | 0.00026 | 0.00388 | 0.00022 | 0.00392 | 0.00026 | 0.00392 | 0.00002 | 0.00274 |
| | $s^- > 0$, $s^+ > s^-$ | 0.00052 | 0.00407 | 0.00019 | 0.00374 | 0.00074 | 0.00465 | -0.00001 | 0.00303 |

(b) On average simulated datasets comprise of 30.05% genotype missing calls with an SD of 3.18% and 1.90% genotype calling errors with an SD of 0.95% ($\phi = 0.85$ and $\psi = 1.0$), corresponding to Figure S2c.

Table S2: Mean bias and RMSE in the selection coefficient estimates across different selection scenarios and data qualities, corresponding to Figure S2, continued.

| Selection coefficient | Selection scenario | Mix | | Fwd | | Bwd | | He2023a | |
|---|---|---|---|---|---|---|---|---|---|
| | | Bias | RMSE | Bias | RMSE | Bias | RMSE | Bias | RMSE |
| | $s^- < 0,\ s^+ < s^-$ | 0.00041 | 0.00365 | 0.00058 | 0.00377 | 0.00058 | 0.00373 | 0.00062 | 0.00356 |
| | $s^- < 0,\ s^+ = s^-$ | 0.00024 | 0.00313 | 0.00017 | 0.00317 | 0.00008 | 0.00322 | 0.00017 | 0.00309 |
| | $s^- < 0,\ s^+ > s^-$ | 0.00008 | 0.00345 | -0.00010 | 0.00313 | 0.00010 | 0.00331 | 0.00009 | 0.00323 |
| | $s^- = 0,\ s^+ < s^-$ | -0.00004 | 0.00317 | 0.00002 | 0.00332 | 0.00000 | 0.00330 | 0.00003 | 0.00315 |
| $s^-$ | $s^- = 0,\ s^+ = s^-$ | 0.00000 | 0.00305 | 0.00005 | 0.00324 | 0.00017 | 0.00322 | 0.00013 | 0.00308 |
| | $s^- = 0,\ s^+ > s^-$ | -0.00005 | 0.00364 | -0.00053 | 0.00372 | -0.00046 | 0.00350 | -0.00047 | 0.00323 |
| | $s^- > 0,\ s^+ < s^-$ | -0.00012 | 0.00338 | -0.00016 | 0.00353 | -0.00014 | 0.00361 | -0.00030 | 0.00324 |
| | $s^- > 0,\ s^+ = s^-$ | -0.00039 | 0.00340 | -0.00034 | 0.00341 | -0.00007 | 0.00335 | -0.00035 | 0.00325 |
| | $s^- > 0,\ s^+ > s^-$ | -0.00030 | 0.00316 | -0.00047 | 0.00341 | -0.00038 | 0.00335 | -0.00034 | 0.00310 |
| | $s^- < 0,\ s^+ < s^-$ | -0.00016 | 0.00283 | -0.00009 | 0.00275 | -0.00021 | 0.00285 | -0.00009 | 0.00258 |
| | $s^- < 0,\ s^+ = s^-$ | 0.00051 | 0.00262 | 0.00056 | 0.00271 | 0.00063 | 0.00285 | 0.00064 | 0.00266 |
| | $s^- < 0,\ s^+ > s^-$ | -0.00035 | 0.00331 | -0.00022 | 0.00308 | -0.00025 | 0.00311 | -0.00035 | 0.00302 |
| | $s^- = 0,\ s^+ < s^-$ | 0.00008 | 0.00276 | -0.00007 | 0.00280 | -0.00003 | 0.00288 | 0.00009 | 0.00266 |
| $s^+$ | $s^- = 0,\ s^+ = s^-$ | -0.00033 | 0.00259 | -0.00031 | 0.00259 | -0.00032 | 0.00272 | -0.00033 | 0.00264 |
| | $s^- = 0,\ s^+ > s^-$ | -0.00025 | 0.00295 | -0.00025 | 0.00286 | -0.00027 | 0.00293 | -0.00026 | 0.00274 |
| | $s^- > 0,\ s^+ < s^-$ | 0.00013 | 0.00262 | 0.00029 | 0.00276 | 0.00021 | 0.00283 | 0.00038 | 0.00255 |
| | $s^- > 0,\ s^+ = s^-$ | 0.00004 | 0.00265 | 0.00010 | 0.00282 | -0.00007 | 0.00278 | 0.00000 | 0.00258 |
| | $s^- > 0,\ s^+ > s^-$ | 0.00010 | 0.00296 | 0.00014 | 0.00296 | 0.00023 | 0.00308 | -0.00011 | 0.00278 |

(c) On average simulated datasets comprise of 7.48% genotype missing calls with an SD of 1.86% and 1.64% genotype calling errors with an SD of 0.86% ($\phi = 0.95$ and $\psi = 0.5$), corresponding to Figure S2d.

Table S2: Mean bias and RMSE in the selection coefficient estimates across different selection scenarios and data qualities, corresponding to Figure S2, continued.

| Selection coefficient | Selection scenario | Mix | | Fwd | | Bwd | | He2023a | |
|---|---|---|---|---|---|---|---|---|---|
| | | Bias | RMSE | Bias | RMSE | Bias | RMSE | Bias | RMSE |
| $s^-$ | $s^- < 0$, $s^+ < s^-$ | 0.00048 | 0.00377 | 0.00054 | 0.00382 | 0.00038 | 0.00381 | 0.00046 | 0.00364 |
| | $s^- < 0$, $s^+ = s^-$ | 0.00003 | 0.00387 | 0.00002 | 0.00328 | 0.00001 | 0.00343 | 0.00016 | 0.00303 |
| | $s^- < 0$, $s^+ > s^-$ | -0.00034 | 0.00337 | -0.00035 | 0.00346 | -0.00024 | 0.00339 | -0.00021 | 0.00314 |
| | $s^- = 0$, $s^+ < s^-$ | 0.00027 | 0.00380 | 0.00008 | 0.00351 | 0.00001 | 0.00339 | 0.00009 | 0.00331 |
| | $s^- = 0$, $s^+ = s^-$ | 0.00000 | 0.00311 | -0.00002 | 0.00314 | -0.00011 | 0.00311 | 0.00012 | 0.00302 |
| | $s^- = 0$, $s^+ > s^-$ | -0.00038 | 0.00352 | -0.00039 | 0.00370 | -0.00029 | 0.00348 | -0.00031 | 0.00338 |
| | $s^- > 0$, $s^+ < s^-$ | -0.00010 | 0.00343 | 0.00002 | 0.00382 | -0.00001 | 0.00379 | -0.00003 | 0.00341 |
| | $s^- > 0$, $s^+ = s^-$ | -0.00053 | 0.00334 | -0.00041 | 0.00339 | -0.00047 | 0.00356 | -0.00043 | 0.00319 |
| | $s^- > 0$, $s^+ > s^-$ | -0.00019 | 0.00338 | -0.00021 | 0.00337 | -0.00033 | 0.00339 | -0.00026 | 0.00315 |
| $s^+$ | $s^- < 0$, $s^+ < s^-$ | -0.00042 | 0.00297 | 0.00060 | 0.00289 | -0.00041 | 0.00308 | -0.00031 | 0.00265 |
| | $s^- < 0$, $s^+ = s^-$ | 0.00035 | 0.00267 | 0.00042 | 0.00264 | 0.00046 | 0.00282 | 0.00044 | 0.00266 |
| | $s^- < 0$, $s^+ > s^-$ | 0.00002 | 0.00342 | -0.00005 | 0.00316 | -0.00005 | 0.00338 | -0.00005 | 0.00302 |
| | $s^- = 0$, $s^+ < s^-$ | -0.00044 | 0.00317 | -0.00022 | 0.00280 | -0.00028 | 0.00318 | -0.00013 | 0.00277 |
| | $s^- = 0$, $s^+ = s^-$ | -0.00018 | 0.00278 | -0.00027 | 0.00260 | -0.00019 | 0.00280 | -0.00031 | 0.00265 |
| | $s^- = 0$, $s^+ > s^-$ | 0.00011 | 0.00298 | 0.00002 | 0.00286 | -0.00009 | 0.00302 | -0.00010 | 0.00282 |
| | $s^- > 0$, $s^+ < s^-$ | -0.00015 | 0.00291 | -0.00016 | 0.00278 | -0.00029 | 0.00296 | -0.00012 | 0.00267 |
| | $s^- > 0$, $s^+ = s^-$ | 0.00015 | 0.00286 | 0.00013 | 0.00294 | -0.00007 | 0.00287 | 0.00000 | 0.00264 |
| | $s^- > 0$, $s^+ > s^-$ | 0.00007 | 0.00303 | 0.00009 | 0.00307 | 0.00030 | 0.00318 | -0.00003 | 0.00277 |

(d) On average simulated datasets comprise of 10.96% genotype missing calls with an SD of 2.08% and 0.53% genotype calling errors with an SD of 0.50% ($\phi = 0.95$ and $\psi = 1.0$), corresponding to Figure S2e.

Table S2: Mean bias and RMSE in the selection coefficient estimates across different selection scenarios and data qualities, corresponding to Figure S2, continued.

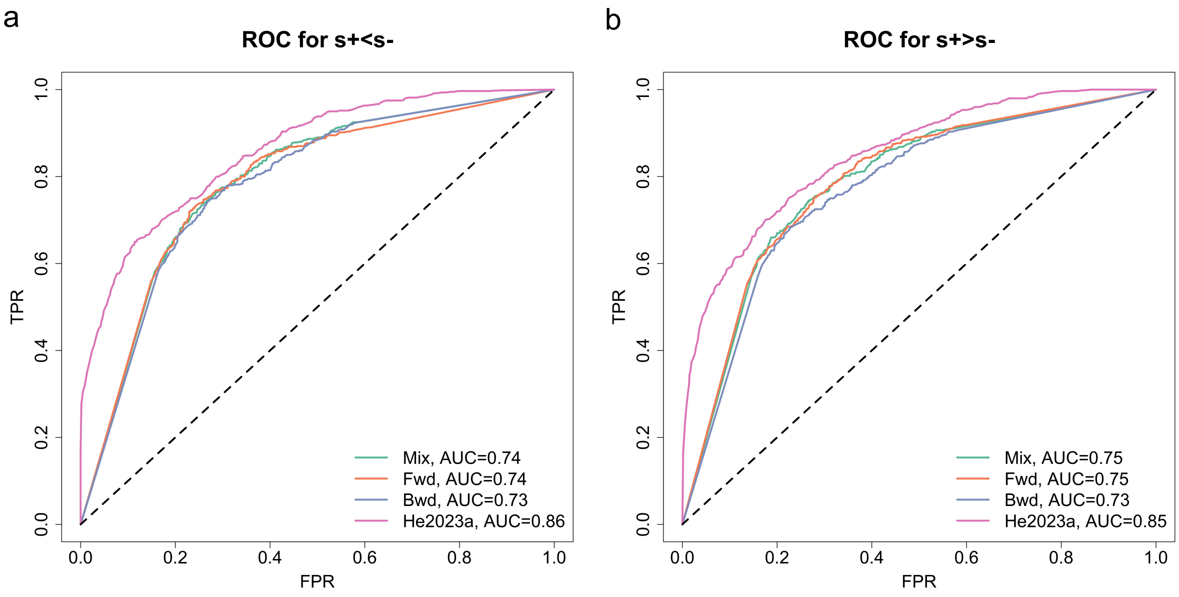


(a) On average simulated datasets comprise of 30.86% genotype missing calls with an SD of 3.27% and 9.60% genotype calling errors with an SD of 2.04% ($\phi = 0.75$ and $\psi = 0.5$).

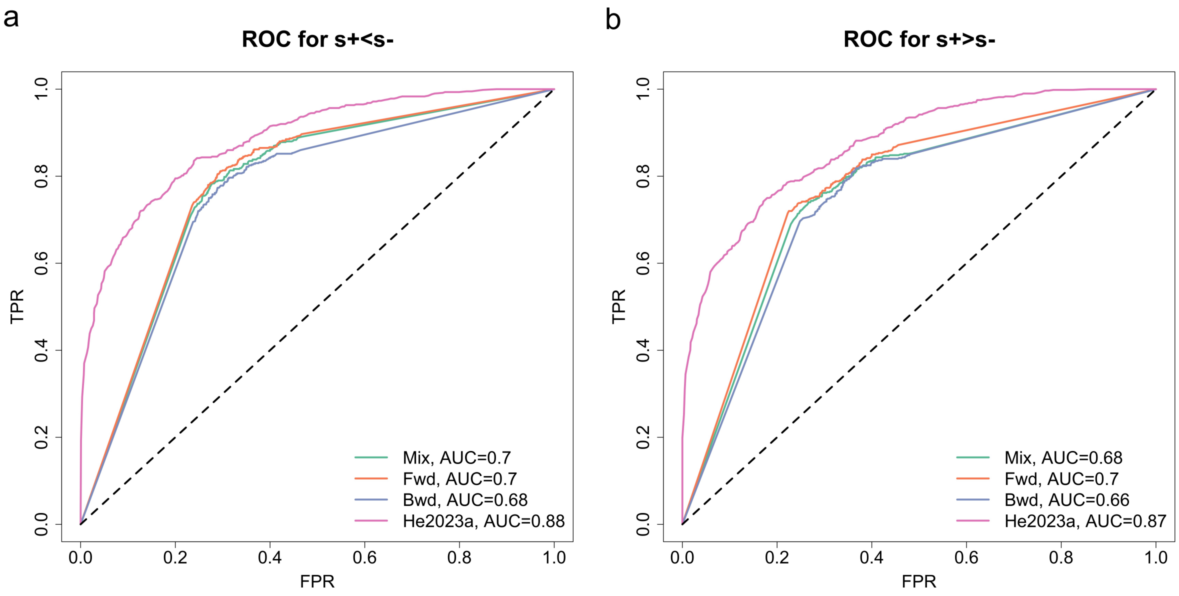


(b) On average simulated datasets comprise of 45.44% genotype missing calls with an SD of 3.35% and 3.83% genotype calling errors with an SD of 1.36% ($\phi = 0.75$ and $\psi = 1.0$).

Figure S3: ROC curves for testing selection changes across different selection scenarios and data qualities.

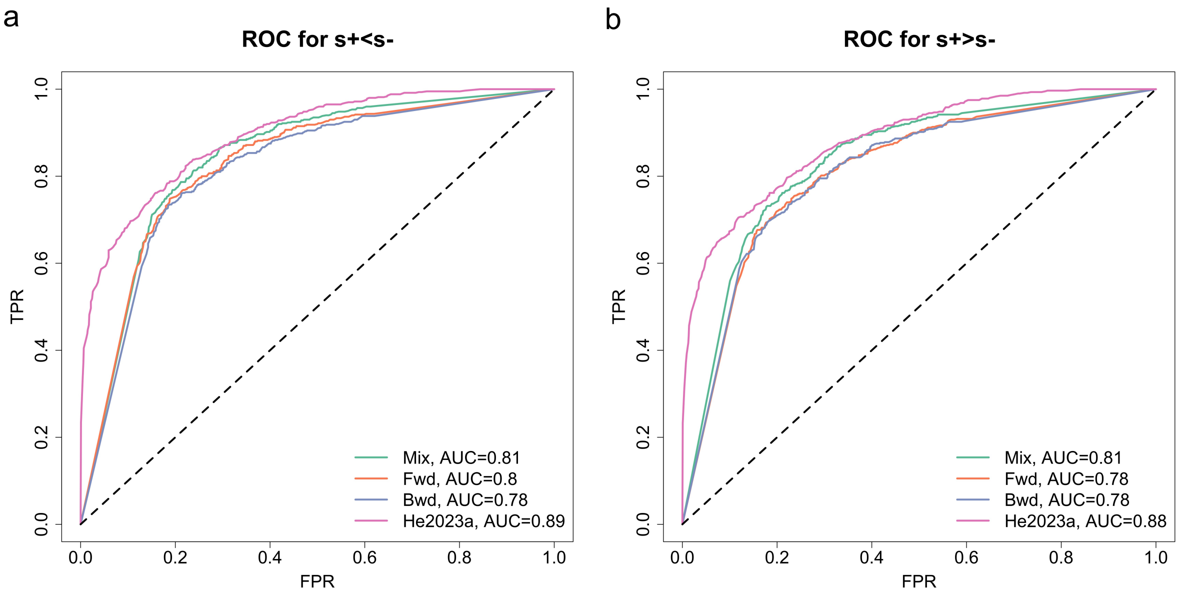


(c) On average simulated datasets comprise of 30.05% genotype missing calls with an SD of 3.18% and 1.90% genotype calling errors with an SD of 0.95% ($\phi = 0.85$ and $\psi = 1.0$).

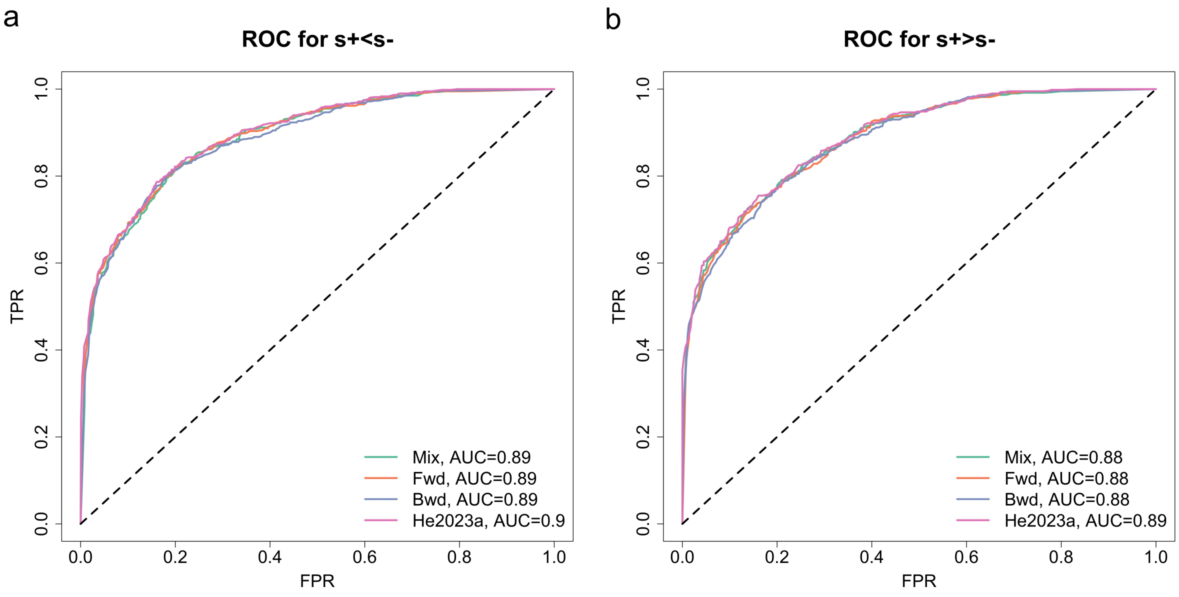


(d) On average simulated datasets comprise of 7.48% genotype missing calls with an SD of 1.86% and 1.64% genotype calling errors with an SD of 0.86% ($\phi = 0.95$ and $\psi = 0.5$).

Figure S3: ROC curves for testing selection changes across different selection scenarios and data qualities, continued.

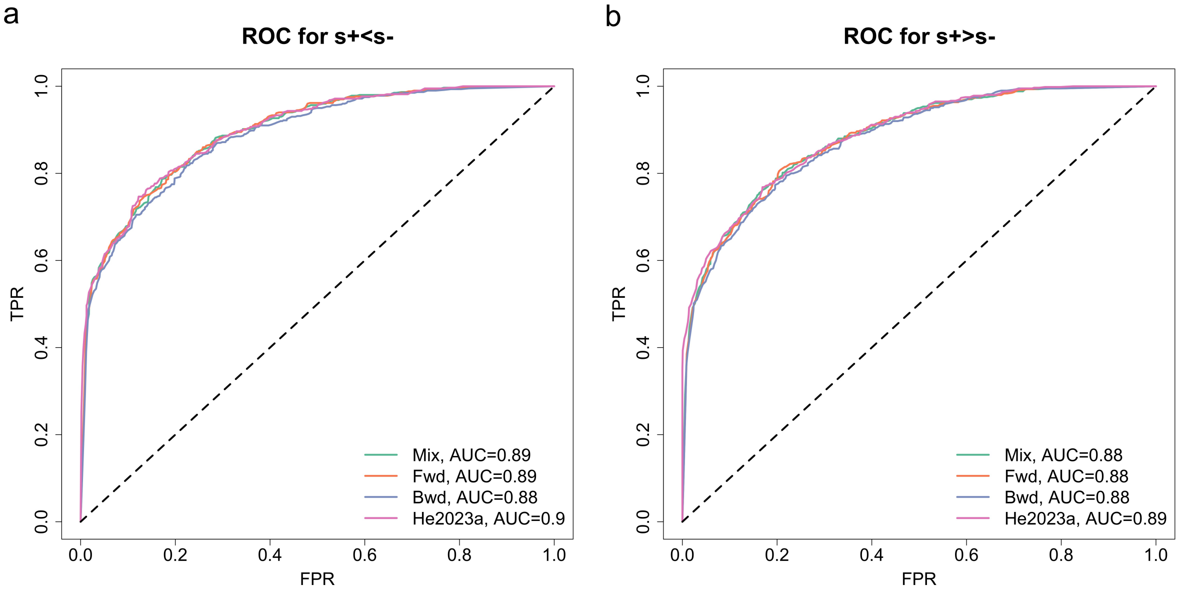


(e) On average simulated datasets comprise of 10.96% genotype missing calls with an SD of 2.08% and 0.53% genotype calling errors with an SD of 0.50% ($\phi = 0.95$ and $\psi = 1.0$).

Figure S3: ROC curves for testing selection changes across different selection scenarios and data qualities, continued.

| | | Genotype posterior | | | | | | Genotype likelihood | |
|---|---|---|---|---|---|---|---|---|---|
| | | Mix | | Fwd | | Bwd | | He2023a | |
| Date quality | Selection scenario | Mean | SD | Mean | SD | Mean | SD | Mean | SD |
| Missing rate | $s^- < 0, s^+ < s^-$ | 0.17907 | 0.03063 | 0.17788 | 0.03182 | 0.17638 | 0.03096 | 0.20493 | 0.02984 |
| | $s^- < 0, s^+ = s^-$ | 0.18048 | 0.02947 | 0.18102 | 0.02943 | 0.18138 | 0.02993 | 0.20417 | 0.02784 |
| | $s^- < 0, s^+ > s^-$ | 0.18033 | 0.03070 | 0.18005 | 0.03070 | 0.17807 | 0.03134 | 0.20674 | 0.02880 |
| | $s^- = 0, s^+ < s^-$ | 0.17810 | 0.02913 | 0.17781 | 0.02932 | 0.17805 | 0.03003 | 0.20231 | 0.02487 |
| | $s^- = 0, s^+ = s^-$ | 0.18510 | 0.03079 | 0.18452 | 0.03062 | 0.18483 | 0.03092 | 0.20245 | 0.02851 |
| | $s^- = 0, s^+ > s^-$ | 0.17750 | 0.03190 | 0.17840 | 0.03151 | 0.17760 | 0.03253 | 0.20176 | 0.02863 |
| | $s^- > 0, s^+ < s^-$ | 0.17850 | 0.03339 | 0.17888 | 0.03252 | 0.17802 | 0.03246 | 0.20505 | 0.02854 |
| | $s^- > 0, s^+ = s^-$ | 0.17836 | 0.02942 | 0.17914 | 0.02983 | 0.17955 | 0.03135 | 0.20155 | 0.02711 |
| | $s^- > 0, s^+ > s^-$ | 0.17529 | 0.02853 | 0.17519 | 0.02965 | 0.17533 | 0.02880 | 0.20576 | 0.02705 |
| Error rate | $s^- < 0, s^+ < s^-$ | 0.04450 | 0.01399 | 0.04526 | 0.01401 | 0.04576 | 0.01434 | 0.05343 | 0.01527 |
| | $s^- < 0, s^+ = s^-$ | 0.04629 | 0.01504 | 0.04607 | 0.01462 | 0.04614 | 0.01495 | 0.05340 | 0.01520 |
| | $s^- < 0, s^+ > s^-$ | 0.04524 | 0.01439 | 0.04510 | 0.01383 | 0.04581 | 0.01429 | 0.05200 | 0.01476 |
| | $s^- = 0, s^+ < s^-$ | 0.04698 | 0.01528 | 0.04664 | 0.01490 | 0.04700 | 0.01508 | 0.05221 | 0.01507 |
| | $s^- = 0, s^+ = s^-$ | 0.04724 | 0.01557 | 0.04676 | 0.01583 | 0.04700 | 0.01554 | 0.05181 | 0.01585 |
| | $s^- = 0, s^+ > s^-$ | 0.04798 | 0.01446 | 0.04810 | 0.01501 | 0.04826 | 0.01487 | 0.05386 | 0.01591 |
| | $s^- > 0, s^+ < s^-$ | 0.04645 | 0.01490 | 0.04621 | 0.01441 | 0.04669 | 0.01454 | 0.05333 | 0.01506 |
| | $s^- > 0, s^+ = s^-$ | 0.04762 | 0.01513 | 0.04743 | 0.01503 | 0.04743 | 0.01455 | 0.05317 | 0.01570 |
| | $s^- > 0, s^+ > s^-$ | 0.04550 | 0.01467 | 0.04507 | 0.01448 | 0.04483 | 0.01442 | 0.05224 | 0.01521 |

Table S3: Mean and SD in the MR and ER across different selection scenarios, corresponding to Figure 5.

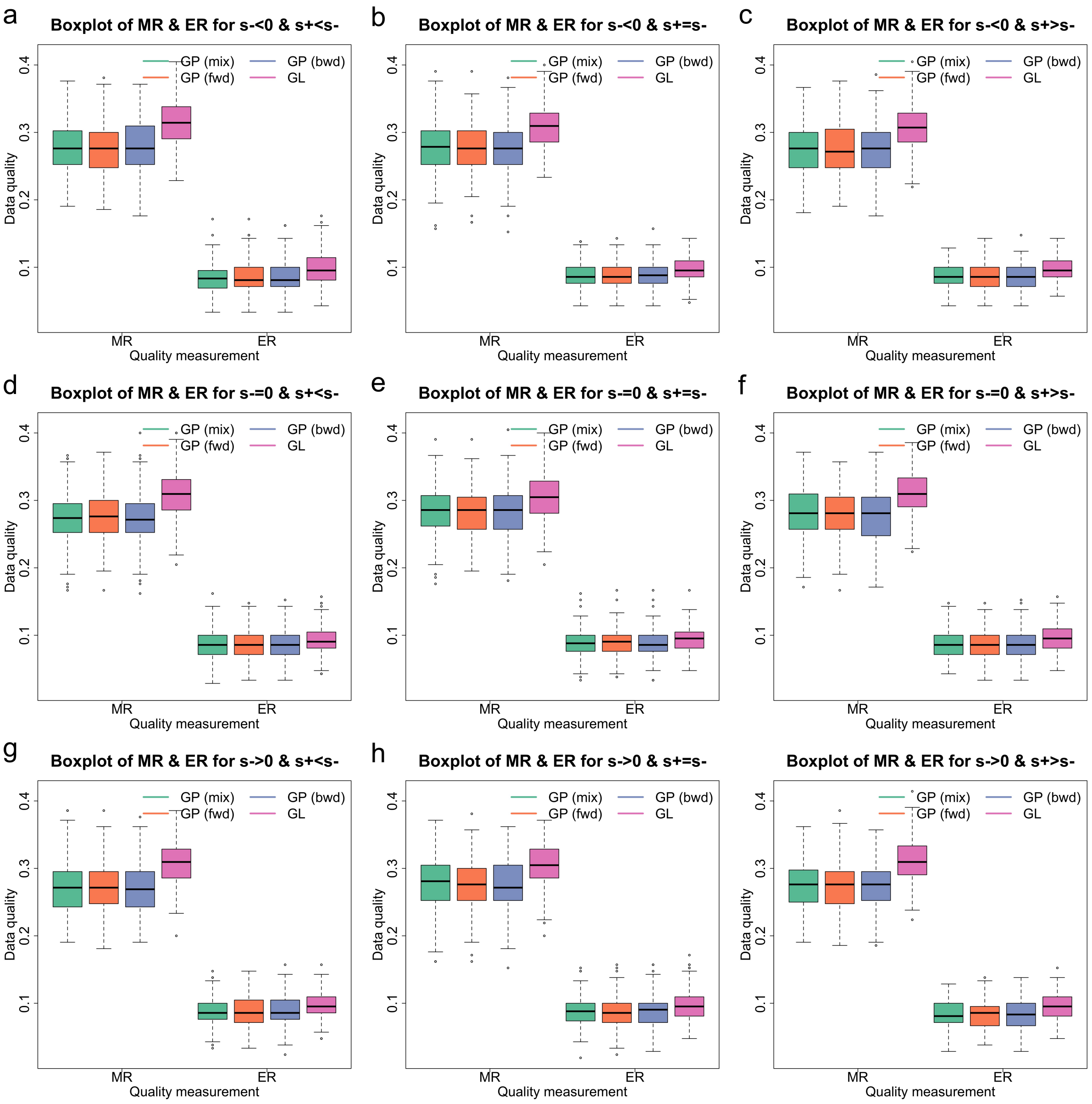


(a) On average simulated datasets comprise of 30.86% genotype missing calls with an SD of 3.27% and 9.60% genotype calling errors with an SD of 2.04% ($\phi = 0.75$ and $\psi = 0.5$).

Figure S4: Boxplots for the MR and ER across different selection scenarios and data qualities. GP and GL are shorthands for genotype posterior and genotype likelihood.

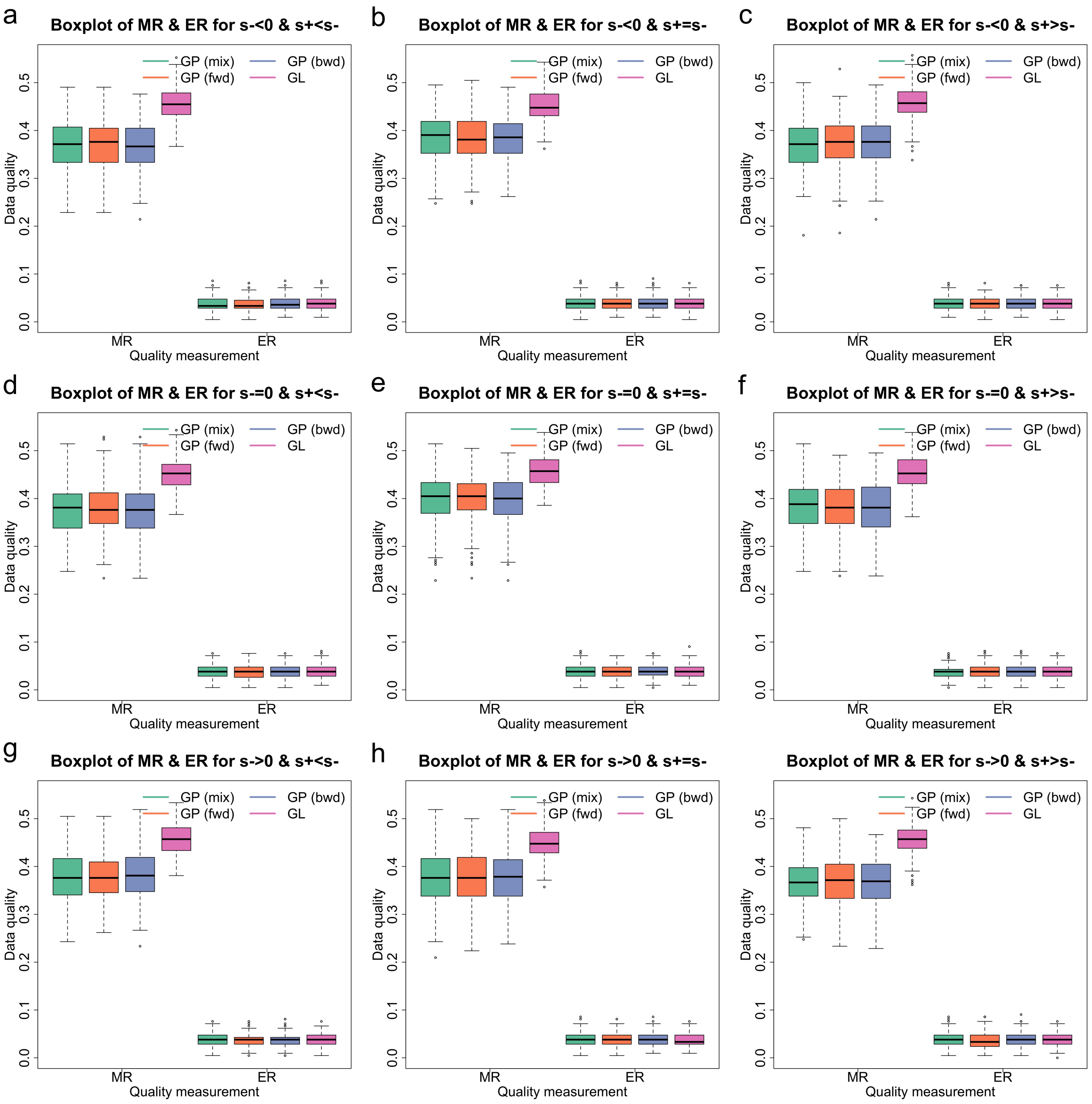


(b) On average simulated datasets comprise of 45.44% genotype missing calls with an SD of 3.35% and 3.83% genotype calling errors with an SD of 1.36% ($\phi = 0.75$ and $\psi = 1.0$).

Figure S4: Boxplots for the MR and ER across different selection scenarios and data qualities, continued. GP and GL are shorthands for genotype posterior and genotype likelihood.

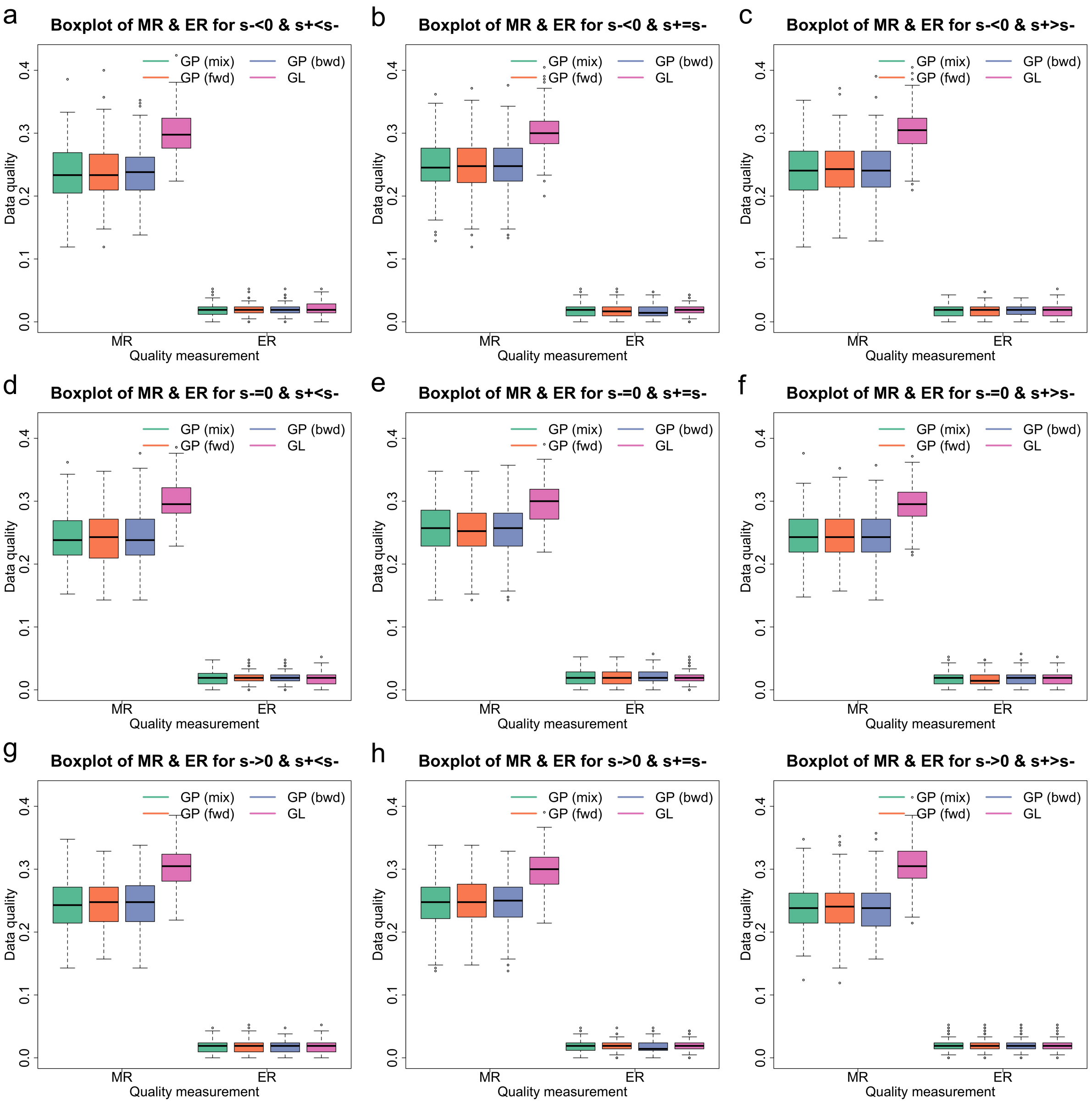


(c) On average simulated datasets comprise of 30.05% genotype missing calls with an SD of 3.18% and 1.90% genotype calling errors with an SD of 0.95% ($\phi = 0.85$ and $\psi = 1.0$).

Figure S4: Boxplots for the MR and ER across different selection scenarios and data qualities, continued. GP and GL are shorthands for genotype posterior and genotype likelihood.

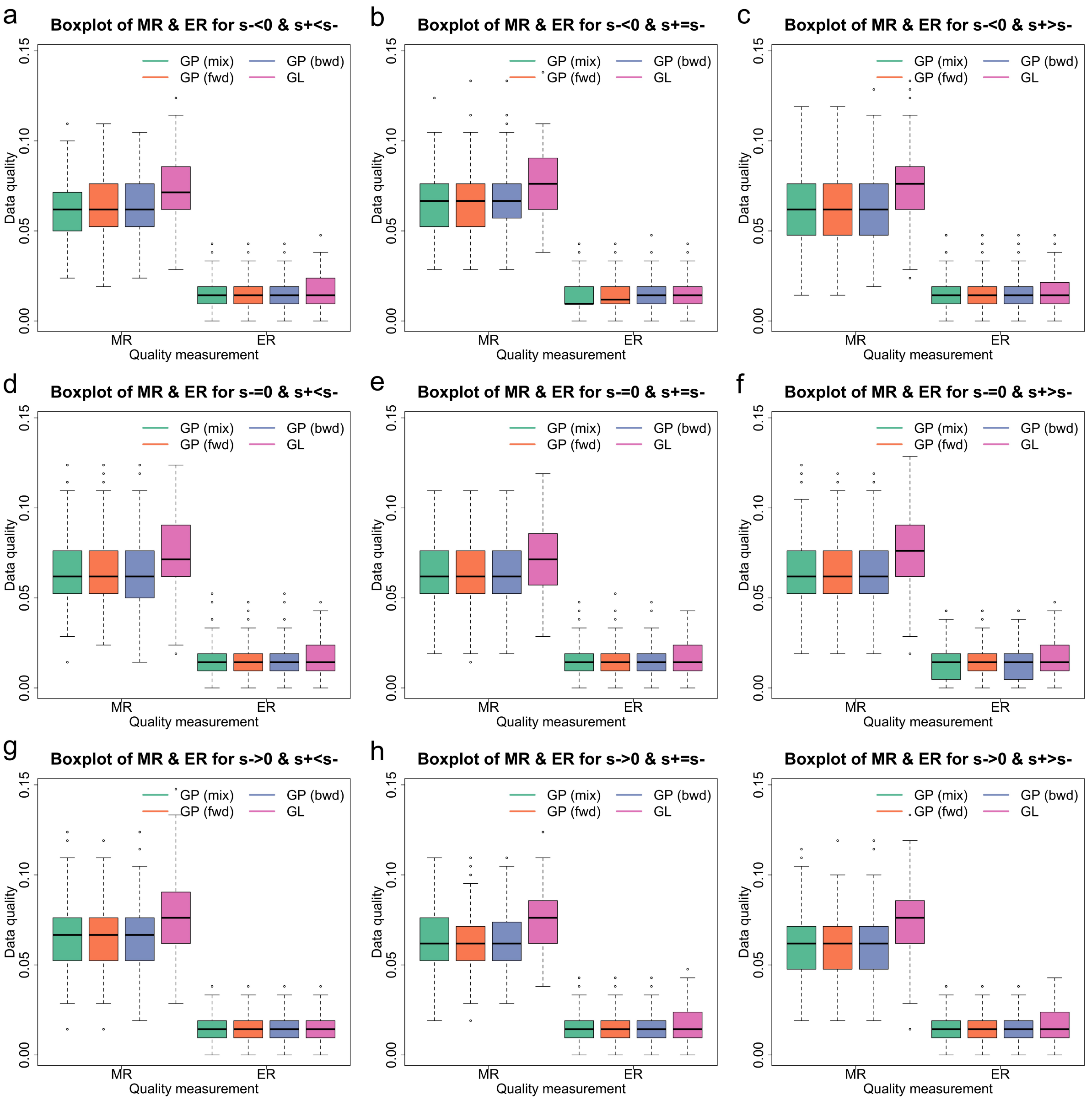


(d) On average simulated datasets comprise of 7.48% genotype missing calls with an SD of 1.86% and 1.64% genotype calling errors with an SD of 0.86% ($\phi = 0.95$ and $\psi = 0.5$).

Figure S4: Boxplots for the MR and ER across different selection scenarios and data qualities, continued. GP and GL are shorthands for genotype posterior and genotype likelihood.

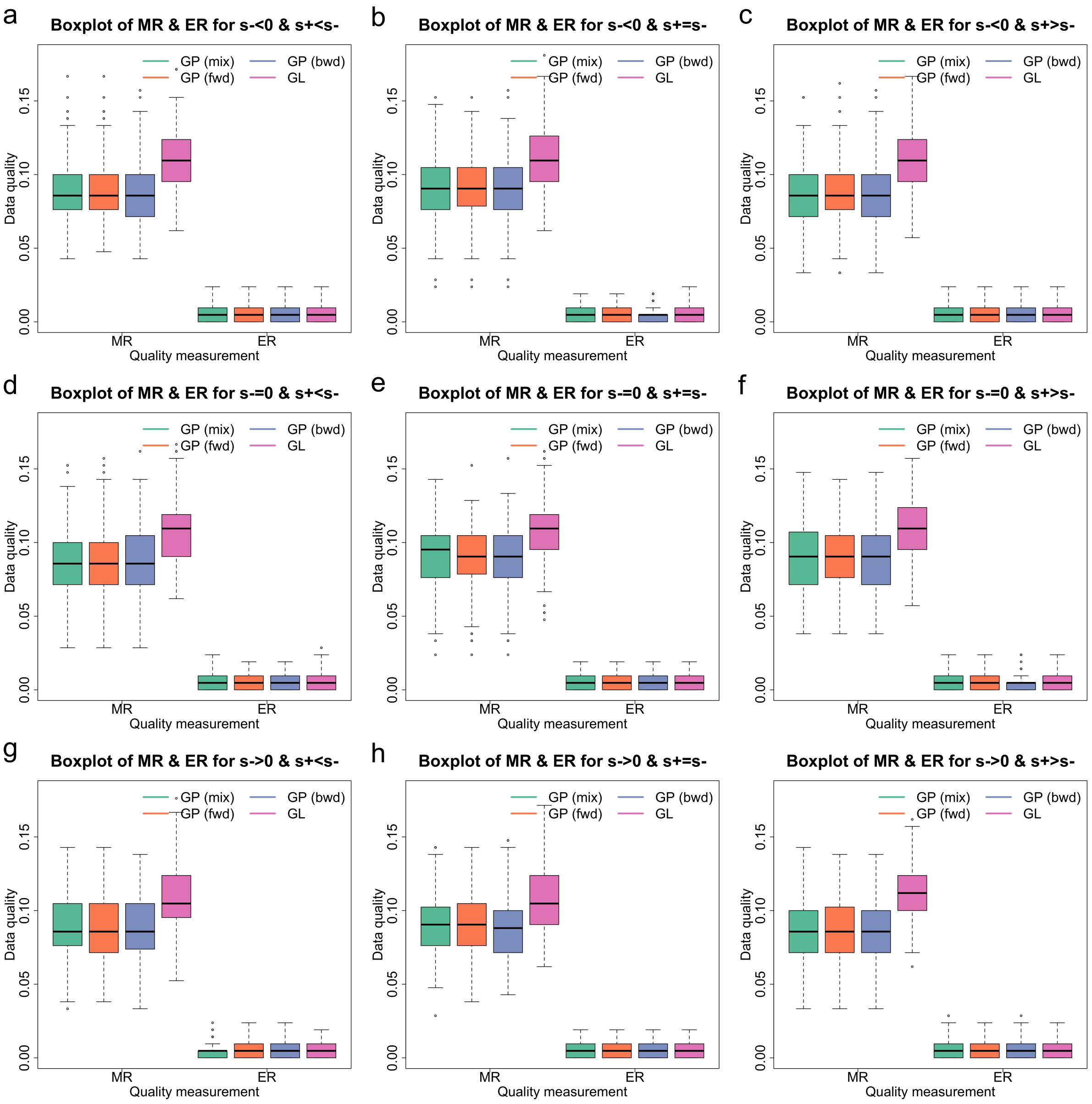


(e) On average simulated datasets comprise of 10.96% genotype missing calls with an SD of 2.08% and 0.53% genotype calling errors with an SD of 0.50% ($\phi = 0.95$ and $\psi = 1.0$).

Figure S4: Boxplots for the MR and ER across different selection scenarios and data qualities, continued. GP and GL are shorthands for genotype posterior and genotype likelihood.

| | | Genotype posterior | | | | | | Genotype likelihood | |
|---|---|---|---|---|---|---|---|---|---|
| | | Mix | | Fwd | | Bwd | | He2023a | |
| Date quality | Selection scenario | Mean | SD | Mean | SD | Mean | SD | Mean | SD |
| Missing rate | $s^- < 0,\, s^+ < s^-$ | 0.27726 | 0.03839 | 0.27612 | 0.03903 | 0.27860 | 0.03992 | 0.31452 | 0.03329 |
| | $s^- < 0,\, s^+ = s^-$ | 0.27762 | 0.03786 | 0.27764 | 0.03550 | 0.27467 | 0.03741 | 0.30769 | 0.03244 |
| | $s^- < 0,\, s^+ > s^-$ | 0.27405 | 0.03649 | 0.27486 | 0.03568 | 0.27383 | 0.03762 | 0.30683 | 0.03180 |
| | $s^- = 0,\, s^+ < s^-$ | 0.27545 | 0.03824 | 0.27650 | 0.03755 | 0.27526 | 0.03864 | 0.30802 | 0.03489 |
| | $s^- = 0,\, s^+ = s^-$ | 0.28310 | 0.03751 | 0.28267 | 0.03649 | 0.28393 | 0.03815 | 0.30712 | 0.03367 |
| | $s^- = 0,\, s^+ > s^-$ | 0.28017 | 0.03713 | 0.27945 | 0.03649 | 0.27817 | 0.03747 | 0.31040 | 0.03236 |
| | $s^- > 0,\, s^+ < s^-$ | 0.27338 | 0.03599 | 0.27174 | 0.03671 | 0.27112 | 0.03528 | 0.30667 | 0.03088 |
| | $s^- > 0,\, s^+ = s^-$ | 0.27736 | 0.03702 | 0.27588 | 0.03630 | 0.27557 | 0.03678 | 0.30576 | 0.03266 |
| | $s^- > 0,\, s^+ > s^-$ | 0.27424 | 0.03632 | 0.27329 | 0.03627 | 0.27248 | 0.03523 | 0.30995 | 0.03154 |
| Error rate | $s^- < 0,\, s^+ < s^-$ | 0.08433 | 0.02121 | 0.08469 | 0.02208 | 0.08424 | 0.02168 | 0.09590 | 0.02224 |
| | $s^- < 0,\, s^+ = s^-$ | 0.08774 | 0.01862 | 0.08786 | 0.01808 | 0.08895 | 0.01815 | 0.09679 | 0.01851 |
| | $s^- < 0,\, s^+ > s^-$ | 0.08614 | 0.01831 | 0.08588 | 0.01845 | 0.08600 | 0.01954 | 0.09740 | 0.01929 |
| | $s^- = 0,\, s^+ < s^-$ | 0.08667 | 0.02157 | 0.08600 | 0.02150 | 0.08633 | 0.02114 | 0.09424 | 0.02033 |
| | $s^- = 0,\, s^+ = s^-$ | 0.08814 | 0.02026 | 0.08833 | 0.02011 | 0.08786 | 0.02045 | 0.09524 | 0.01892 |
| | $s^- = 0,\, s^+ > s^-$ | 0.08583 | 0.02083 | 0.08612 | 0.02051 | 0.08636 | 0.02095 | 0.09557 | 0.02158 |
| | $s^- > 0,\, s^+ < s^-$ | 0.08719 | 0.02109 | 0.08776 | 0.02147 | 0.08821 | 0.02207 | 0.09767 | 0.02021 |
| | $s^- > 0,\, s^+ = s^-$ | 0.08738 | 0.02057 | 0.08724 | 0.02120 | 0.08790 | 0.02139 | 0.09719 | 0.02203 |
| | $s^- > 0,\, s^+ > s^-$ | 0.08352 | 0.01971 | 0.08374 | 0.02027 | 0.08338 | 0.01995 | 0.09393 | 0.02025 |

(a) On average simulated datasets comprise of 30.86% genotype missing calls with an SD of 3.27% and 9.60% genotype calling errors with an SD of 2.04% ($\phi = 0.75$ and $\psi = 0.5$), corresponding to Figure S4a.

Table S4: Mean and SD in the MR and ER across different selection scenarios and data qualities, corresponding to Figure S4.

| | | Genotype posterior | | | | | | Genotype likelihood | |
|---|---|---|---|---|---|---|---|---|---|
| | | Mix | | Fwd | | Bwd | | He2023a | |
| Date quality | Selection scenario | Mean | SD | Mean | SD | Mean | SD | Mean | SD |
| Missing rate | $s^- < 0$, $s^+ < s^-$ | 0.36743 | 0.05060 | 0.36786 | 0.05267 | 0.36652 | 0.05145 | 0.45455 | 0.03259 |
| | $s^- < 0$, $s^+ = s^-$ | 0.38712 | 0.04599 | 0.38471 | 0.04590 | 0.38405 | 0.04437 | 0.45374 | 0.03441 |
| | $s^- < 0$, $s^+ > s^-$ | 0.37324 | 0.05218 | 0.37602 | 0.05211 | 0.37395 | 0.05047 | 0.45652 | 0.03428 |
| | $s^- = 0$, $s^+ < s^-$ | 0.37545 | 0.05037 | 0.37667 | 0.05174 | 0.37405 | 0.05247 | 0.45269 | 0.03456 |
| | $s^- = 0$, $s^+ = s^-$ | 0.39824 | 0.05021 | 0.39914 | 0.05025 | 0.39690 | 0.04960 | 0.45626 | 0.03309 |
| | $s^- = 0$, $s^+ > s^-$ | 0.38498 | 0.05190 | 0.38195 | 0.05323 | 0.38352 | 0.05493 | 0.45486 | 0.03319 |
| | $s^- > 0$, $s^+ < s^-$ | 0.37724 | 0.05186 | 0.37719 | 0.04764 | 0.38181 | 0.05082 | 0.45583 | 0.03347 |
| | $s^- > 0$, $s^+ = s^-$ | 0.37569 | 0.05522 | 0.37545 | 0.05516 | 0.37526 | 0.05383 | 0.44893 | 0.03357 |
| | $s^- > 0$, $s^+ > s^-$ | 0.36662 | 0.04697 | 0.36824 | 0.05234 | 0.36557 | 0.05008 | 0.45640 | 0.03244 |
| Error rate | $s^- < 0$, $s^+ < s^-$ | 0.03714 | 0.01360 | 0.03726 | 0.01365 | 0.03702 | 0.01308 | 0.03876 | 0.01350 |
| | $s^- < 0$, $s^+ = s^-$ | 0.03750 | 0.01357 | 0.03790 | 0.01367 | 0.03824 | 0.01437 | 0.03831 | 0.01478 |
| | $s^- < 0$, $s^+ > s^-$ | 0.03762 | 0.01472 | 0.03674 | 0.01389 | 0.03764 | 0.01405 | 0.03771 | 0.01399 |
| | $s^- = 0$, $s^+ < s^-$ | 0.03755 | 0.01362 | 0.03719 | 0.01382 | 0.03721 | 0.01343 | 0.03781 | 0.01318 |
| | $s^- = 0$, $s^+ = s^-$ | 0.03902 | 0.01357 | 0.03886 | 0.01289 | 0.03940 | 0.01382 | 0.03986 | 0.01347 |
| | $s^- = 0$, $s^+ > s^-$ | 0.03762 | 0.01406 | 0.03843 | 0.01503 | 0.03833 | 0.01460 | 0.03867 | 0.01323 |
| | $s^- > 0$, $s^+ < s^-$ | 0.03717 | 0.01311 | 0.03664 | 0.01243 | 0.03645 | 0.01297 | 0.03733 | 0.01285 |
| | $s^- > 0$, $s^+ = s^-$ | 0.03764 | 0.01405 | 0.03819 | 0.01419 | 0.03836 | 0.01394 | 0.03793 | 0.01382 |
| | $s^- > 0$, $s^+ > s^-$ | 0.03812 | 0.01520 | 0.03724 | 0.01515 | 0.03807 | 0.01422 | 0.03805 | 0.01368 |

(b) On average simulated datasets comprise of 45.44% genotype missing calls with an SD of 3.35% and 3.83% genotype calling errors with an SD of 1.36% ($\phi = 0.75$ and $\psi = 1.0$), corresponding to Figure S4b.

Table S4: Mean and SD in the MR and ER across different selection scenarios and data qualities, corresponding to Figure S4, continued.

| | | Genotype posterior | | | | | | Genotype likelihood | |
|---|---|---|---|---|---|---|---|---|---|
| | | Mix | | Fwd | | Bwd | | He2023a | |
| Date quality | Selection scenario | Mean | SD | Mean | SD | Mean | SD | Mean | SD |
| Missing rate | $s^- < 0,\ s^+ < s^-$ | 0.23650 | 0.04253 | 0.23695 | 0.04398 | 0.23586 | 0.04182 | 0.29886 | 0.03226 |
| | $s^- < 0,\ s^+ = s^-$ | 0.24840 | 0.04100 | 0.24786 | 0.04069 | 0.24871 | 0.04076 | 0.30302 | 0.03205 |
| | $s^- < 0,\ s^+ > s^-$ | 0.23848 | 0.04143 | 0.24038 | 0.04093 | 0.24019 | 0.04085 | 0.30314 | 0.03046 |
| | $s^- = 0,\ s^+ < s^-$ | 0.24055 | 0.04009 | 0.24210 | 0.04177 | 0.24174 | 0.04246 | 0.30105 | 0.03289 |
| | $s^- = 0,\ s^+ = s^-$ | 0.25621 | 0.04018 | 0.25464 | 0.03819 | 0.25450 | 0.04053 | 0.29636 | 0.03291 |
| | $s^- = 0,\ s^+ > s^-$ | 0.24348 | 0.03947 | 0.24379 | 0.03905 | 0.24345 | 0.03893 | 0.29524 | 0.03036 |
| | $s^- > 0,\ s^+ < s^-$ | 0.24443 | 0.03964 | 0.24514 | 0.03804 | 0.24586 | 0.04006 | 0.30321 | 0.03096 |
| | $s^- > 0,\ s^+ = s^-$ | 0.24619 | 0.03876 | 0.24717 | 0.03912 | 0.24819 | 0.03919 | 0.29748 | 0.03124 |
| | $s^- > 0,\ s^+ > s^-$ | 0.23879 | 0.03852 | 0.23905 | 0.03914 | 0.23788 | 0.03981 | 0.30602 | 0.03188 |
| Error rate | $s^- < 0,\ s^+ < s^-$ | 0.01838 | 0.00943 | 0.01829 | 0.00921 | 0.01881 | 0.00930 | 0.02033 | 0.00940 |
| | $s^- < 0,\ s^+ = s^-$ | 0.01781 | 0.00949 | 0.01750 | 0.00948 | 0.01750 | 0.00929 | 0.01798 | 0.00872 |
| | $s^- < 0,\ s^+ > s^-$ | 0.01907 | 0.00951 | 0.01852 | 0.00953 | 0.01850 | 0.00930 | 0.01893 | 0.00974 |
| | $s^- = 0,\ s^+ < s^-$ | 0.01926 | 0.01014 | 0.01914 | 0.00952 | 0.01938 | 0.00955 | 0.01836 | 0.00971 |
| | $s^- = 0,\ s^+ = s^-$ | 0.01943 | 0.01113 | 0.01981 | 0.01119 | 0.01993 | 0.01095 | 0.01940 | 0.01039 |
| | $s^- = 0,\ s^+ > s^-$ | 0.01817 | 0.00938 | 0.01769 | 0.00934 | 0.01790 | 0.00943 | 0.01902 | 0.00953 |
| | $s^- > 0,\ s^+ < s^-$ | 0.01819 | 0.00957 | 0.01814 | 0.00953 | 0.01790 | 0.00891 | 0.01845 | 0.00921 |
| | $s^- > 0,\ s^+ = s^-$ | 0.01800 | 0.00884 | 0.01769 | 0.00837 | 0.01762 | 0.00891 | 0.01917 | 0.00955 |
| | $s^- > 0,\ s^+ > s^-$ | 0.01931 | 0.00951 | 0.01943 | 0.00989 | 0.01943 | 0.00978 | 0.01926 | 0.00959 |

(c) On average simulated datasets comprise of 30.05% genotype missing calls with an SD of 3.18% and 1.90% genotype calling errors with an SD of 0.95% ($\phi = 0.85$ and $\psi = 1.0$), corresponding to Figure S4c.

Table S4: Mean and SD in the MR and ER across different selection scenarios and data qualities, corresponding to Figure S4, continued.

| | | Genotype posterior | | | | | | Genotype likelihood | |
|---|---|---|---|---|---|---|---|---|---|
| | | Mix | | Fwd | | Bwd | | He2023a | |
| Date quality | Selection scenario | Mean | SD | Mean | SD | Mean | SD | Mean | SD |
| Missing rate | $s^- < 0, s^+ < s^-$ | 0.06190 | 0.01678 | 0.06229 | 0.01682 | 0.06243 | 0.01627 | 0.07379 | 0.01853 |
| | $s^- < 0, s^+ = s^-$ | 0.06574 | 0.01756 | 0.06607 | 0.01799 | 0.06650 | 0.01792 | 0.07621 | 0.01737 |
| | $s^- < 0, s^+ > s^-$ | 0.06319 | 0.01954 | 0.06310 | 0.01959 | 0.06369 | 0.01962 | 0.07571 | 0.01869 |
| | $s^- = 0, s^+ < s^-$ | 0.06379 | 0.01920 | 0.06379 | 0.01841 | 0.06381 | 0.01924 | 0.07457 | 0.01981 |
| | $s^- = 0, s^+ = s^-$ | 0.06324 | 0.01822 | 0.06350 | 0.01842 | 0.06317 | 0.01818 | 0.07102 | 0.01859 |
| | $s^- = 0, s^+ > s^-$ | 0.06514 | 0.02001 | 0.06438 | 0.01917 | 0.06450 | 0.01904 | 0.07548 | 0.01964 |
| | $s^- > 0, s^+ < s^-$ | 0.06538 | 0.01768 | 0.06519 | 0.01745 | 0.06576 | 0.01769 | 0.07721 | 0.01990 |
| | $s^- > 0, s^+ = s^-$ | 0.06381 | 0.01674 | 0.06360 | 0.01709 | 0.06369 | 0.01632 | 0.07407 | 0.01591 |
| | $s^- > 0, s^+ > s^-$ | 0.06198 | 0.01831 | 0.06150 | 0.01790 | 0.06181 | 0.01837 | 0.07529 | 0.01877 |
| Error rate | $s^- < 0, s^+ < s^-$ | 0.01405 | 0.00814 | 0.01390 | 0.00808 | 0.01376 | 0.00797 | 0.01667 | 0.00831 |
| | $s^- < 0, s^+ = s^-$ | 0.01360 | 0.00895 | 0.01357 | 0.00879 | 0.01357 | 0.00886 | 0.01529 | 0.00909 |
| | $s^- < 0, s^+ > s^-$ | 0.01510 | 0.00835 | 0.01488 | 0.00834 | 0.01490 | 0.00825 | 0.01624 | 0.00827 |
| | $s^- = 0, s^+ < s^-$ | 0.01455 | 0.00887 | 0.01440 | 0.00870 | 0.01460 | 0.00884 | 0.01683 | 0.00844 |
| | $s^- = 0, s^+ = s^-$ | 0.01521 | 0.00868 | 0.01521 | 0.00885 | 0.01526 | 0.00861 | 0.01705 | 0.00885 |
| | $s^- = 0, s^+ > s^-$ | 0.01386 | 0.00917 | 0.01405 | 0.00914 | 0.01400 | 0.00919 | 0.01619 | 0.00940 |
| | $s^- > 0, s^+ < s^-$ | 0.01376 | 0.00796 | 0.01395 | 0.00760 | 0.01379 | 0.00784 | 0.01586 | 0.00767 |
| | $s^- > 0, s^+ = s^-$ | 0.01512 | 0.00815 | 0.01505 | 0.00807 | 0.01479 | 0.00815 | 0.01705 | 0.00874 |
| | $s^- > 0, s^+ > s^-$ | 0.01374 | 0.00820 | 0.01383 | 0.00828 | 0.01402 | 0.00833 | 0.01612 | 0.00882 |

(d) On average simulated datasets comprise of 7.48% genotype missing calls with an SD of 1.86% and 1.64% genotype calling errors with an SD of 0.86% ($\phi = 0.95$ and $\psi = 0.5$), corresponding to Figure S4d.

Table S4: Mean and SD in the MR and ER across different selection scenarios and data qualities, corresponding to Figure S4, continued.

| | | Genotype posterior | | | | | | Genotype likelihood | |
|---|---|---|---|---|---|---|---|---|---|
| | | Mix | | Fwd | | Bwd | | He2023a | |
| Date quality | Selection scenario | Mean | SD | Mean | SD | Mean | SD | Mean | SD |
| Missing rate | $s^- < 0$, $s^+ < s^-$ | 0.08826 | 0.02111 | 0.08860 | 0.02153 | 0.08798 | 0.02149 | 0.11200 | 0.02085 |
| | $s^- < 0$, $s^+ = s^-$ | 0.09176 | 0.02220 | 0.09069 | 0.02141 | 0.09088 | 0.02191 | 0.11117 | 0.02092 |
| | $s^- < 0$, $s^+ > s^-$ | 0.08681 | 0.02159 | 0.08693 | 0.02161 | 0.08710 | 0.02159 | 0.10955 | 0.02163 |
| | $s^- = 0$, $s^+ < s^-$ | 0.08712 | 0.02104 | 0.08657 | 0.02099 | 0.08771 | 0.02116 | 0.10767 | 0.02101 |
| | $s^- = 0$, $s^+ = s^-$ | 0.09114 | 0.02201 | 0.09062 | 0.02226 | 0.09055 | 0.02262 | 0.10812 | 0.02135 |
| | $s^- = 0$, $s^+ > s^-$ | 0.08945 | 0.02373 | 0.08888 | 0.02287 | 0.08921 | 0.02350 | 0.11045 | 0.02172 |
| | $s^- > 0$, $s^+ < s^-$ | 0.08736 | 0.02113 | 0.08750 | 0.02170 | 0.08736 | 0.02100 | 0.10748 | 0.02132 |
| | $s^- > 0$, $s^+ = s^-$ | 0.08888 | 0.02128 | 0.08871 | 0.02120 | 0.08852 | 0.02104 | 0.10812 | 0.01953 |
| | $s^- > 0$, $s^+ > s^-$ | 0.08640 | 0.02074 | 0.08626 | 0.02055 | 0.08698 | 0.02052 | 0.11174 | 0.01844 |
| Error rate | $s^- < 0$, $s^+ < s^-$ | 0.00545 | 0.00541 | 0.00545 | 0.00539 | 0.00540 | 0.00522 | 0.00548 | 0.00535 |
| | $s^- < 0$, $s^+ = s^-$ | 0.00464 | 0.00466 | 0.00486 | 0.00487 | 0.00476 | 0.00487 | 0.00469 | 0.00499 |
| | $s^- < 0$, $s^+ > s^-$ | 0.00512 | 0.00518 | 0.00505 | 0.00504 | 0.00524 | 0.00510 | 0.00560 | 0.00535 |
| | $s^- = 0$, $s^+ < s^-$ | 0.00514 | 0.00471 | 0.00521 | 0.00476 | 0.00502 | 0.00458 | 0.00548 | 0.00522 |
| | $s^- = 0$, $s^+ = s^-$ | 0.00531 | 0.00438 | 0.00521 | 0.00447 | 0.00543 | 0.00438 | 0.00517 | 0.00437 |
| | $s^- = 0$, $s^+ > s^-$ | 0.00469 | 0.00462 | 0.00483 | 0.00454 | 0.00467 | 0.00445 | 0.00505 | 0.00479 |
| | $s^- > 0$, $s^+ < s^-$ | 0.00481 | 0.00475 | 0.00490 | 0.00472 | 0.00495 | 0.00479 | 0.00510 | 0.00464 |
| | $s^- > 0$, $s^+ = s^-$ | 0.00498 | 0.00464 | 0.00526 | 0.00471 | 0.00510 | 0.00452 | 0.00538 | 0.00497 |
| | $s^- > 0$, $s^+ > s^-$ | 0.00540 | 0.00524 | 0.00538 | 0.00508 | 0.00521 | 0.00511 | 0.00574 | 0.00549 |

(e) On average simulated datasets comprise of 10.96% genotype missing calls with an SD of 2.08% and 0.53% genotype calling errors with an SD of 0.50% ($\phi = 0.95$ and $\psi = 1.0$), corresponding to Figure S4e.

Table S4: Mean and SD in the MR and ER across different selection scenarios and data qualities, corresponding to Figure S4, continued.

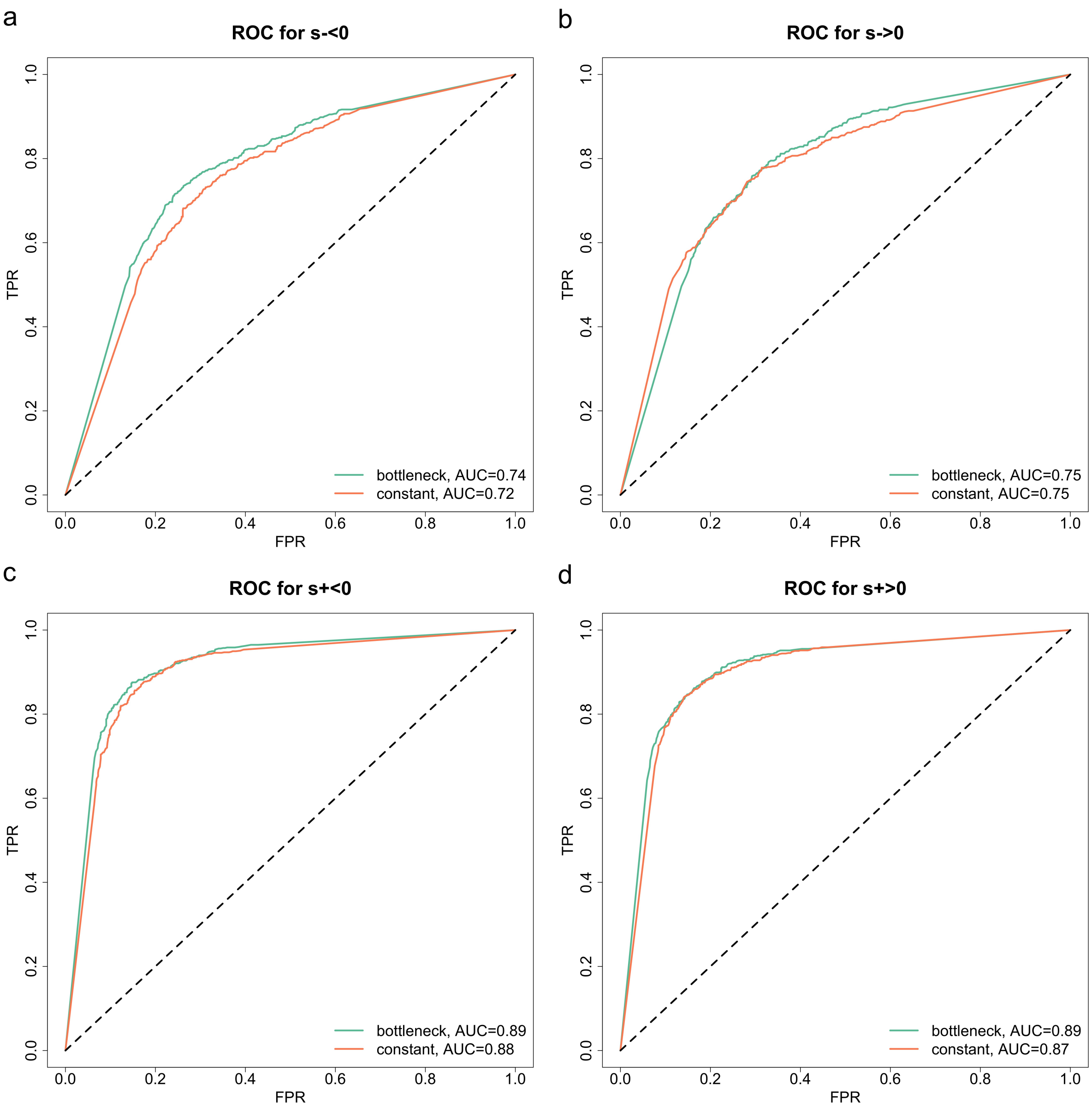


(a) ROC curves for detecting selection signatures.

Figure S5: Performance of the PMMH-within-Gibbs with the mix of forward- and backward-in-time simulations across different selection scenarios and demographic histories. On average simulated datasets comprise of 30.16% genotype missing calls with an SD of 3.22% and 1.93% genotype calling errors with an SD of 0.95% ($\phi = 0.85$ and $\psi = 0.5$).

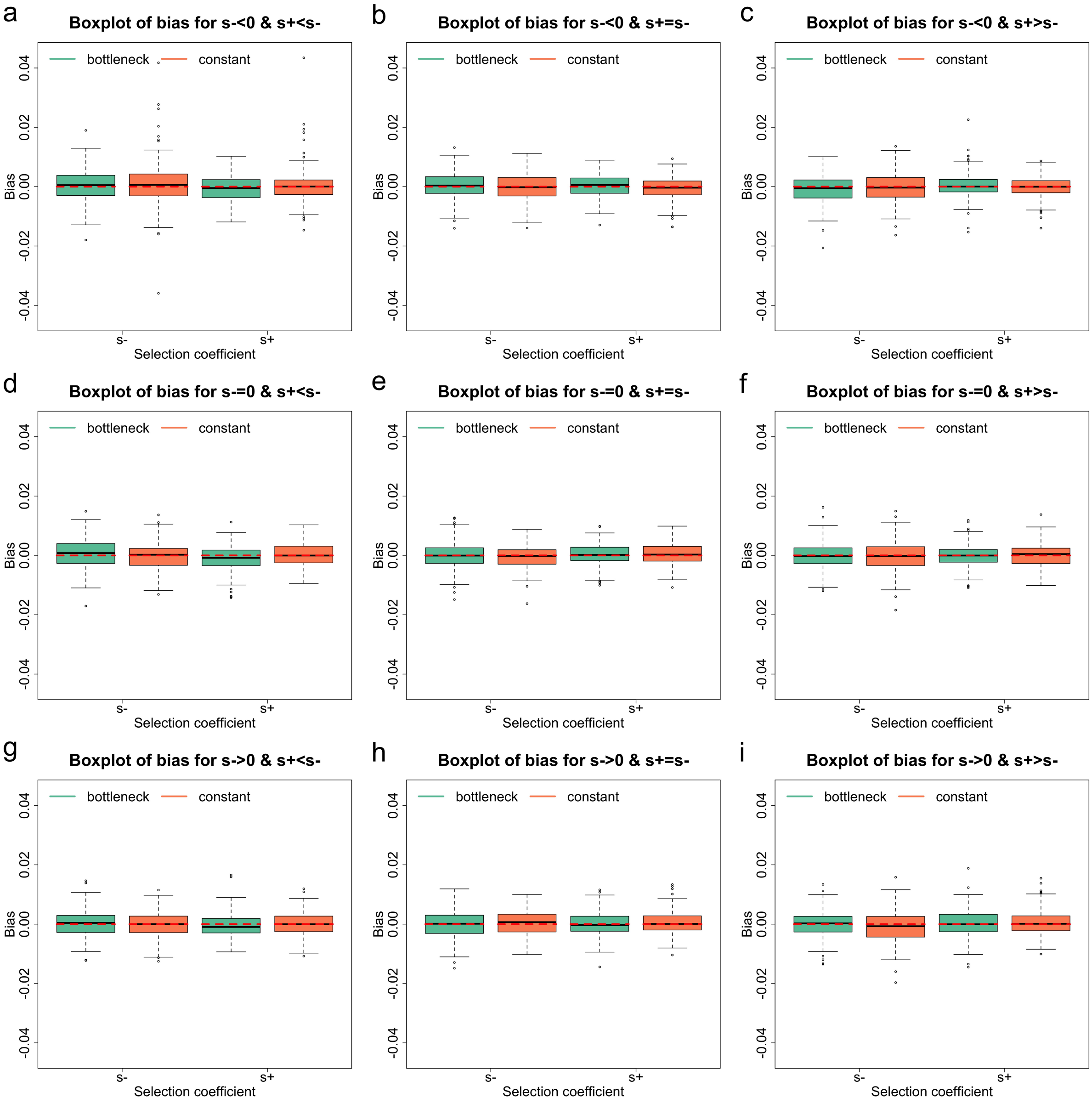


(b) Boxplots for the bias of the selection coefficient estimates.

Figure S5: Performance of the PMMH-within-Gibbs with the mix of forward- and backward-in-time simulations across different selection scenarios and demographic histories, continued. On average simulated datasets comprise of 30.16% genotype missing calls with an SD of 3.22% and 1.93% genotype calling errors with an SD of 0.95% ($\phi = 0.85$ and $\psi = 0.5$).

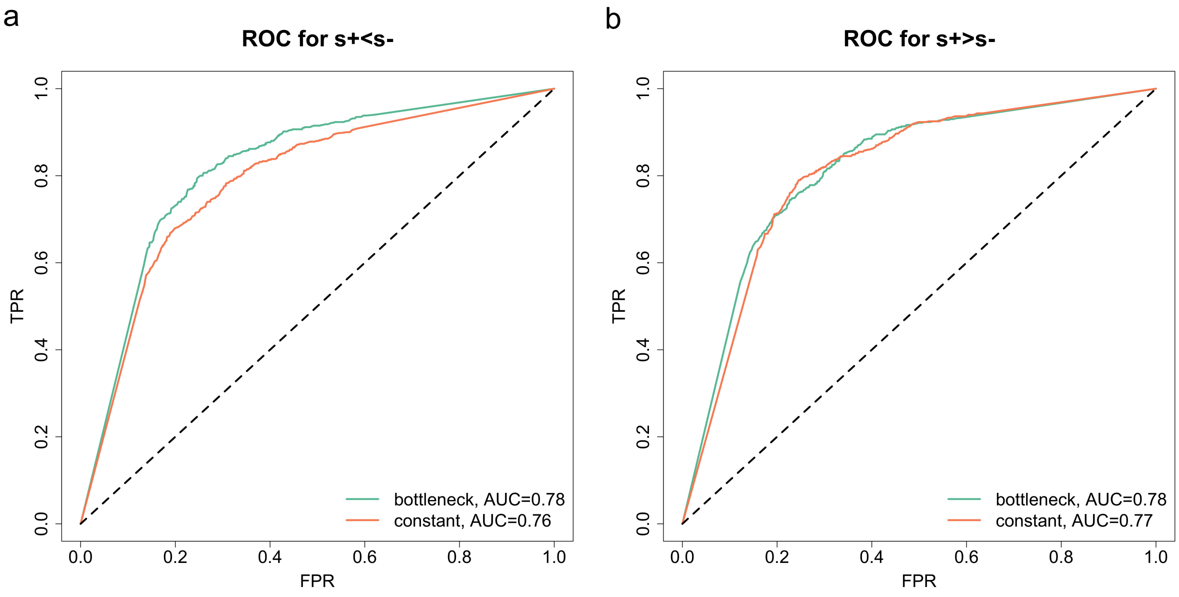


(c) ROC curves for testing selection changes.

Figure S5: Performance of the PMMH-within-Gibbs with the mix of forward- and backward-in-time simulations across different selection scenarios and demographic histories, continued. On average simulated datasets comprise of 30.16% genotype missing calls with an SD of 3.22% and 1.93% genotype calling errors with an SD of 0.95% ($\phi = 0.85$ and $\psi = 0.5$).

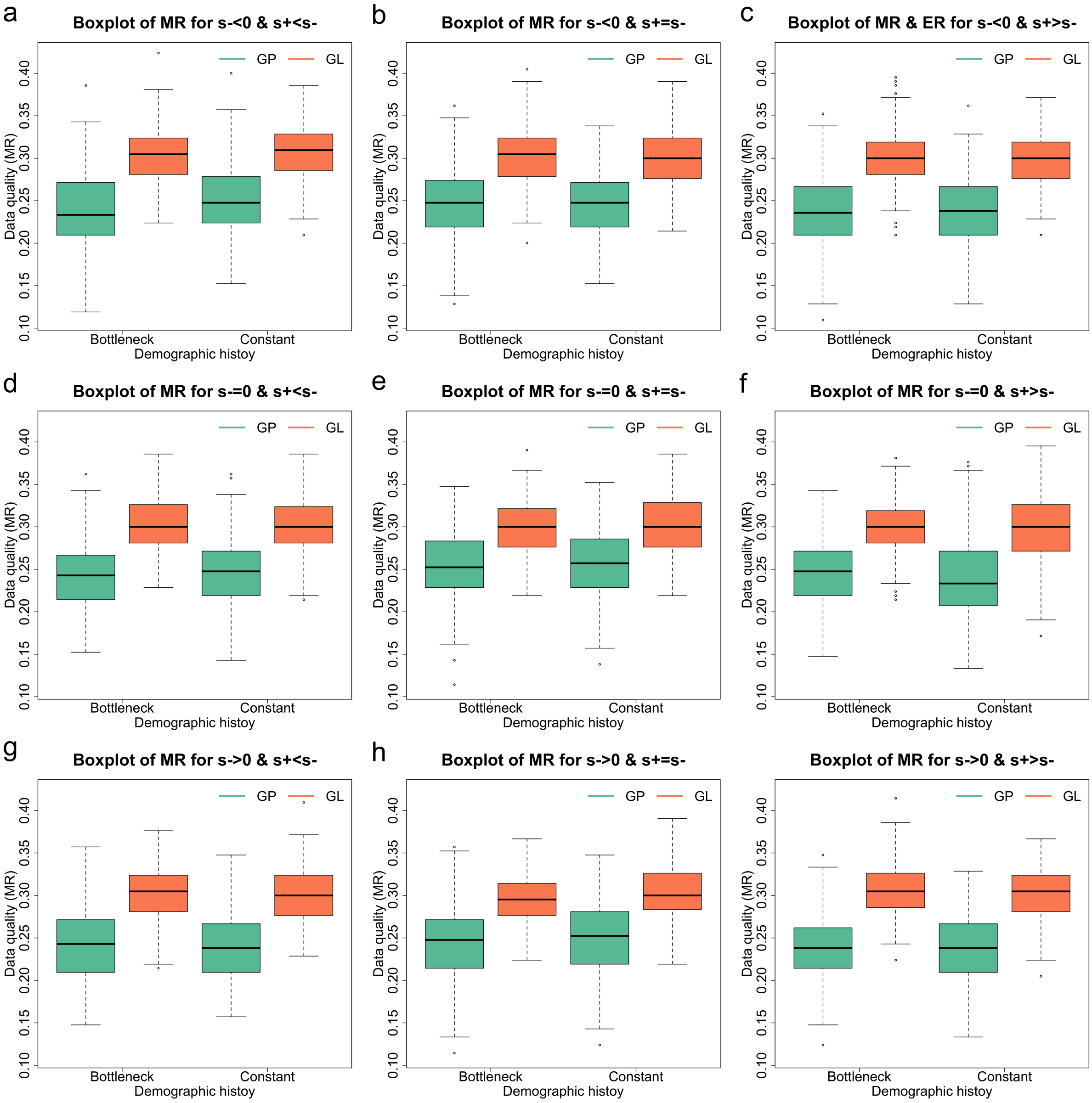


(d) Boxplots for the MR. GP and GL are shorthands for genotype posterior and genotype likelihood.

Figure S5: Performance of the PMMH-within-Gibbs with the mix of forward- and backward-in-time simulations across different selection scenarios and demographic histories, continued. On average simulated datasets comprise of 30.16% genotype missing calls with an SD of 3.22% and 1.93% genotype calling errors with an SD of 0.95% ($\phi = 0.85$ and $\psi = 0.5$).

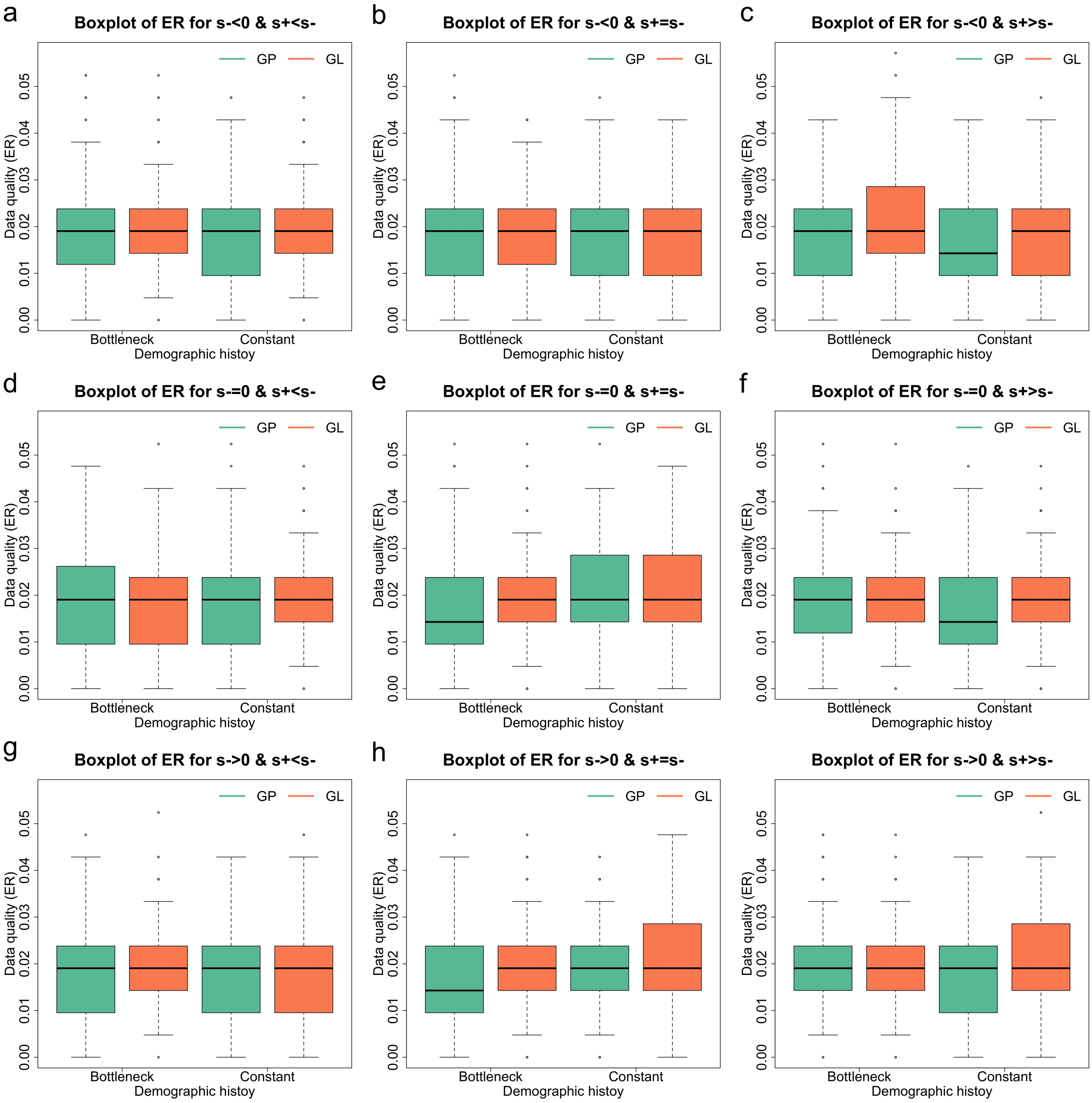


(e) Boxplots for the ER. GP and GL are shorthands for genotype posterior and genotype likelihood.

Figure S5: Performance of the PMMH-within-Gibbs with the mix of forward- and backward-in-time simulations across different selection scenarios and demographic histories, continued. On average simulated datasets comprise of 30.16% genotype missing calls with an SD of 3.22% and 1.93% genotype calling errors with an SD of 0.95% ($\phi = 0.85$ and $\psi = 0.5$).

| | | Bottleneck | | Constant | |
|---|---|---|---|---|---|
| Selection coefficient | Selection scenario | Bias | RMSE | Bias | RMSE |
| | $s^- < 0$, $s^+ < s^-$ | 0.00038 | 0.00525 | 0.00077 | 0.00761 |
| | $s^- < 0$, $s^+ = s^-$ | 0.00040 | 0.00482 | -0.00003 | 0.00474 |
| | $s^- < 0$, $s^+ > s^-$ | -0.00081 | 0.00482 | -0.00002 | 0.00506 |
| | $s^- = 0$, $s^+ < s^-$ | 0.00057 | 0.00486 | -0.00033 | 0.00469 |
| $s^-$ | $s^- = 0$, $s^+ = s^-$ | 0.00014 | 0.00457 | -0.00039 | 0.00396 |
| | $s^- = 0$, $s^+ > s^-$ | -0.00023 | 0.00441 | -0.00018 | 0.00483 |
| | $s^- > 0$, $s^+ < s^-$ | 0.00024 | 0.00451 | -0.00018 | 0.00445 |
| | $s^- > 0$, $s^+ = s^-$ | -0.00003 | 0.00460 | 0.00027 | 0.00434 |
| | $s^- > 0$, $s^+ > s^-$ | 0.00000 | 0.00456 | -0.00081 | 0.00517 |
| | $s^- < 0$, $s^+ < s^-$ | -0.00078 | 0.00412 | 0.00022 | 0.00572 |
| | $s^- < 0$, $s^+ = s^-$ | 0.00029 | 0.00359 | -0.00047 | 0.00369 |
| | $s^- < 0$, $s^+ > s^-$ | 0.00046 | 0.00425 | -0.00017 | 0.00362 |
| | $s^- = 0$, $s^+ < s^-$ | -0.00090 | 0.00429 | 0.00007 | 0.00393 |
| $s^+$ | $s^- = 0$, $s^+ = s^-$ | 0.00010 | 0.00368 | 0.00037 | 0.00358 |
| | $s^- = 0$, $s^+ > s^-$ | -0.00005 | 0.00392 | 0.00013 | 0.00400 |
| | $s^- > 0$, $s^+ < s^-$ | -0.00052 | 0.00402 | 0.00001 | 0.00382 |
| | $s^- > 0$, $s^+ = s^-$ | 0.00000 | 0.00394 | 0.00037 | 0.00401 |
| | $s^- > 0$, $s^+ > s^-$ | 0.00031 | 0.00434 | 0.00049 | 0.00416 |

(a) Mean bias and RMSE in the selection coefficient estimates, corresponding to Figure S5b.

Table S8: Performance of the PMMH-within-Gibbs with the mix of forward- and backward-in-time simulations across different selection scenarios and demographic histories, corresponding to Figure S5. On average simulated datasets comprise of 30.16% genotype missing calls with an SD of 3.22% and 1.93% genotype calling errors with an SD of 0.95% ($\phi = 0.85$ and $\psi = 0.5$).

| | | Genotype posterior | | | | Genotype likelihood | | | |
|---|---|---|---|---|---|---|---|---|---|
| | | Bottleneck | | Constant | | Bottleneck | | Constant | |
| Date quality | Selection scenario | Mean | SD | Mean | SD | Mean | SD | Mean | SD |
| Missing rate | $s^- < 0, s^+ < s^-$ | 0.23869 | 0.04314 | 0.25007 | 0.02149 | 0.30295 | 0.03372 | 0.30600 | 0.03165 |
| | $s^- < 0, s^+ = s^-$ | 0.24776 | 0.04104 | 0.24576 | 0.03913 | 0.30379 | 0.03462 | 0.30155 | 0.03258 |
| | $s^- < 0, s^+ > s^-$ | 0.23610 | 0.03949 | 0.23679 | 0.03928 | 0.30055 | 0.03159 | 0.29710 | 0.03030 |
| | $s^- = 0, s^+ < s^-$ | 0.24002 | 0.03996 | 0.24526 | 0.04285 | 0.30464 | 0.03273 | 0.30329 | 0.03205 |
| | $s^- = 0, s^+ = s^-$ | 0.25317 | 0.04115 | 0.25676 | 0.04336 | 0.29812 | 0.03139 | 0.30183 | 0.03509 |
| | $s^- = 0, s^+ > s^-$ | 0.24619 | 0.04075 | 0.24000 | 0.04832 | 0.29945 | 0.03249 | 0.29824 | 0.03654 |
| | $s^- > 0, s^+ < s^-$ | 0.24131 | 0.04095 | 0.23952 | 0.03974 | 0.30269 | 0.03082 | 0.29950 | 0.03157 |
| | $s^- > 0, s^+ = s^-$ | 0.24290 | 0.04106 | 0.24971 | 0.04240 | 0.29721 | 0.02929 | 0.30321 | 0.02967 |
| | $s^- > 0, s^+ > s^-$ | 0.23795 | 0.03819 | 0.23912 | 0.03740 | 0.30590 | 0.03112 | 0.30229 | 0.03016 |
| Error rate | $s^- < 0, s^+ < s^-$ | 0.01869 | 0.01039 | 0.01867 | 0.00959 | 0.01993 | 0.00943 | 0.01926 | 0.00935 |
| | $s^- < 0, s^+ = s^-$ | 0.01831 | 0.00968 | 0.01805 | 0.00961 | 0.01852 | 0.00934 | 0.01857 | 0.00965 |
| | $s^- < 0, s^+ > s^-$ | 0.01893 | 0.00947 | 0.01648 | 0.00976 | 0.01986 | 0.01031 | 0.01867 | 0.00964 |
| | $s^- = 0, s^+ < s^-$ | 0.01893 | 0.01014 | 0.01926 | 0.00993 | 0.01821 | 0.00946 | 0.01967 | 0.00936 |
| | $s^- = 0, s^+ = s^-$ | 0.01824 | 0.01014 | 0.02069 | 0.01019 | 0.01914 | 0.00929 | 0.02012 | 0.01035 |
| | $s^- = 0, s^+ > s^-$ | 0.01831 | 0.00921 | 0.01731 | 0.00956 | 0.01902 | 0.00946 | 0.01912 | 0.00906 |
| | $s^- > 0, s^+ < s^-$ | 0.01831 | 0.00956 | 0.01840 | 0.00900 | 0.01852 | 0.00867 | 0.01940 | 0.01002 |
| | $s^- > 0, s^+ = s^-$ | 0.01812 | 0.00936 | 0.01912 | 0.00864 | 0.01886 | 0.00940 | 0.01990 | 0.00929 |
| | $s^- > 0, s^+ > s^-$ | 0.01881 | 0.00879 | 0.01867 | 0.00914 | 0.01952 | 0.00897 | 0.02029 | 0.01004 |

(b) Mean and SD in the MR and ER, corresponding to Figures S5d and S5e.

Table S8: Performance of the PMMH-within-Gibbs with the mix of forward- and backward-in-time simulations across different selection scenarios and demographic histories, corresponding to Figure S5, continued. On average simulated datasets comprise of 30.16% genotype missing calls with an SD of 3.22% and 1.93% genotype calling errors with an SD of 0.95% ($\phi = 0.85$ and $\psi = 0.5$).

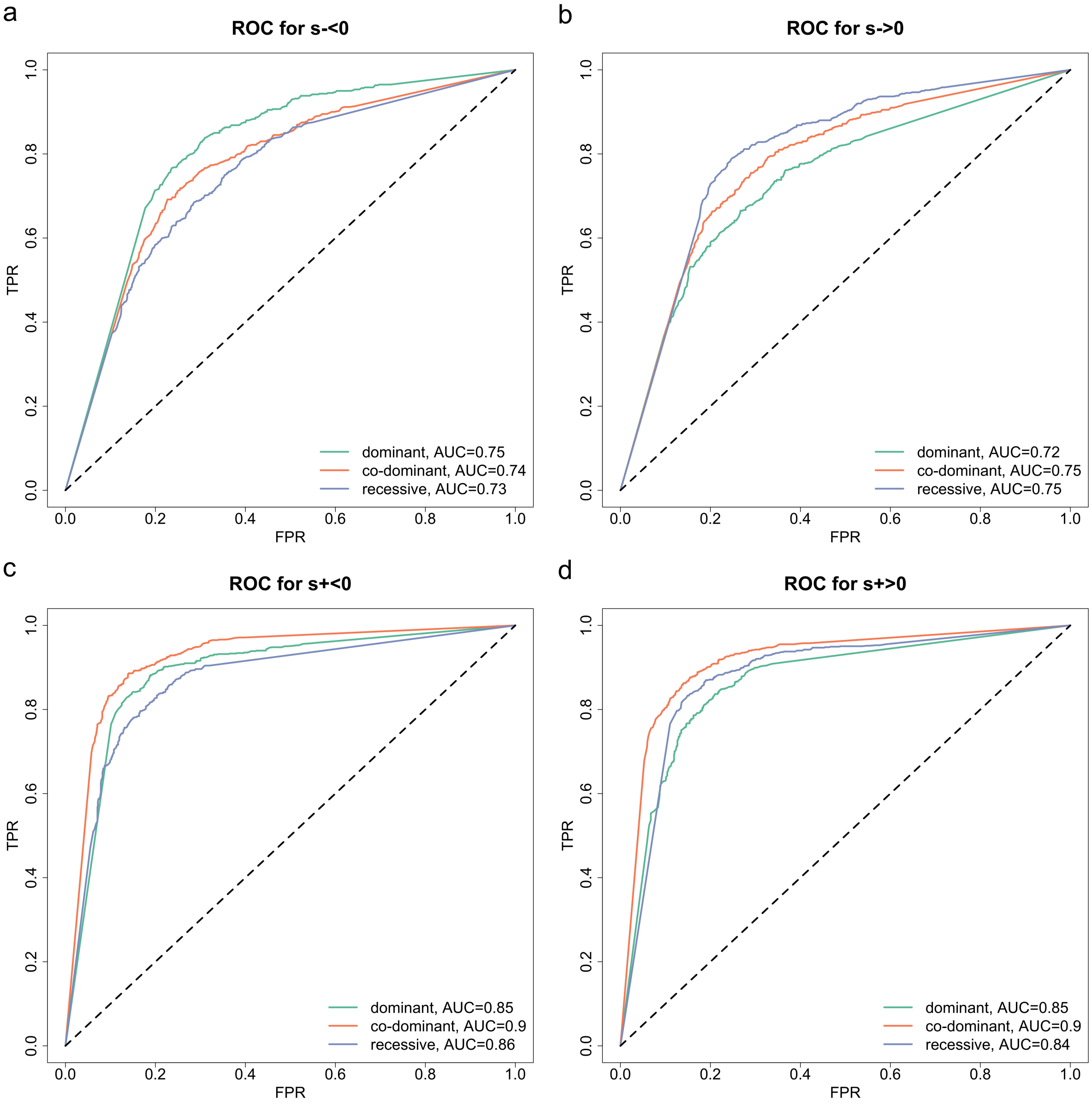


(a) ROC curves for detecting selection signatures.

Figure S6: Performance of the PMMH-within-Gibbs with the mix of forward- and backward-in-time simulations across different selection scenarios and dominance levels. On average simulated datasets comprise of 30.14% genotype missing calls with an SD of 3.18% and 1.91% genotype calling errors with an SD of 0.93% ($\phi = 0.85$ and $\psi = 0.5$).

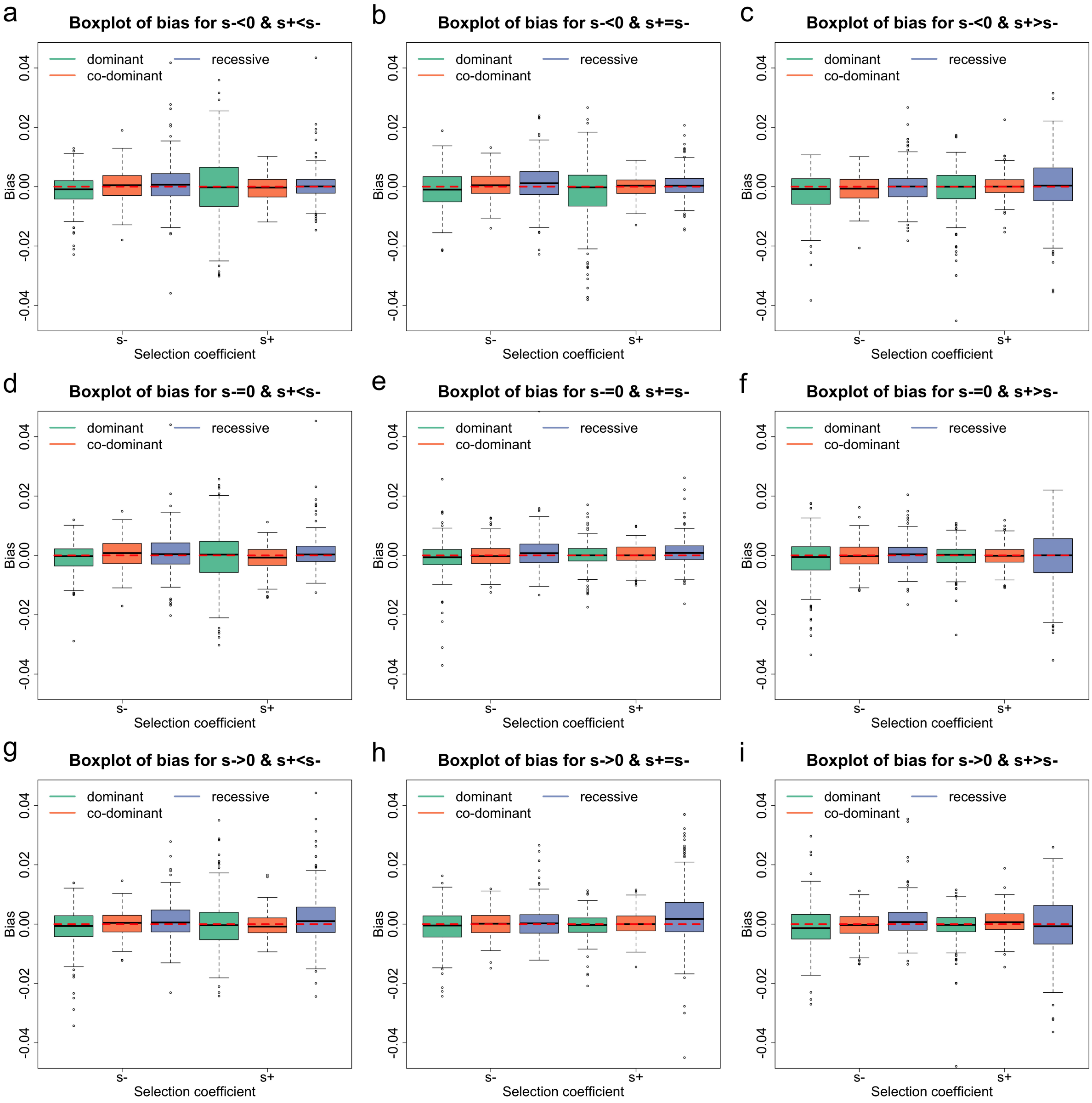


(b) Boxplots for the bias of the selection coefficient estimates.

Figure S6: Performance of the PMMH-within-Gibbs with the mix of forward- and backward-in-time simulations across different selection scenarios and dominance levels, continued. On average simulated datasets comprise of 30.14% genotype missing calls with an SD of 3.18% and 1.91% genotype calling errors with an SD of 0.93% ($\phi = 0.85$ and $\psi = 0.5$).

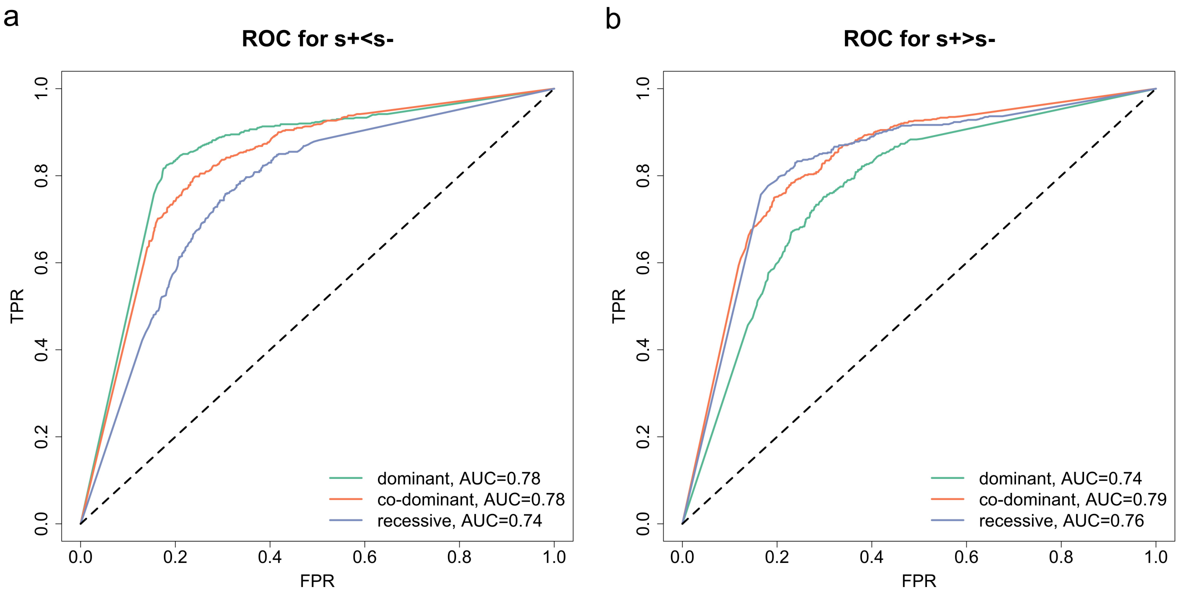


(c) ROC curves for testing selection changes.

Figure S6: Performance of the PMMH-within-Gibbs with the mix of forward- and backward-in-time simulations across different selection scenarios and dominance levels, continued. On average simulated datasets comprise of 30.14% genotype missing calls with an SD of 3.18% and 1.91% genotype calling errors with an SD of 0.93% ($\phi = 0.85$ and $\psi = 0.5$).

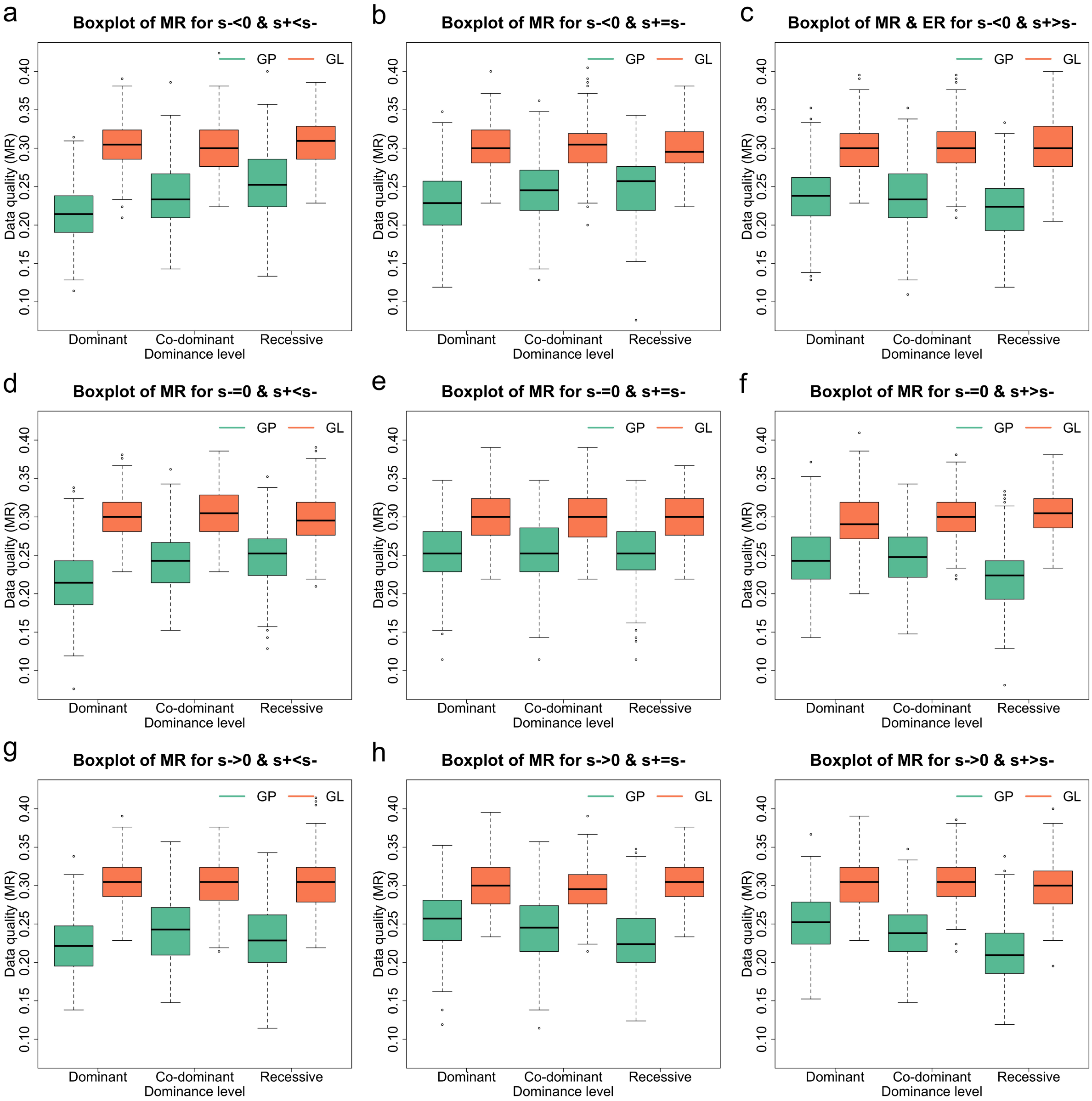


(d) Boxplots for the MR. GP and GL are shorthands for genotype posterior and genotype likelihood.

Figure S6: Performance of the PMMH-within-Gibbs with the mix of forward- and backward-in-time simulations across different selection scenarios and dominance levels, continued. On average simulated datasets comprise of 30.14% genotype missing calls with an SD of 3.18% and 1.91% genotype calling errors with an SD of 0.93% ($\phi = 0.85$ and $\psi = 0.5$).

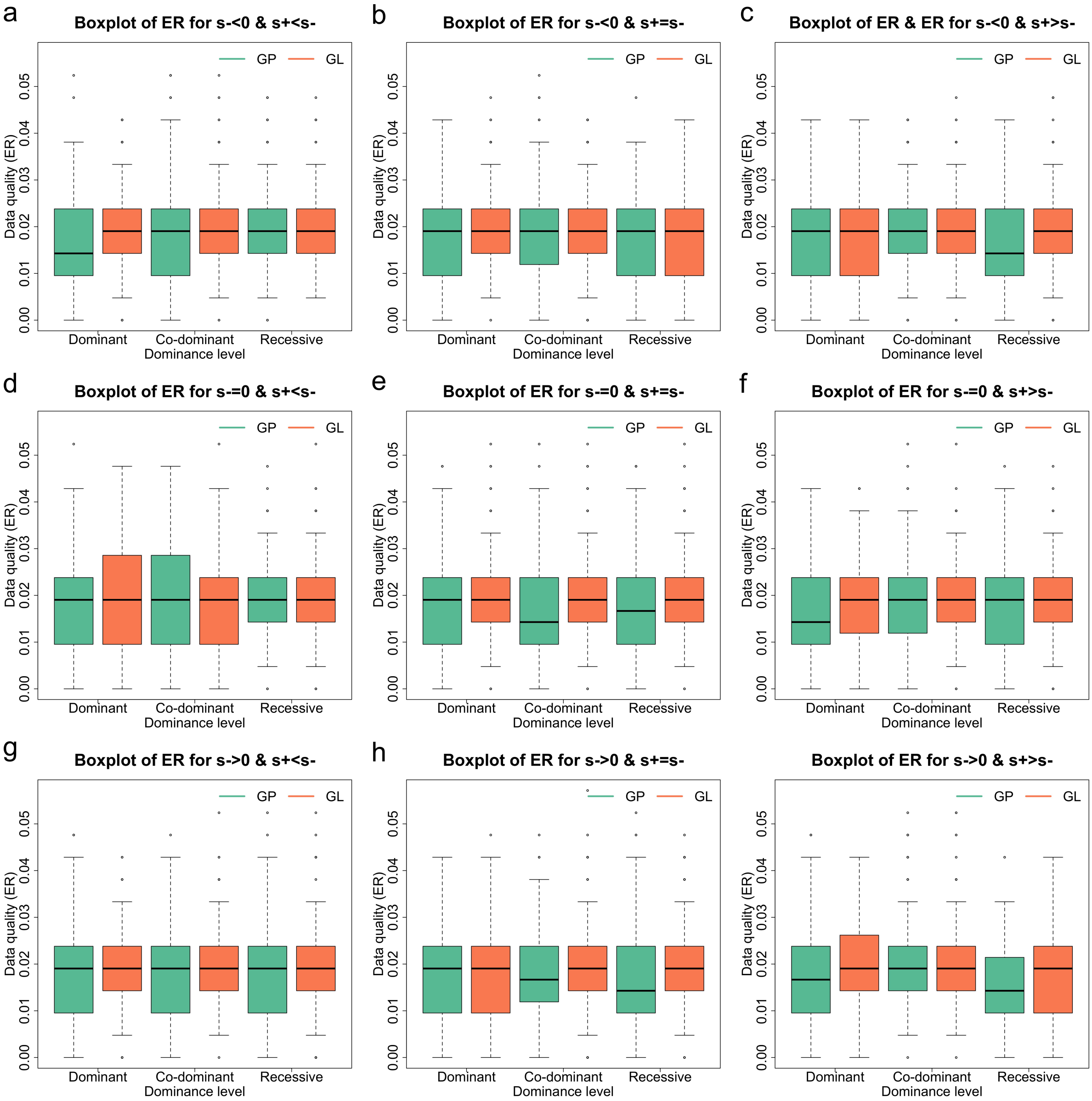


(e) Boxplots for the ER. GP and GL are shorthands for genotype posterior and genotype likelihood.

Figure S6: Performance of the PMMH-within-Gibbs with the mix of forward- and backward-in-time simulations across different selection scenarios and dominance levels, continued. On average simulated datasets comprise of 30.14% genotype missing calls with an SD of 3.18% and 1.91% genotype calling errors with an SD of 0.93% ($\phi = 0.85$ and $\psi = 0.5$).

| Selection coefficient | Selection scenario | Dominant | | Co-dominant | | Recessive | |
|---|---|---|---|---|---|---|---|
| | | Bias | RMSE | Bias | RMSE | Bias | RMSE |
| $s^-$ | $s^- < 0$, $s^+ < s^-$ | -0.00108 | 0.00552 | 0.00035 | 0.00511 | 0.00093 | 0.00774 |
| | $s^- < 0$, $s^+ = s^-$ | -0.00115 | 0.00648 | 0.00054 | 0.00489 | 0.00132 | 0.00693 |
| | $s^- < 0$, $s^+ > s^-$ | -0.00203 | 0.00712 | -0.00083 | 0.00468 | 0.00028 | 0.00620 |
| | $s^- = 0$, $s^+ < s^-$ | -0.00090 | 0.00514 | 0.00048 | 0.00492 | 0.00080 | 0.00692 |
| | $s^- = 0$, $s^+ = s^-$ | -0.00078 | 0.00625 | -0.00002 | 0.00440 | 0.00115 | 0.00627 |
| | $s^- = 0$, $s^+ > s^-$ | -0.00157 | 0.00799 | -0.00023 | 0.00443 | 0.00046 | 0.00497 |
| | $s^- > 0$, $s^+ < s^-$ | -0.00151 | 0.00669 | 0.00036 | 0.00445 | 0.00103 | 0.00629 |
| | $s^- > 0$, $s^+ = s^-$ | -0.00114 | 0.00610 | 0.00004 | 0.00443 | 0.00089 | 0.00606 |
| | $s^- > 0$, $s^+ > s^-$ | -0.00104 | 0.00780 | -0.00042 | 0.00471 | 0.00132 | 0.00636 |
| $s^+$ | $s^- < 0$, $s^+ < s^-$ | -0.00027 | 0.01184 | -0.00060 | 0.00404 | 0.00032 | 0.00572 |
| | $s^- < 0$, $s^+ = s^-$ | -0.00211 | 0.01121 | 0.00004 | 0.00344 | 0.00044 | 0.00478 |
| | $s^- < 0$, $s^+ > s^-$ | -0.00099 | 0.00828 | 0.00033 | 0.00426 | 0.00028 | 0.00990 |
| | $s^- = 0$, $s^+ < s^-$ | 0.00002 | 0.00922 | -0.00079 | 0.00428 | 0.00103 | 0.00592 |
| | $s^- = 0$, $s^+ = s^-$ | 0.00000 | 0.00443 | 0.00020 | 0.00362 | 0.00124 | 0.00504 |
| | $s^- = 0$, $s^+ > s^-$ | -0.00018 | 0.00462 | 0.00004 | 0.00388 | -0.00042 | 0.00973 |
| | $s^- > 0$, $s^+ < s^-$ | 0.00035 | 0.00903 | -0.00045 | 0.00407 | 0.00221 | 0.00943 |
| | $s^- > 0$, $s^+ = s^-$ | -0.00052 | 0.00443 | 0.00015 | 0.00396 | 0.00294 | 0.01150 |
| | $s^- > 0$, $s^+ > s^-$ | -0.00069 | 0.00586 | 0.00067 | 0.00430 | -0.00084 | 0.01075 |

(a) Mean bias and RMSE in the selection coefficient estimates across different selection scenarios, corresponding to Figure S6b.

Table S9: Performance of the PMMH-within-Gibbs with the mix of forward- and backward-in-time simulations across different demographic histories, corresponding to Figure S6. On average simulated datasets comprise of 30.14% genotype missing calls with an SD of 3.18% and 1.91% genotype calling errors with an SD of 0.93% ($\phi = 0.85$ and $\psi = 0.5$).

| | | Genotype posterior | | | | | | Genotype likelihood | | | | | |
|---|---|---|---|---|---|---|---|---|---|---|---|---|---|
| | | Dominant | | Co-dominant | | Recessive | | Dominant | | Co-dominant | | Recessive | |
| Date quality | Selection scenario | Mean | SD | Mean | SD | Mean | SD | Mean | SD | Mean | SD | Mean | SD |
| Missing rate | $s^- < 0, s^+ < s^-$ | 0.21619 | 0.03934 | 0.23845 | 0.04178 | 0.25295 | 0.04523 | 0.30445 | 0.03261 | 0.30076 | 0.03375 | 0.30702 | 0.03114 |
| | $s^- < 0, s^+ = s^-$ | 0.22886 | 0.04264 | 0.24693 | 0.04003 | 0.24995 | 0.04210 | 0.30167 | 0.02959 | 0.30307 | 0.03262 | 0.29886 | 0.03104 |
| | $s^- < 0, s^+ > s^-$ | 0.23564 | 0.04309 | 0.23567 | 0.04031 | 0.22236 | 0.04165 | 0.30119 | 0.03204 | 0.30026 | 0.03185 | 0.30114 | 0.03426 |
| | $s^- = 0, s^+ < s^-$ | 0.21598 | 0.04204 | 0.24017 | 0.03953 | 0.24721 | 0.04010 | 0.29929 | 0.03034 | 0.30543 | 0.03257 | 0.29783 | 0.03118 |
| | $s^- = 0, s^+ = s^-$ | 0.25290 | 0.04149 | 0.25402 | 0.04255 | 0.25321 | 0.04129 | 0.29943 | 0.03227 | 0.29833 | 0.03209 | 0.29864 | 0.03024 |
| | $s^- = 0, s^+ > s^-$ | 0.24607 | 0.04246 | 0.24729 | 0.03999 | 0.22095 | 0.04189 | 0.29590 | 0.03395 | 0.30024 | 0.03246 | 0.30512 | 0.02908 |
| | $s^- > 0, s^+ < s^-$ | 0.22271 | 0.03909 | 0.24293 | 0.04093 | 0.22990 | 0.04501 | 0.30462 | 0.03060 | 0.30312 | 0.03026 | 0.30255 | 0.03468 |
| | $s^- > 0, s^+ = s^-$ | 0.25236 | 0.04001 | 0.24345 | 0.04161 | 0.22971 | 0.04523 | 0.30093 | 0.03191 | 0.29776 | 0.03059 | 0.30400 | 0.02909 |
| | $s^- > 0, s^+ > s^-$ | 0.25098 | 0.03831 | 0.23693 | 0.03834 | 0.21317 | 0.03908 | 0.30188 | 0.03131 | 0.30514 | 0.03039 | 0.29945 | 0.03340 |
| Error rate | $s^- < 0, s^+ < s^-$ | 0.01693 | 0.00894 | 0.01867 | 0.00998 | 0.01888 | 0.00922 | 0.01979 | 0.00971 | 0.02021 | 0.00943 | 0.01921 | 0.00882 |
| | $s^- < 0, s^+ = s^-$ | 0.01736 | 0.00883 | 0.01867 | 0.00975 | 0.01755 | 0.00841 | 0.01936 | 0.00920 | 0.01890 | 0.00943 | 0.01867 | 0.00949 |
| | $s^- < 0, s^+ > s^-$ | 0.01776 | 0.00879 | 0.01893 | 0.00931 | 0.01698 | 0.00920 | 0.01824 | 0.00853 | 0.01952 | 0.00968 | 0.01938 | 0.00930 |
| | $s^- = 0, s^+ < s^-$ | 0.01717 | 0.00982 | 0.01895 | 0.01017 | 0.01848 | 0.00851 | 0.01945 | 0.00976 | 0.01817 | 0.00982 | 0.01888 | 0.00909 |
| | $s^- = 0, s^+ = s^-$ | 0.01855 | 0.01020 | 0.01800 | 0.00991 | 0.01824 | 0.00981 | 0.01936 | 0.00971 | 0.01940 | 0.00945 | 0.01929 | 0.00938 |
| | $s^- = 0, s^+ > s^-$ | 0.01633 | 0.00871 | 0.01829 | 0.00889 | 0.01795 | 0.00902 | 0.01810 | 0.00869 | 0.01881 | 0.00912 | 0.01862 | 0.00875 |
| | $s^- > 0, s^+ < s^-$ | 0.01800 | 0.00948 | 0.01812 | 0.00954 | 0.01795 | 0.01015 | 0.01921 | 0.00879 | 0.01867 | 0.00909 | 0.01921 | 0.00952 |
| | $s^- > 0, s^+ = s^-$ | 0.01833 | 0.00930 | 0.01819 | 0.00958 | 0.01767 | 0.01070 | 0.01902 | 0.00979 | 0.01936 | 0.00990 | 0.01852 | 0.00928 |
| | $s^- > 0, s^+ > s^-$ | 0.01824 | 0.00982 | 0.01940 | 0.00954 | 0.01660 | 0.00810 | 0.01964 | 0.00909 | 0.02012 | 0.00952 | 0.01871 | 0.00871 |

(b) Mean and SD in the MR and ER across different selection scenarios, corresponding to Figures S6d and S6e.

Table S9: Performance of the PMMH-within-Gibbs with the mix of forward- and backward-in-time simulations across different demographic histories, corresponding to Figure S6, continued. On average simulated datasets comprise of 30.14% genotype missing calls with an SD of 3.18% and 1.91% genotype calling errors with an SD of 0.93% ($\phi = 0.85$ and $\psi = 0.5$).

#### File S2. Additional results for analysis of real data

| Gene | Time period | Selection coefficient | MAP estimate | 95% HPD | Pr($s<0$) | Pr($s>0$) |
|---|---|---|---|---|---|---|
| *MC1R* | [−514, −106] | $s^-$ | 0.01157 | [ 0.00386, 0.02265] | 0.0050 | 0.9950 |
| | | $s^+$ | 0.00919 | [−0.02376, 0.03106] | 0.3065 | 0.6935 |
| | | $\Delta s$ | −0.00148 | [−0.04302, 0.02613] | 0.6405 | 0.3595 |
| *KIT13* | [−514, −106] | $s^-$ | 0.01002 | [ 0.00105, 0.01905] | 0.0160 | 0.9840 |
| | | $s^+$ | −0.05437 | [−0.09222, −0.01335] | 0.9995 | 0.0005 |
| | | $\Delta s$ | −0.06311 | [−0.11077, −0.01746] | 0.9995 | 0.0005 |
| *TRPM1* | [−514, −106] | $s^-$ | 0.00903 | [−0.01994, 0.02682] | 0.3335 | 0.6665 |
| | | $s^+$ | −0.05113 | [−0.12987, 0.04373] | 0.8025 | 0.1975 |
| | | $\Delta s$ | −0.05597 | [−0.14028, 0.06648] | 0.7960 | 0.2040 |

Table S6: Estimates of the selection coefficients and their changes with their 95% HPD intervals, as well as posterior probabilities for negative selection/change and positive selection/change, for *MC1R*, *KIT13* and *TRPM1*, corresponding to Figures 6–8, produced through the PMMH-within-Gibbs procedure with the mix of the forward- and backward-in-time simulation. The time period is measured in generations (8 years per generation) and offset so that 2000 AD is 0.

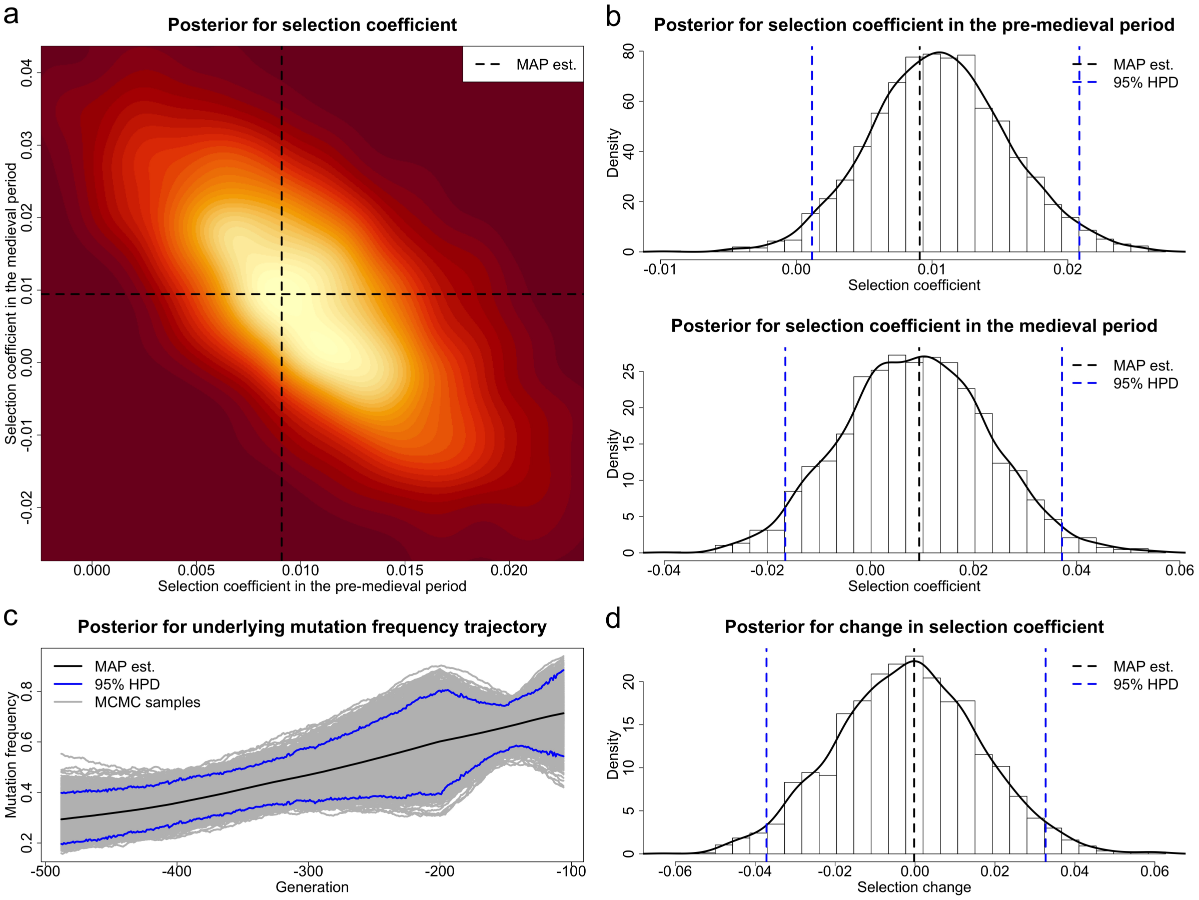


Figure S7: Posteriors for the selection coefficients of the *MC1R* mutation in the pre-medieval and medieval period and the underlying frequency trajectory of the *MC1R* mutation in the population produced through the PMMH-within-Gibbs procedure with the full forward-in-time simulation. The samples drawn in the period $[-514, -488)$ are excluded.

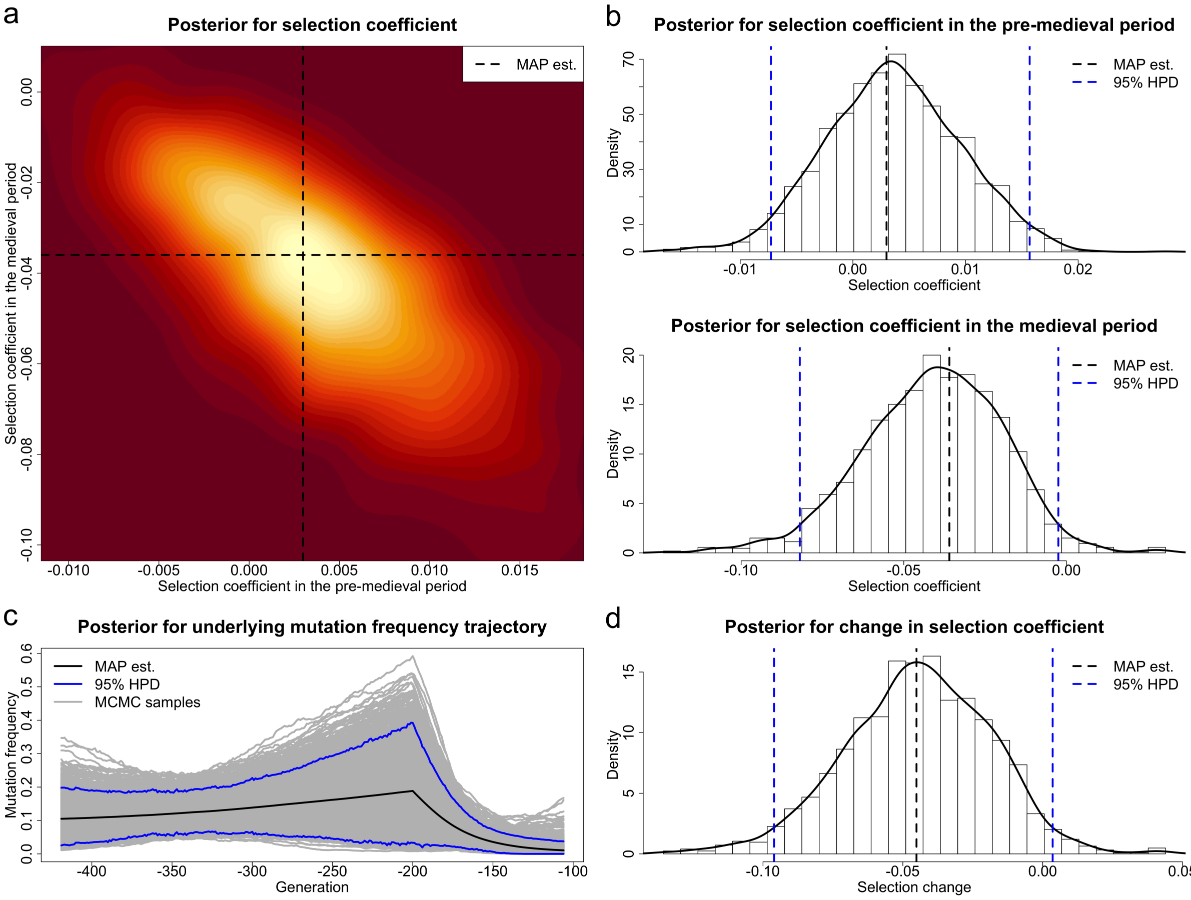


Figure S8: Posteriors for the selection coefficients of the *KIT13* mutation in the pre-medieval and medieval period and the underlying frequency trajectory of the *KIT13* mutation in the population produced through the PMMH-within-Gibbs procedure with the full forward-in-time simulation. The samples drawn in the period $[-514, -419)$ are excluded.

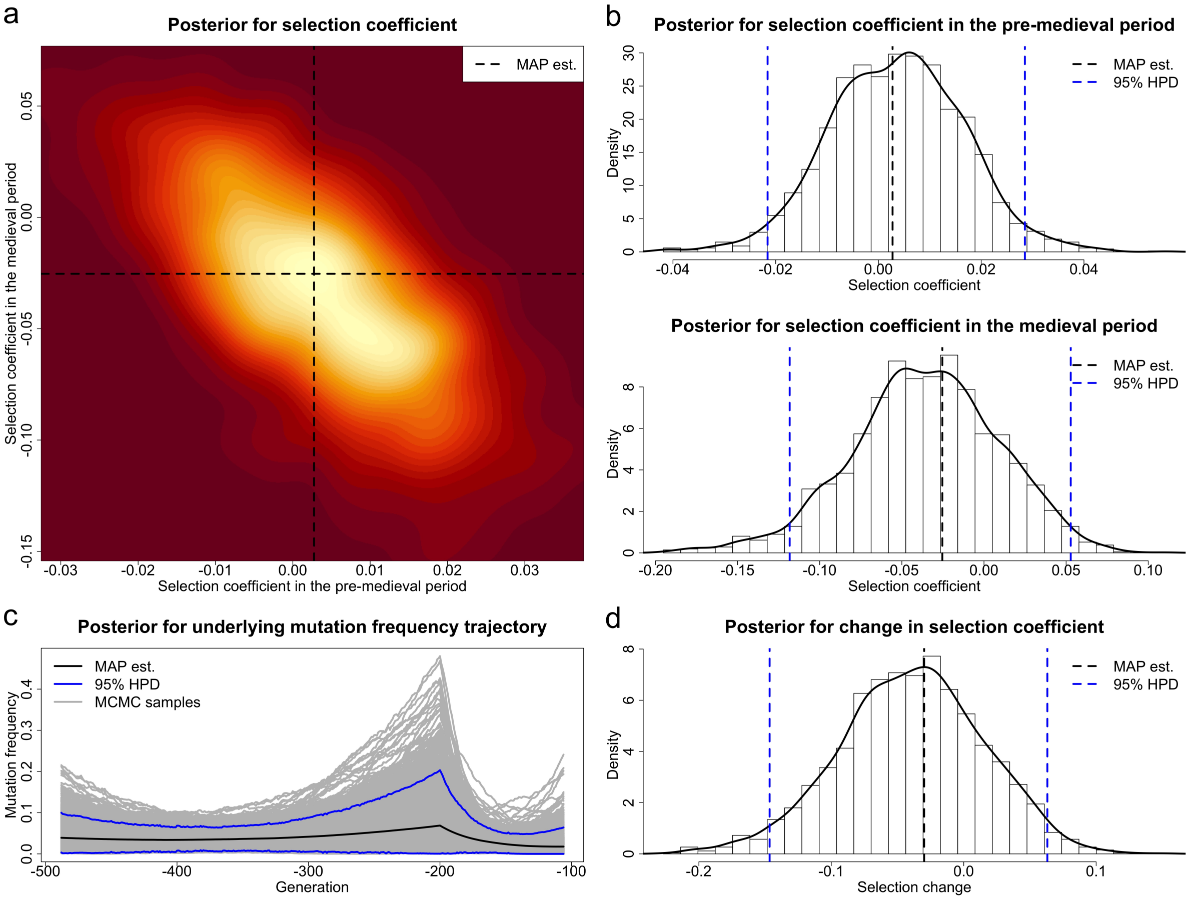


Figure S9: Posteriors for the selection coefficients of the *TRPM1* mutation in the pre-medieval and medieval period and the underlying frequency trajectory of the *TRPM1* mutation in the population produced through the PMMH-within-Gibbs procedure with the full forward-in-time simulation. The samples drawn in the period $[-514, -488)$ are excluded.

| Gene | Time period | Selection coefficient | MAP estimate | 95% HPD | $\Pr(s < 0)$ | $\Pr(s > 0)$ |
|---|---|---|---|---|---|---|
| | | $s^-$ | 0.00909 | [ 0.00115, 0.02085] | 0.0150 | 0.9850 |
| *MC1R* | [−488, −106] | $s^+$ | 0.00945 | [−0.01651, 0.03714] | 0.2725 | 0.7275 |
| | | $\Delta s$ | −0.00023 | [−0.03716, 0.03268] | 0.5345 | 0.4655 |
| | | $s^-$ | 0.00299 | [−0.00728, 0.01570] | 0.2750 | 0.7250 |
| *KIT13* | [−419, −106] | $s^+$ | −0.03599 | [−0.08197, −0.00248] | 0.9835 | 0.0165 |
| | | $\Delta s$ | −0.04512 | [−0.09605, 0.00357] | 0.9695 | 0.0305 |
| | | $s^-$ | 0.00275 | [−0.02158, 0.02850] | 0.3975 | 0.6025 |
| *TRPM1* | [−488, −106] | $s^+$ | −0.02535 | [−0.11828, 0.05281] | 0.7885 | 0.2115 |
| | | $\Delta s$ | −0.02990 | [−0.14657, 0.06316] | 0.7655 | 0.2345 |

Table S10: Estimates of the selection coefficients and their changes with their 95% HPD intervals, as well as posterior probabilities for negative selection/change and positive selection/change, for *MC1R*, *KIT13* and *TRPM1*, corresponding to Figures S7–S9, produced through the PMMH-within-Gibbs procedure with the full forward-in-time simulation. The time period is measured in generations (8 years per generation) and offset so that 2000 AD is 0.

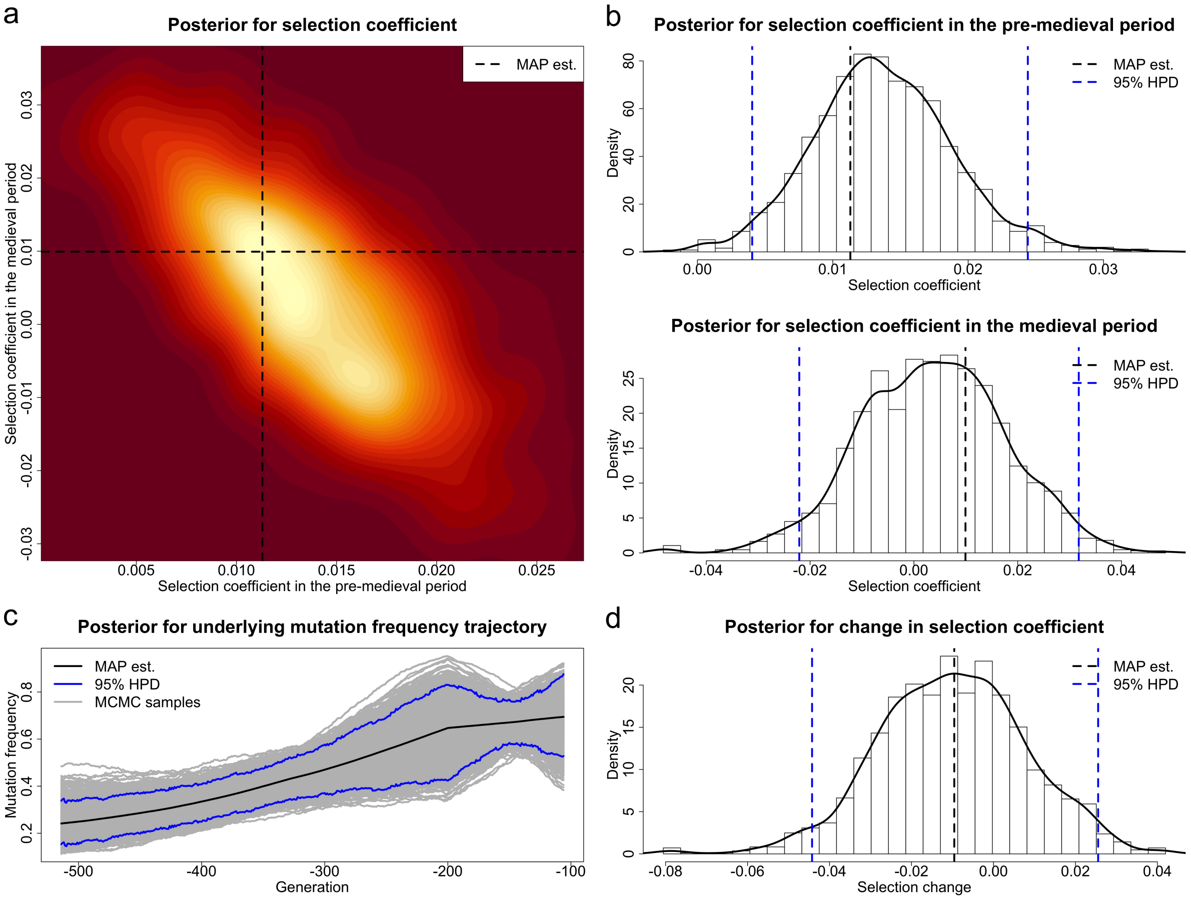


Figure S10: Posteriors for the selection coefficients of the *MC1R* mutation in the pre-medieval and medieval period and the underlying frequency trajectory of the *MC1R* mutation in the population produced through the PMMH-within-Gibbs procedure with the full backward-in-time simulation.

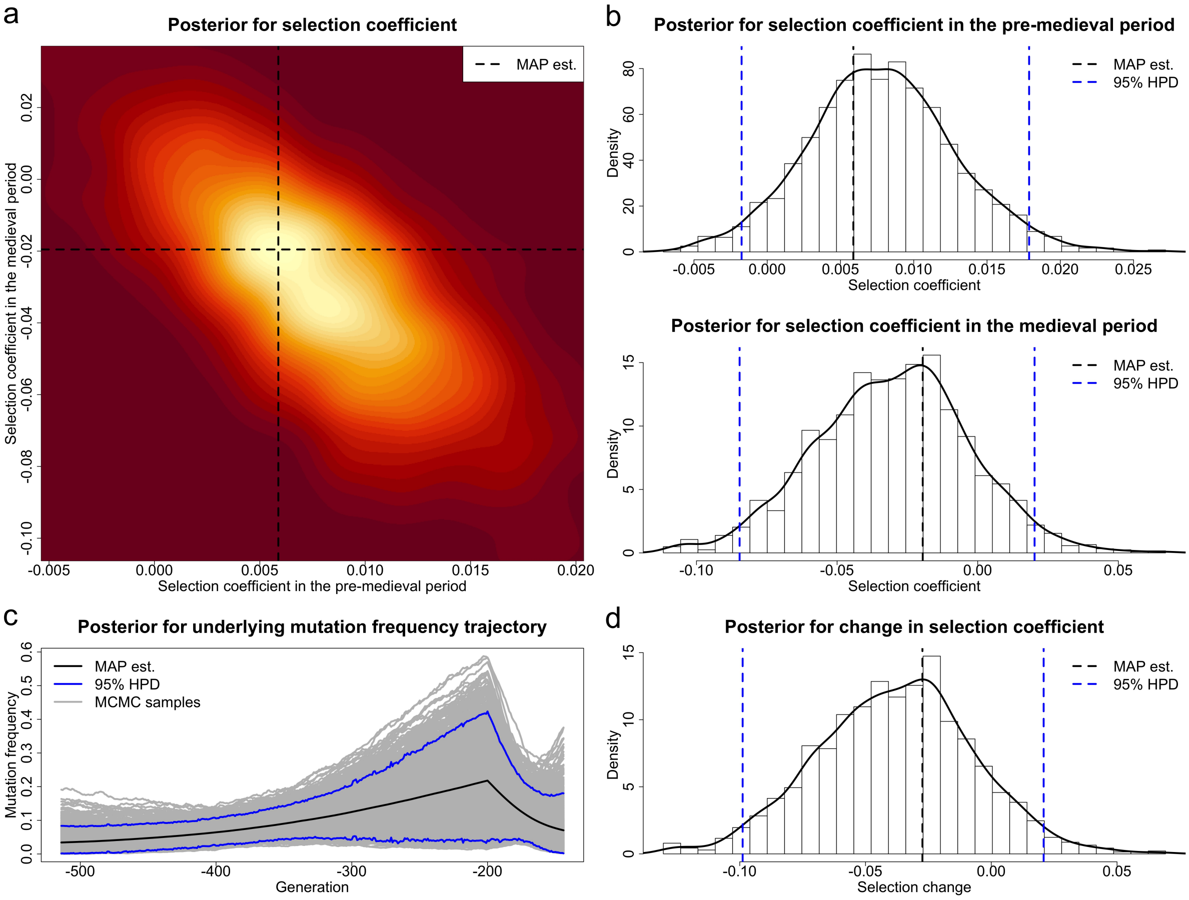


Figure S11: Posteriors for the selection coefficients of the *KIT13* mutation in the pre-medieval and medieval period and the underlying frequency trajectory of the *KIT13* mutation in the population produced through the PMMH-within-Gibbs procedure with the full backward-in-time simulation. The samples drawn in the period $(-144, -106]$ are excluded.

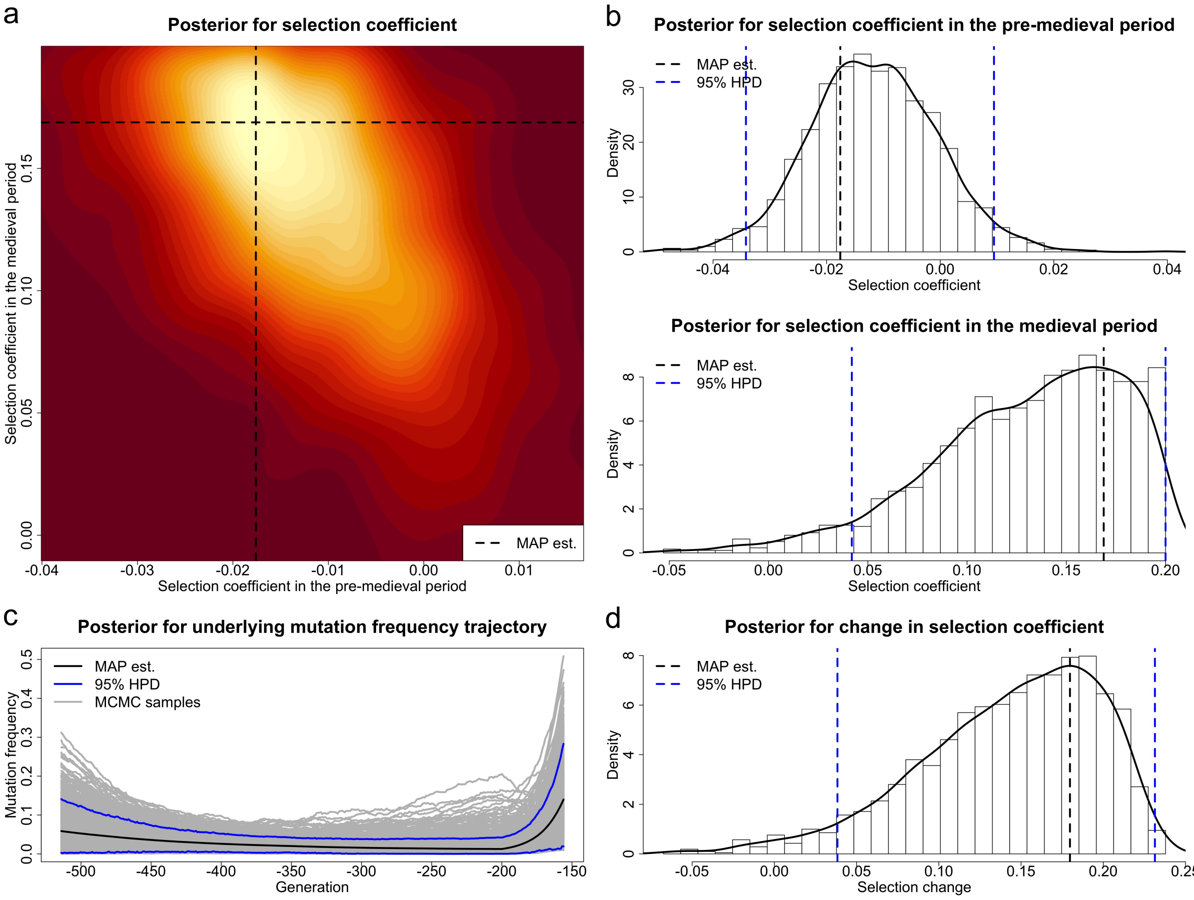


Figure S12: Posteriors for the selection coefficients of the *TRPM1* mutation in the pre-medieval and medieval period and the underlying frequency trajectory of the *TRPM1* mutation in the population produced through the PMMH-within-Gibbs procedure with the full backward-in-time simulation. The samples drawn in the period $(-156, -106]$ are excluded.

| Gene | Time period | Selection coefficient | MAP estimate | 95% HPD | $\Pr(s < 0)$ | $\Pr(s > 0)$ |
|---|---|---|---|---|---|---|
| *MC1R* | [−514, −106] | $s^-$ | 0.01130 | [ 0.00405, 0.02441] | 0.0015 | 0.9985 |
| | | $s^+$ | 0.00993 | [−0.02208, 0.03176] | 0.3935 | 0.6065 |
| | | $\Delta s$ | −0.00957 | [−0.04426, 0.02554] | 0.7115 | 0.2885 |
| *KIT13* | [−514, −144] | $s^-$ | 0.00588 | [−0.00175, 0.01787] | 0.0580 | 0.9420 |
| | | $s^+$ | −0.01955 | [−0.08466, 0.02022] | 0.8760 | 0.1240 |
| | | $\Delta s$ | −0.02737 | [−0.09885, 0.02082] | 0.8990 | 0.1010 |
| *TRPM1* | [−514, −156] | $s^-$ | −0.01759 | [−0.03420, 0.00947] | 0.8590 | 0.1410 |
| | | $s^+$ | 0.16883 | [ 0.04195, 0.19998] | 0.0120 | 0.9880 |
| | | $\Delta s$ | 0.17975 | [ 0.03836, 0.23141] | 0.0155 | 0.9845 |

Table S11: Estimates of the selection coefficients and their changes with their 95% HPD intervals, as well as posterior probabilities for negative selection/change and positive selection/change, for *MC1R*, *KIT13* and *TRPM1*, corresponding to Figures S10–S12, produced through the PMMH-within-Gibbs procedure with the full backward-in-time simulation. The time period is measured in generations (8 years per generation) and offset so that 2000 AD is 0.
